# We get by with a little help from our friends: shared adaptive variation provides a bridge to novel ecological specialists during adaptive radiation

**DOI:** 10.1101/2021.07.01.450755

**Authors:** Emilie J. Richards, Christopher H. Martin

## Abstract

Adaptive radiations involve astounding bursts of phenotypic, ecological, and species diversity. However, the microevolutionary processes that underlie the origins of these bursts are still poorly understood. We report the discovery of an intermediate ‘wide-mouth’ scale-eating ecomorph in a sympatric radiation of *Cyprinodon* pupfishes, illuminating the transition from a widespread algae-eating generalist to a novel microendemic scale-eating specialist. We first show that this ecomorph occurs in sympatry with generalist *C. variegatus* and scale-eating specialist *C. desquamator* on San Salvador Island, Bahamas, but is genetically differentiated, morphologically distinct, and often consumes scales. We then compared the timing of selective sweeps on shared and unique adaptive variants in trophic specialists to characterize their adaptive walk. Shared adaptive regions swept first in both the specialist *desquamator* and the intermediate ‘wide-mouth’ ecomorph, followed by unique sweeps of introgressed variation in ‘wide-mouth’ and de novo variation in *desquamator*. The two scale-eating populations additionally shared 9% of their hard selective sweeps with molluscivores *C. brontotheroides*, despite no single common ancestor among specialists. Our work provides a new microevolutionary framework for investigating how major ecological transitions occur and illustrates how both shared and unique genetic variation can provide a bridge for multiple species to access novel ecological niches.

## Introduction

Rapid bursts of diversification and repeated bouts of speciation like those seen in adaptive radiations contradict current mechanistic speciation models that predict diversification should slow with time as available niche space becomes increasingly subdivided and disruptive selection becomes weaker with each recurrent speciation event (e.g. [1–3]). Diversification on complex adaptive landscapes with multiple empty fitness peaks corresponding to different niches provides an alternative mechanism to niche subdivision [4–6]. However, these landscapes present a new problem to our mechanistic understanding of adaptive radiations: How do populations manage to escape local optima, cross fitness valleys, and access new fitness peaks [7–10]? Colonizing new fitness peaks on the adaptive landscape presents challenges because it requires transitions in behaviors, morphological traits, or a combination of the two that allow organisms to adapt to new ecological niches [11]. Spectacular ecological transitions do often occur during adaptive radiation, such as blood-drinking [12] or plant carnivory [13, 14], yet it is still poorly understood how such seemingly discontinuous transitions occur.

Recent conceptual frameworks for understanding adaptation to novel fitness peaks suggest that these major ecological transitions likely occur in stages of potentiation, actualization and refinement [15, 16]. The initial emergence of a novel trait likely requires further refinement to become successfully incorporated into the functional ecology of an organism. Several experimental lab studies suggest that novel ecological transitions are highly contingent on accruing a series of mutations that incrementally refine adaptations to colonize new fitness peaks [16, 17]. This idea that genetic background is important in setting the stage for adaptation also underlies many hypotheses for adaptive radiation, such as the hybrid swarm and syngameon hypotheses – in which radiations are driven by acquiring novel combinations of alleles through the exchange of genetic variation either from distinct lineages outside the radiation or within the radiation itself [18]. However, we are only just beginning to explore how gene flow and shared genetic variation gives recipient lineages access to new fitness peaks in the wild and generates adaptive radiations [6].

An adaptive radiation of trophic specialist pupfishes on San Salvador Island (SSI) in the Bahamas is an excellent system for understanding how the rapid evolution of major ecological transitions occurs in nature. This radiation contains a widespread generalist pupfish species (*Cyprinodon variegatus*) that occurs in sympatry with two previously described trophic specialists that are endemic to the hypersaline lakes on the island: a molluscivore (*C. bronotheroides*) with a novel nasal protrusion which is an oral-sheller of gastropods [19] and a scale-eating specialist (*C. desquamator*) with two-fold larger oral jaws [20]. The evolutionary novelties in this system originated recently; the lakes on SSI were dry during the last glacial maximum 6-20 kya years ago [21, 22]. Intriguingly, we recently discovered a fourth species of pupfish living in sympatry with the two specialists and generalist on SSI [23]. This species exhibits intermediate jaw morphology between *C. desquamator* and *C. variegatus* (figure 1). Here we refer to this new ecomorph as the ‘wide-mouth’ because its mouth is wider than any other species in the radiation. The multi-peak fitness landscape driving this radiation suggests that *C. desquamator* is isolated by a large fitness valley from *C. variegatus* and *C. brontotheroides* [9] and this intermediate ‘wide-mouth’ may provide clues about the microevolutionary processes underlying how the observed novel fitness peaks are traversed in the wild.

**Figure 1.**
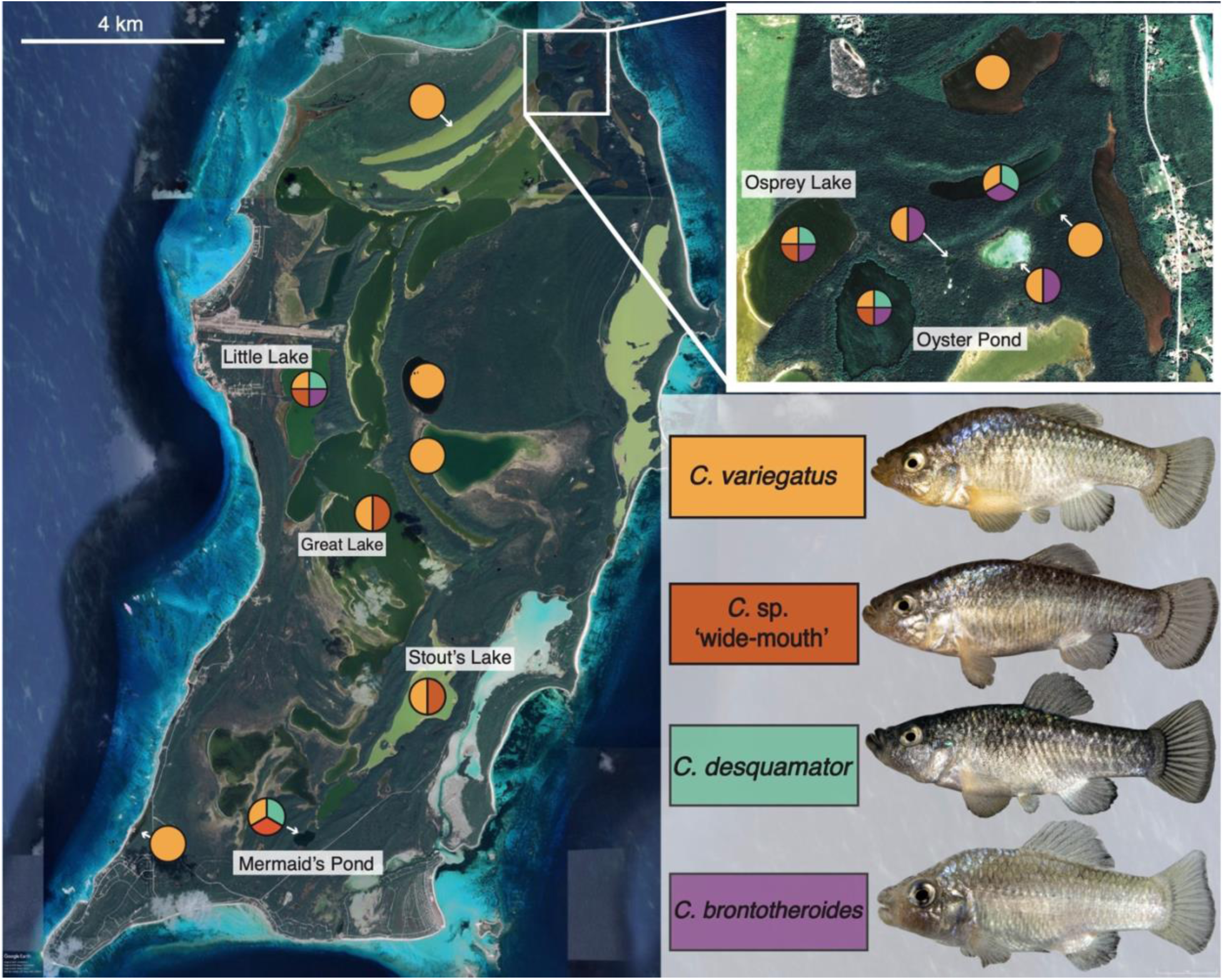
The San Salvador Island radiation of pupfish. Pie charts indicate the presence of sympatric *Cyprinodon* species in each lake and are color-coded with representative pictures of generalist *C. variegatus* (gold), recently discovered *C.* sp. ‘wide-mouth’ ecomorph (red-orange) with intermediate jaws, scale-eater *C. desquamator* (teal) with the largest oral jaws, and molluscivore *C. brontotheroides* (purple) with characteristic nasal protrusion. Labeled lakes contain all known *C.* sp. ‘wide-mouth’ populations sampled for this study. Satellite image from Google Earth.

Here we first investigate the position of the ‘wide-mouth’ on the ecological spectrum from generalist to scale-eating specialist using a combination of morphological, behavioral, dietary, and genomic data. We then estimated the demographic history of the ‘wide-mouth’ and explored the spatial origins and timing of selection on shared and unique genetic variation involved in adaptation to scale-eating to better understand this ecological transition. Our results suggest that while intermediate in jaw length known to be relevant for the highly specialized scale-eater *C. desquamator*, *C sp*. ‘wide-mouth’ demonstrates transgressive morphology and a distinct genetic background. Our investigation of the timing of selection and genetic origins of the adaptive alleles shared and unique between the two scale-eating species indicates divergent adaptive walks that are highly dependent on their genetic background. Despite shared origins, access to unique genetic variation in each of the two scale-eating sister species likely resulted in distinct adaptive walks and ultimately contributed to the diversity of ecological specialists observed in this radiation.

## Methods

### Ecological and morphological characterization of ‘wide-mouth’ scale-eater

*C. variegatus, C. desquamator,* and *C.* sp. ‘wide-mouth’ individuals from 3 lake populations (Osprey Lake, Great Lake, and Oyster Pond) in which we had sufficient specimens (n=84; *C. brontotheroides* not shown) were measured for 9 external morphological traits using digital calipers. Traits were selected for specific connections to foraging performance which differed across the three species in a previous study [9]. We also characterized diet for *C. variegatus, C. desquamator,* and ‘wide-mouth’ in Osprey Lake from stomach content analyses (*n*=10 per species) and stable isotope analyses of muscle tissue from wild-collected samples (*n*=75). Dietary overlap was characterized by comparison of population mean scale count from gut contents using ANOVA and ellipse areas and bivariate means on isotope biplots using SIBER [24]. See supplemental methods for more details on sample sizes and analyses.

### Genomic library preparation and variant filtration

To explore the evolutionary history of *C.* sp. ‘wide-mouth’, we sequenced whole genomes of 24 individuals following protocols used in a previous study [25] that included genomes from *C. variegatus, C. desquamator*, and *C. brontotheroides*. Our final genetic dataset after filtering contained 6.4 million variants across 110 individuals from the four species (7.9x median coverage). See supplemental methods for the full sequencing and genotyping protocol.

### Genomic origins of the ‘wide-mouth’ scale-eater

We first tested whether these *C*. sp. ‘wide-mouth’ individuals represented recent (e.g. F1/F2) hybrids of *C. variegatus* and *C. desquamator* in the wild using principal component and ADMIXTURE analyses to look for the genome-wide pattern expected in PCAs when recent hybrids between two populations are included. We also used formal tests for introgression and admixed populations, *f_3_* and *f_4_*-statistics [26], to assess whether ‘wide-mouth’ are the byproduct of recent admixture. Finally, we used *fastsimcoal2* (v2.6.0.3;[27]), a demographic modeling approach based on the folded minor allele frequency spectrum (mSFS), to discriminate among alternative evolutionary scenarios for the origin of ‘wide-mouth’ and estimated divergence times among all four species based on the best model fit from an AIC test (see supplementary methods for more detail).

### Characterization of unique and shared adaptive alleles among specialists

Across all four populations in Osprey Lake, we looked for regions that showed evidence of a hard selective sweep using SweeD (v.3.3.4;[28]). The composite likelihood ratio (CLR) for a hard selective sweep was calculated in 50-kb windows across scaffolds that were at least 100-kb in length (99 scaffolds; 85.6% of the genome). Significance thresholds were determined using CLR values from neutral sequences simulated under MSMC inferred demographic scenarios of historical effective population size changes (Supplemental methods; figure S1; table S3).

Next, we searched for candidate adaptive alleles associated with species divergence by overlapping selective sweep regions with regions of high genetic divergence based on fixed or nearly fixed SNPs between species. We chose to also look at regions with nearly fixed SNPs (F_st_ ≥ 0.95) to accommodate ongoing gene flow between these young species. F_st_ between the populations and species was calculated per variant site using –weir-pop-fist function in vcftools (v.0.1.15;[29]).

### Timing of selection on candidate adaptive alleles

We also determined the relative age of candidate adaptive alleles by generating estimates of coalescent times using starTMRCA (v0.6.1;[30]). For each candidate adaptive allele that was unique to the three specialists and the 16 shared alleles between *C. desquamator* and ‘wide-mouth’, a 1-Mb window surrounding the variant was extracted into separate vcfs for each species. These sets of variants were then analyzed in starTMRCA with a mutation rate of 1.56 x 10^-8^ substitutions per base pair (from Caribbean pupfishes [25]) and a recombination rate of 3.11x10^-8^ (from stickleback; [31]). Each analysis was run three times per focal adaptive allele and all runs were checked for convergence between and within runs. Most runs rapidly converged within the initial 6000 steps, but 5 runs did not converge after an additional 4000 steps and were discarded from further analysis. See supplementary methods for more details on timing analyses.

### Characterization of adaptive introgression adaptive alleles in *C.* sp ‘wide-mouth’

Lastly, we investigated the spatial origins of adaptive alleles shared and unique to the two scale-eating specialists by searching in our previous study spanning Caribbean-wide outgroup populations for these alleles [25]. Adaptive alleles were assigned as standing genetic variation if observed in any population outside SSI or de novo if they were only observed within populations on SSI. Additionally, we investigated signatures of introgression across the genome of *C.* sp ‘wide-mouth’ and *C. desquamator* to determine if they showed evidence of adaptive introgression from outgroup generalist populations as observed previously [25]. See supplementary methods for more details on introgression analyses.

## Results

### ‘Wide-mouth’ ecomorph is ecologically intermediate and morphologically distinct

We found that the ‘wide-mouth’ ecomorph is morphologically distinct from *C. desquamator* and *C. variegatus* across a suite of craniofacial traits (figure 2A-B). The lower jaw length of ‘wide-mouth’ was intermediate between *C. desquamator* and *C. variegatus* (figure 2C), while the buccal width and adductor mandibulae height were 8% larger in ‘wide-mouth’ than *C. desquamator* (figure 2E-F). These morphological differences were consistent across multiple lakes (figure S2). Small modifications in craniofacial morphology among these species have major impacts on scale-eating performance in this system by altering kinematic traits such as peak gap size which is partially controlled by the length of the lower jaw, jaw protrusion distance, and the angle of the lower jaw relative to the suspensorium [32].

**Figure 2.**
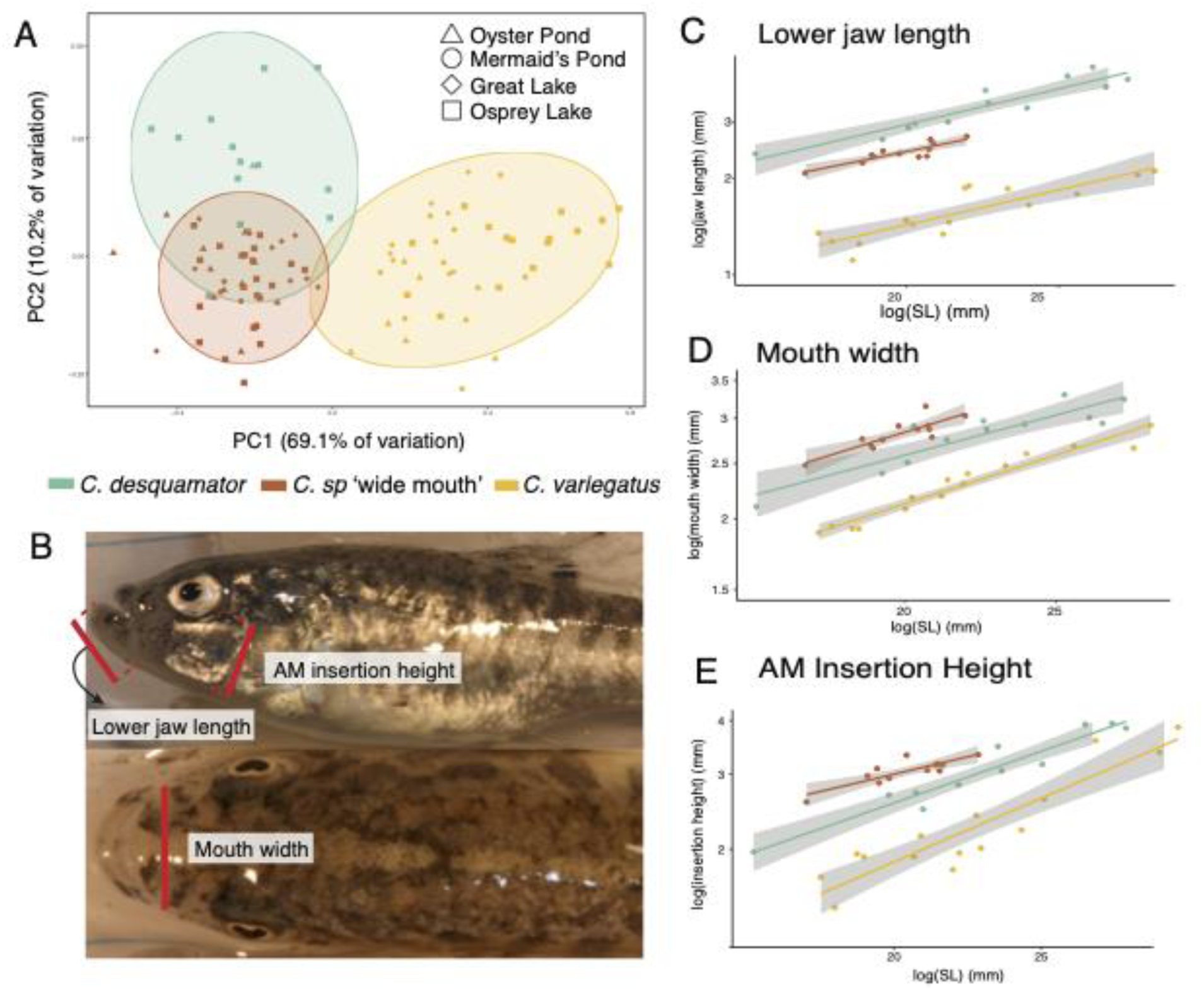
*C.* sp. ‘wide-mouth’ has distinct morphology within the San Salvador Island adaptive radiation. A) First two principal components of morphological diversity for 8 size-corrected traits and 95% confidence ellipses by species (*C. variegatus* : gold; *C.* sp. ‘wide-mouth’: red-orange; *C. desquamator*: teal; *C. brontotheriodes* not shown). PC1 is mainly described by lower jaw length and PC2 by adductor mandibulae insertion height, buccal width, and neurocranium width. B) Depictions of the three external measurements that best distinguished *C.* sp. ‘wide-mouth’ from both *C. desquamator* and *C. variegatus*, measured using digital calipers. C-E) The relationship between standard length (mm) of individuals and their C) lower jaw length, D) buccal cavity width, and E) adductor mandibulae insertion height (AM insertion) across individuals of the three species in Osprey Lake. 95% confidence bands for linear models in gray.

*C. sp* ‘wide-mouth’ also did not show morphological divergence comparable to that observed in the molluscivore *C. brontotheroides.* The molluscivore specialist presents an opposing axis of morphological divergence to the scale-eating specialists, with shorter oral jaw length and larger eye diameter than even the generalist *C. variegatus*, in addition to a novel nasal protrusion of the maxilla not observed in any other Cyprinodontidae species [33].

Morphological traits were heritable in a common garden laboratory environment after one generation: lab-reared *C.* sp. ‘wide-mouth’ displayed significantly larger buccal width than *C. desquamator* (t-test; *P* = 0.003) and maintained their characteristic intermediate jaw lengths (ANOVA; *P* = 0.03, figure S3). There was also some evidence of phenotypic plasticity in both *C. desquamator* and *C. sp* ‘wide-mouth’ compared to wild individuals likely caused by the common lab diet. See supplementary results for more details.

### ‘Wide-mouth’ occupies a distinct intermediate scale-eating niche

We found that ‘wide-mouth’ ingested scales, but at a significantly lower frequency than C. *desquamator* (Wilcoxon Rank Sum test, *P* = 0.004; figure 3A). We did not detect any scales in *C. variegatus* guts (figure 3A). Detritus made up the rest of the *C. sp.* ‘wide-mouth’ and *C. desquamator* diets and was the dominant component of *C. variegatus* gut contents, except for a single individual with one mollusc shell. A previous study that characterized contents of *C. variegatus*, *C. brontotheroides*, and *C. desquamator* populations across several ponds also found detritus to be the dominant component of each species’ diet (49-71%) and nearly zero scales in the gut contents of *C. variegatus* and *C. brontotheroides* [33].

**Figure 3.**
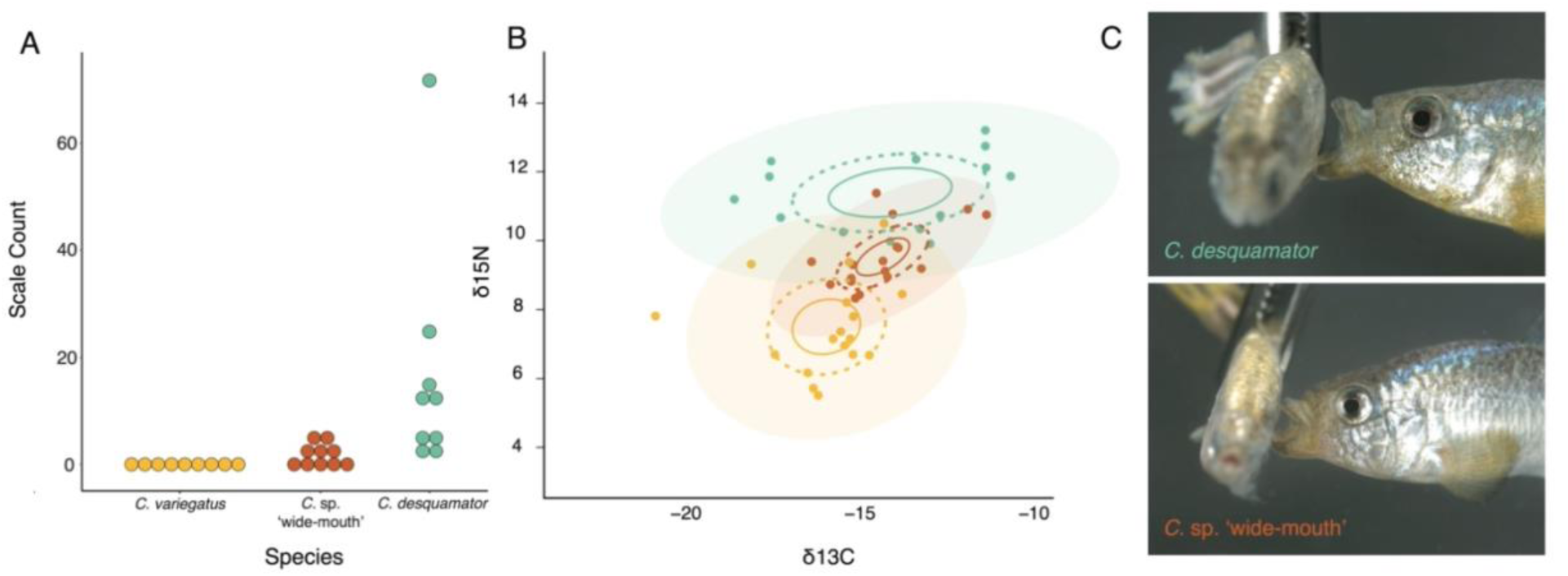
*C.* sp. ‘wide-mouth’ ingests scales. A) Scale counts from gut content analysis of the hindgut of Osprey pupfish populations (10 individuals per species). B) Relative trophic position (δ15N stable isotope ratio) and dietary carbon source (δ13C stable isotope ratio) with 95% confidence ellipses for generalist and scale-eating species. Solid lines represent 95% confidence intervals around bivariate mean, dotted lines represent standard ellipse areas corrected for sample size (contain 40% of data; SEAc), shaded circles represent ellipse area that contain 95% of the data calculated using the R package SIBER. C) Still images of scale-eating strikes in *C. desquamator* and *C.* sp. ‘wide-mouth’ filmed at 1100 fps on a Phantom VEO 440S camera.

The intermediate scale-eating dietary niche of the wide-mouth ecomorph is complemented by our stable isotope analyses, which provide long-term snapshots of the carbon sources and relative trophic levels in these species. Osprey Lake individuals collected on the same day from the same site differed in δ15N levels across species (ANOVA, *P* = 4.55x10^-6^; figure 3B and S4); ‘wide-mouth’ δ15N was intermediate between *C. variegatus* and *C. desquamator* (Tukey HSD; *P*= 1.34 x10^-5^ & 1.11 x10^-4^ respectively), supporting its intermediate trophic position. SIBER analyses of trophic niche position indicate distinct positioning of the wide-mouth ecomorph based on the lack of extensive overlap in niche space measured by standard ellipse areas and bivariate means with 95% confidence intervals of isotope values among the species (figure 3B).

### *C*. sp. ‘wide-mouth’ did not result from hybridization between *C. variegatus* and *C. desquamator*

Several lines of genomic evidence from PCA, ADMIXTURE, and *f-statistic* analyses support the ‘wide-mouth’ ecomorph as a genetically distinct species rather than a recent hybrid between *C. desquamator* and *C. variegatus* on SSI as their intermediate jaw morphology might suggest (figure 4A-C & S5-6; see Supplementary material for more details). Demographic modeling of divergence and gene flow on SSI places *C. sp* ‘wide-mouth’ as sister to *C. desquamator*, supporting previous phylogenetic inference [34]. In the best supported model of 28 demographic models tested (table S2), ‘wide-mouth’ and *C. desquamator* diverged 11,658 years ago (95 CI: 8,257-20,113 years; figure S7B; table S2) with ongoing gene flow. Additionally, the amount of genetic divergence between populations indicates that *C. desquamator* and *C.* sp. ‘wide-mouth’ are more genetically diverged from each other than to the generalist *C. variegatus* (e.g. *F_st_* in figure 4C).

**Figure 4.**
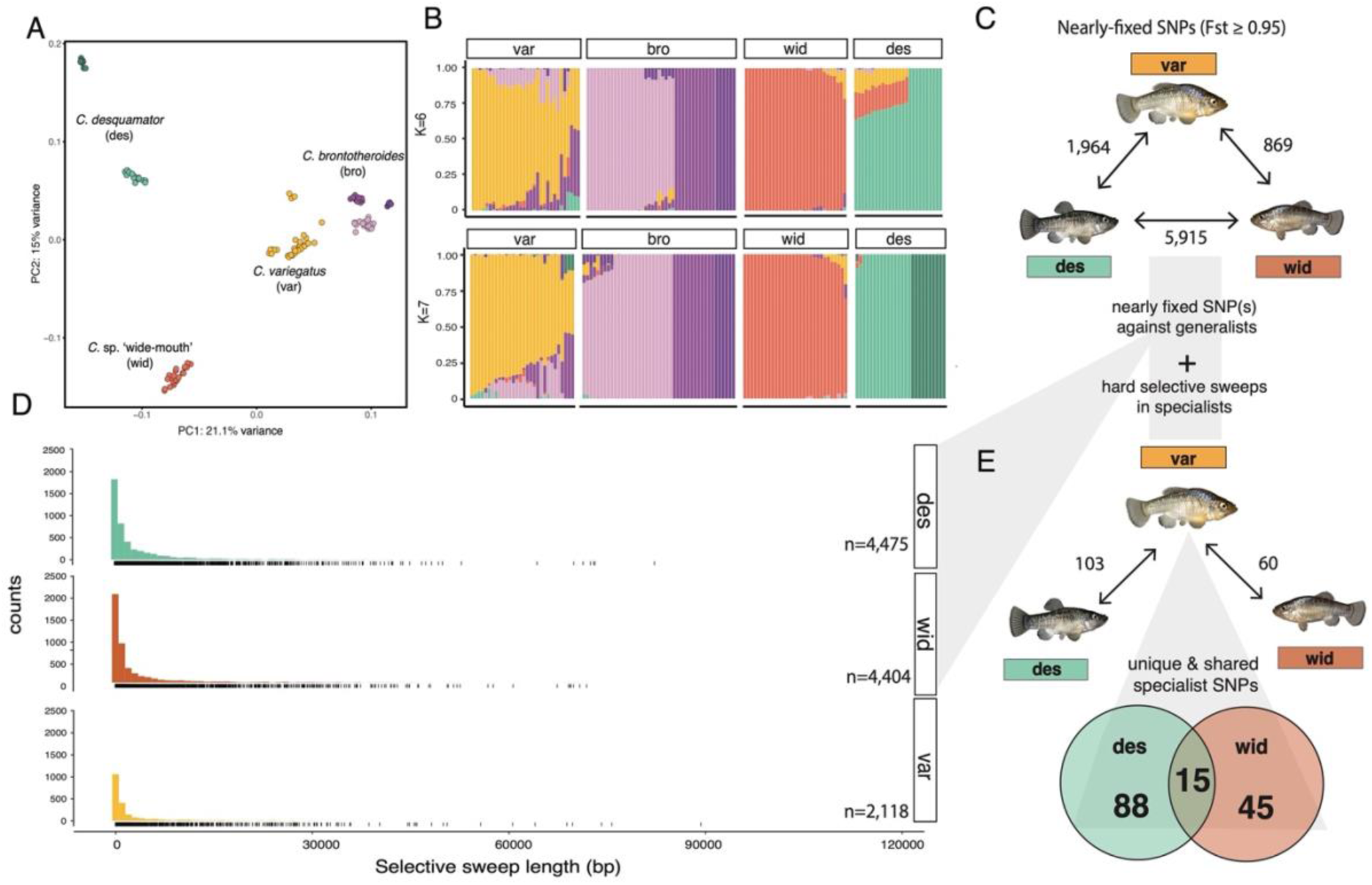
Patterns of selection and genetic divergence in specialist genomes. A) Principal components analysis of the four focal groups on San Salvador Island based on an LD-pruned subset of genetic variants (78,840 SNPs). B) Ancestry proportions across individuals of the four focal groups. Proportions were inferred from ADMIXTURE analyses with 2 values of K with the highest likelihood on the same LD-pruned dataset in A. C) Selective sweep length distributions across generalist and scale-eating species. Rug plot below each histogram represents the counts of selective sweeps in different length bins. D) The total number of fixed or nearly-fixed SNPs (*F_st_*≥ 0.95) between each group in Osprey Pond. E) The number of adaptive alleles (fixed or nearly-fixed SNPS [*F_st_*≥ 0.95] relative to *C. variegatus* and under selection in each population of specialists in Osprey Lake. Venn diagram highlights those adaptive alleles that are unique to each specialist and shared with other specialists.

### *C.* sp ‘wide-mouth’ is comprised of both shared and unique adaptive alleles

Next we looked at regions of the genome in both *C. desquamator* and *C*. sp. wide-mouth that showed strong evidence of hard selective sweeps. We found 6 shared hard selective sweeps in both species containing a total of 15 SNPS that were fixed or nearly-fixed compared to the sympatric generalist *C. variegatus* (figure 4E, 5): 10 SNPs were in unannotated regions, two were in the introns of the gene *daam2,* and three were in putative regulatory regions (with 20-kb) of the genes *usp50, atp8a1,* and *znf214* (one variant each). Shared adaptive alleles in the gene *daam2,* a wnt signaling regulator, are intriguing because knockdown of this gene causes abnormal snout morphology in mice [35] and abnormal cranial and skeletal development in zebrafish [36].

**Figure 5.**
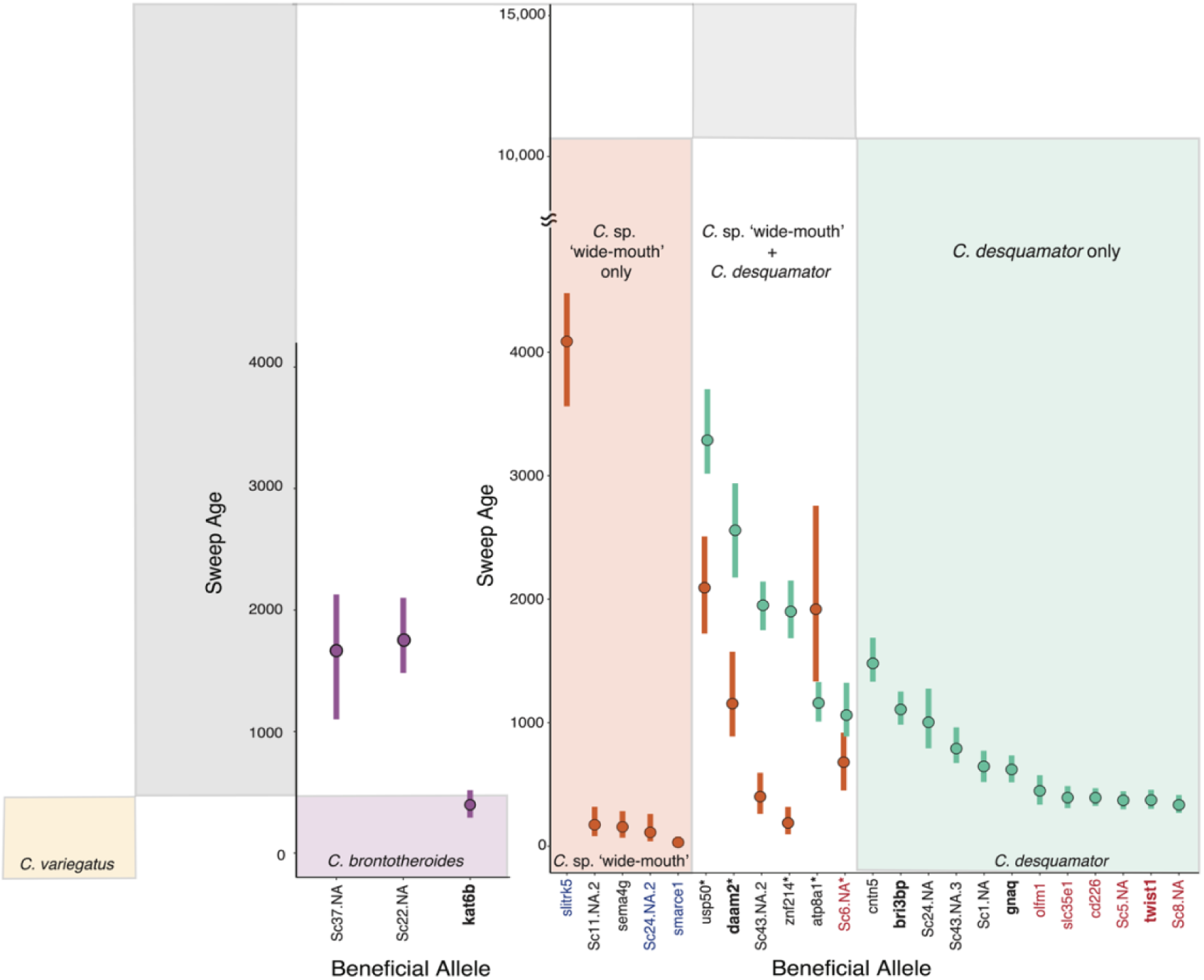
Timing of selection on adaptive alleles in trophic specialists nested within the demographic history of the radiation. The mean and 95% HPD estimates for the timing of selection on sets of fixed or nearly fixed SNPs (named by the gene they are in or within 20-kb of) for the three specialist populations found in sympatry in Osprey Lake (sweeps in *C. variegatus* not shown). The age of each beneficial allele is color coded by the species and the inferred demographic history is displayed in the background for comparison. Gene names highlighted in bold are associated with oral jaw size. Gene names are colored by source of genetic variation (de novo: red; introgressed with outgroup: blue; standing genetic variation: black). Gene names with asterisk indicate those inferred as introgressed between *C. desquamator* and *C.* sp ‘wide-mouth’.

We also found unique sets of adaptive alleles in *C*. sp ‘wide-mouth’ and *C. desquamator* (figure 4E;figure 5). None of the adaptive alleles unique to *C*. sp. ‘wide-mouth’ were in or near genes annotated for craniofacial phenotypes in model organisms, despite its distinctive craniofacial morphologies. In *C. desquamator,* three of 12 unique adaptive alleles were in or near genes associated with or known to affect craniofacial phenotypes: a *de novo* non-synonymous coding substitution in the gene *twist1*, several putative regulatory variants near the gene *gnaq,* and 8 variants in or near the gene *bri3bp*, which is located inside a QTL region for cranial height in pupfish [37]*. C. brontotheriodes* also contained at least one unique candidate craniofacial adaptive allele: a non-synonymous coding substitution in the gene *kat6b* (figure 5), which is associated with abnormal craniofacial morphologies, including shorter mandibles, in mice [38]. This pattern of unique alleles relevant to craniofacial phenotypes in specialists *C. brontotheriodes* and *C. desquamator*, but not *C.* sp ‘wide-mouth’, holds even if we lower the threshold to the top 1 percentile of *F_st_* outliers between specialists and generalist (see supplemental results; figure S10).

#### The origins of adaptive alleles in C. sp ‘wide-mouth’ and desquamator

The adaptive alleles shared by *C. desquamator* and *C.* sp. ‘wide-mouth’ occurred as low frequency standing genetic variation in the Caribbean, with the exception of a single de novo allele on SSI located in an unannotated region on scaffold 6 (figure 5). The adaptive alleles unique to *C. desquamator* and *C*. sp ‘wide-mouth’ also predominantly came from standing genetic variation (84% and 81%, respectively). 14% of adaptive alleles unique to *C. desquamator* were de novo mutations to SSI and 2% occurred in candidate introgression regions (table S7). We found the opposite in *C. sp* ‘wide-mouth’: only 4% of their unique adaptive alleles were de novo mutations whereas 15% occurred in candidate introgression regions (table S8). This adaptive introgression was detected for generalist populations sampled from North Carolina and Laguna Bavaro in the Dominican Republic (table S8; figure S11). Using the Relative Node Depth (RND) statistic, we also discovered that 5 of the 6 shared adaptive alleles (all except for the unannotated region on scaffold 43;table S6) appear introgressed between *C. desquamator* and *C.* sp. ‘wide-mouth’, suggesting a substantial contribution of introgression to the adaptive alleles observed in *C.* sp. ‘wide-mouth’.

### Timing of selection on adaptive alleles reveals features of the adaptive walk to scale-eating

Selective sweeps occurred much more recently in both populations than their inferred divergence times (figure 5). Intriguingly, selection on 4 of the 6 adaptive alleles occurred significantly earlier in *C. desquamator* than ‘wide-mouth’. Only a single adaptive allele had an older median age estimate in ‘wide-mouth’ than *C. desquamator*, although the 95% HPD intervals overlapped between the species (figure 5). Additionally, overall we found a significant difference in timing of selection between shared and unique adaptive alleles in the two scale-eater populations (ANOVA *P*-value = 0.00478). In *C. desquamator*, shared adaptive alleles swept before any unique adaptive alleles (Tukey HSD *P-*value = 0.003217; figure 5). For the ‘wide-mouth’, shared adaptive alleles with *C. desquamator* also generally swept before those unique to the species, despite these unique alleles being standing and introgressed variation from the Caribbean (figure 5). However, this difference was not significant due to one unique adaptive allele (*slitrk5)* that swept first (figure 5; ANOVA, Tukey HSD; *P* = 0.8367). This adaptive allele resides in a region that appears to be introgressed with the Laguna Bavaro generalist population in the Dominican Republic where this allele also show signs of a hard selective sweep [25]. The older age estimate of this sweep in *C*. sp ‘wide-mouth’ might be due to older shared selection for the alleles in other Caribbean populations before introgression with *C*. sp ‘wide-mouth’. All other introgressed adaptive alleles in *C.* sp. ‘wide-mouth’ swept more recently than shared sweeps with *desquamator*, including the shared de novo allele, and were not under selection in outgroup generalist populations.

Intriguingly, all but one of the de novo adaptive alleles in *desquamator* swept at the same time in the recent past (figure 5). Only one of these adaptive alleles in *olfm1* region has 95% HPD age range that overlaps with the next oldest selective sweep on standing genetic variation (*gnaq*; figure 5), suggesting a discrete stage of selection on new mutations in *C. desquamator*.

### Shared signatures of selection across the three specialists in the radiation

Lastly, we compared the genetic divergence and selection patterns observed in the two scale-eating specialists to the divergent molluscivores specialist *C. brontotheroides* to investigate the extent of allele sharing among all three trophic specialists in this adaptive radiation. We found that no fixed or nearly-fixed alleles relative to the generalist *C. variegatus* were shared across all three specialists (figure S9-S10; supplementary results). However, we did find evidence of 44 shared selective sweeps across all three specialist populations that were not shared with *C. variegatus* populations (figure S12C). These shared regions were significantly enriched for genes annotated for metabolic processes (figure S12D), suggesting shared selection for metabolizing the more protein-rich diet across all three trophic specialists (also see [39]).

## Discussion

### Discovery of a new cryptic intermediate scale-eater highlights the power of reusing genetic variation to access novel niches

The hallmark of adaptive radiation is a rapid burst of diversification which is predicted by theory to slow down over time as niche subdivision increases [6]. An alternative possibility is that radiations can be self-propagating and that the diversity generated within the first stages of radiation helps beget further diversity [40]. This could happen through exploitation of new trophic levels created by new species or physical alterations of the environment by new species that may create additional opportunities for speciation (reviewed in [6, 41]). The diversity begets diversity hypothesis can also be visualized as the exploration of a complex multi-peaked fitness landscape; as species in the radiation colonize new peaks, this provides access to additional neighboring fitness peaks to fuel rapid radiation. Our discovery of a cryptic new scale-eating species through morphological, dietary, and genomic analyses revealed shared nearly-fixed or fixed adaptive alleles in both scale-eating species relative to the generalist *C. variegatus*. While *C.* sp. ‘wide-mouth’ is ecologically intermediate in its scale-eating behavior, our estimates of the relative timing of selective sweeps suggest that these shared alleles were first selected upon in the more specialized scale-eater *C. desquamator*, although unaccounted for demographic differences may also be contributing to differences in estimated timing between species.

Intriguingly, the shared adaptive alleles between *C. desquamator* and *C.* sp. ‘wide-mouth’ have potentially introgressed more recently rather than selected upon in their shared common ancestor. Five of the six regions surrounding these shared adaptive alleles showed patterns of high genetic similarity consistent with introgression (table S10). Alternatively, this genetic similarity may also be caused by strong background selection on shared ancestral variation. Effective population sizes are not drastically different between the two species and exon density is not in the upper tail of the genome wide-distribution (figure S1;table S10), two factors found in other studies where background selection tends to confound adaptive introgression inferences [42, 43]. However, we do not have extensive knowledge of recombination breakpoints in this non-model system to distinguish between strong background selection on shared ancestral variation and adaptive introgression scenarios for each candidate adaptative introgression region.

We also found strong signatures of introgression in *C.* sp. ‘wide-mouth’ genomes from outgroup generalist populations that were not shared with *C. desquamator* (figure S11; table S11). Craniofacial morphology is a major axis of diversification between trophic specialists in this system [44], yet *C.* sp. ‘wide-mouth’ appears to have little unique genetic variation relevant for craniofacial traits compared to the other two specialists (figure S10). Despite this, they do exhibit transgressive craniofacial phenotypes not seen in the other specialists. An intriguing implication of these findings is that hybridization may allow different species to share many of the same adaptive alleles to occupy distinct but similar niches, in line with the syngameon and ‘diversity begets diversity’ hypotheses of adaptive radiation [18, 40].

### An adaptive walk underlies the major ecological transition from generalist to scale-eating specialist

The foundational model of adaptation is that it proceeds in ‘adaptive walks’ towards fitness optima that involve the sequential fixation of adaptive alleles that move a population in the phenotypic direction of the local optimum [45]. The distinct timing of selection across different adaptive alleles in both *C. desquamator* and *C.* sp ‘wide-mouth’ suggests that the ecological transition from generalist to novel scale-eating specialist involved an adaptive walk in which selection on a beneficial allele was contingent on prior fixation of other adaptive alleles in each specialists’ genetic background (see supplemental materials for further discussion). This is best highlighted by the pattern observed in *C. desquamator* in which nearly all de novo mutations swept at the same time in a distinct selective stage from other adaptive variants rather than being selected upon as they originated (figure 5).

### The (un)predictability of adaptive walks to novel ecological niches

A recent study characterizing genotypic fitness landscapes underlying the transition from *C. variegatus* and *C. desquamator* revealed a rugged landscape with many local fitness peaks, likely due to epistatic interactions among alleles [46]. The staggered timing of selection on alleles lends support to this finding. Epistasis can reduce the number of adaptive walks selection will promote [47], and might explain why the same adaptive alleles were the first to undergo hard selective sweeps in both ‘wide-mouth’ and *desquamator*.

We also found evidence for shared selective sweeps across all three specialists in regions that are enriched for genes annotated for metabolic processes such as short chain fatty acid and propionate metabolism (figure S12D). The lack of fixed alleles in these regions relevant to dietary specialization suggests polygenic selection (see supplemental for more discussion). Subtle shifts of allele frequencies across the genome can lead to divergent genomic backgrounds that give populations access to different ecological niches (e.g. [48, 49]).

While both shared sweeps among all specialists and shared adaptive alleles among the two scale-eating species suggest constrained adaptive walks along overlapping genotypic pathways, we still see most selective sweeps are unique to each species in this radiation (figure 4; figure S11). Curiously, some adaptive standing genetic variation rose to high frequency in *C. desquamator* but did not similarly undergo selection in *C.* sp ‘wide-mouth’, despite its adaptation to a similar ecological niche and the presence of these alleles segregating at low frequency in the ‘wide-mouth’ population. This highlights the dual influence of epistatic interactions on adaptive walks in rugged landscapes – epistasis reduces number of available paths but increases the number of local fitness peaks populations can get stuck on [50]. Selection on the same adaptive alleles may have allowed both scale-eating species access to the same area of the fitness landscape but epistatic interactions with private mutations and introgressed variation in each lineage may have resulted in divergent paths to scale-eating, ultimately contributing to diverse evolutionary outcomes even from a shared starting point.

The use of adaptive alleles from distinct spatial sources, the distinct morphologies and divergent genomic backgrounds, and potential introgression of adaptive alleles from the more specialized scale-eater *C. desquamator* into *C.* sp ‘wide-mouth’ reveals a tangled path for novel ecological transitions in nature. The complex epistatic interactions at microevolutionary scales implicated in this study make it all the more fascinating that novel ecological transitions are a common macroevolutionary feature of adaptive radiation.

## Acknowledgements

We thank Priya Moorjani, Michelle St. John, Joseph McGirr, Jacquelyn Galvez, David Tian, Austin Patton, and Joseph Heras for helpful comments on the manuscript; the Gerace Research Centre and Troy Day for logistical support; the governments of the Bahamas for permission to collect and export samples. This research was funded by the National Science Foundation DEB CAREER grant #1749764, National Institutes of Health grant 5R01DE027052-02, the University of North Carolina at Chapel Hill, and the University of California, Berkeley to CHM.

## Author Contributions

Conceptualization: EJR,CHM; Data Collection: EJR,CHM; Statistical analyses: EJR; Resources: CHM; Visualization: EJR,CHM; Original draft: EJR; Revising: EJR, CHM

## Ethics Statement

Fishes were euthanized in an overdose of buffered MS-222 (Finquel, Inc.) following approved protocols from the University of North Carolina at Chapel Hill Animal Care and Use Committee (#18-061.0), and the University of California, Berkeley Animal Care and Use Committee (AUP-2015-01-7053) and preserved in 95-100% ethanol. All animals were collected and exported with 2017 and 2018 permits from the Bahamas Environmental Science and Technology commission and Ministry of Agriculture.

## Data Accessibility

All genomic data are archived at the National Center for Biotechnology Information BioProject Database (Accessions: PRJNA690558; PRJNA394148, PRJNA391309; and PRJNA305422). Ecological and morphological data are provided in the supplemental materials.

## Supplemental Text, Figures and Tables

### Methods

#### Sampling

*C.* sp. ‘wide-mouth’ individuals were collected on San Salvador Island in the Bahamas using hand and seine nets between 2017 and 2018. ‘Wide-mouth’ individuals were collected from 5 lakes in which taxa occurs in sympatry with just *C. variegatus* (Great Lake, and Stout’s Lake) or with generalist and *C. desquamator* (Oyster Pond, Osprey Lake, and Mermaid’s Pond,; Fig. 1). In Oyster Pond and Little Lake, *C. desquamator* is extremely rare (only a single Oyster Pond individual grouped genetically and morphologically with *C. desquamator*), so Osprey Lake appears to be the only lake where all three species and the recently discovered ‘wide-mouth’ coexist at appreciable frequencies in sympatry. ‘Wide-mouth’ individuals have also been collected in Little Lake in the past (Bruce Turner, pers. obs. 1990s).

Fishes were euthanized in an overdose of buffered MS-222 (Finquel, Inc.) following approved protocols from the University of North Carolina at Chapel Hill Animal Care and Use Committee (#18-061.0), and the University of California, Berkeley Animal Care and Use Committee (AUP-2015-01-7053) and preserved in 95-100% ethanol.

#### Ecological and morphological characterization of ‘wide-mouth’ scale-eater

Multiple individuals from 3 lake populations (Osprey Lake, Great Lake, and Oyster Pond) in which we had measurable specimens of the species (n=84) were measured for 9 traits using digital calipers. Traits were selected for specific connections to foraging performance [1, 2] and that differed across the three groups in a previous study when externally measured [3]. For each specimen we measured (1) standard length from the anterior tip of the premaxilla to the posterior margin on the midline of the caudal peduncle, (2) lower jaw length from the medial point of the dentary on the lower jaw to the distal point of rotation on the quadrate-articular joint, (3) the width of the buccal from the distance between the proximal sides of the buccal cavity in dorsal view, (4) the height of the adductor mandibulae insertion from the vertex between the vertical and horizontal arms of the preopercle to the dorsal margin of the hyomandibula, (5) the distance from the insertion of the first ray of the dorsal fin to the insertion of the first ray of the anal fin, (6) the height of the caudal peduncle from the dorsal and ventral procurrent rays on the caudal fin, (7) neurocranium width from the narrowest distance between the eyes from the dorsal view, (8) body width from just behind the posterior margins of each operculum in dorsal view, and (9) diameter of the orbit. Measurements were made by a single observer (EJR). Repeatability of measurements was assessed by remeasuring a random subset of 10 individuals per species twice. Measurements on the full dataset were not taken until repeatability was above 85% per trait (*r^2^* in a linear regression of both sets of measurements). To remove the effects of body size on trait size, we used the residuals from a linear regression of the log-transformed trait on log transformed standard length. We also compared these morphological measurements between lab-reared F2 and wild caught specimens to rule out strong plasticity in traits of interest.

For a subset of individuals collected from each population (*C. variegatus, C. desquamator,* and ‘wide-mouth’) in Osprey Lake, diet was characterized from stomach content analyses (*n*=10 each) and stable isotope analyses (*n*=75; 42 *C. variegatus*, 16 *C. desquamator*, 17 ‘wide-mouth’ individuals). These are two complementary approaches that reflect short-term and long-term snapshots of the dietary differences among these groups. For stomach content analysis, a single observer (EJR) dissected out a 1 cm portion of the gut from the posterior hindgut region and counted the number of scales found. For stable isotope analyses, muscle tissue samples were taken from the caudal peduncle region of each individual immediately after euthanasia in an overdose of MS-222. Muscle samples were individually labeled and dried at 60° C for at least 24 hours. Foil-wrapped tissue samples were then sent to the UC Davis Stable isotope facility and analyzed on a PDZ Europa ANCA-GSL elemental analyzer interfaced to a PDZ Europa 20-20 isotope ratio mass spectrometer to characterize the natural abundances of δ13C and δ15N among individuals. δ13C and δ15N levels were summarized across each of the three species to look at overlap among species by 40% and 95% Standard Area Ellipses and 95% confidence ellipses around bivariate means on isotope biplots (figure 3) of the data using R package SIBER [4]. Additionally we assessed whether there were significant differences in δ15N values between species through an ANOVA using aov() function from the base statistics package in R (v 4.1.0).

#### Genomic library preparation

We sequenced 24 ‘wide-mouth’ individuals following protocols used in a previous study [5] in which we sequenced genomes from *Cyprinodon variegatus*, *C. desquamator*, and *C. brontotheroides*. Raw reads were mapped from these 24 individuals to a de-novo assembly of *Cyprinodon brontotheroides* reference genome (v 1.0; total sequence length = 1,162,855,435 bp; number of scaffolds = 15,698, scaffold N50 = 32 Mbp) with bwa-mem (v 0.7.12;[6]). Duplicate reads were identified using MarkDuplicates and BAM indices were created using BuildBamIndex in the Picard software package (http://picard.sourceforge.net(v.2.0.1)). We followed the best practices guide recommended in the Genome Analysis Toolkit (v 3.5;[7]) to call and refine our single nucleotide polymorphism (SNP) variant dataset using the program HaplotypeCaller. Variants in ‘wide-mouth’ individuals were called jointly with the 202 individuals sequenced from a previous study [5]. We filtered SNPs based on the recommended hard filter criteria (i.e. QD < 2.0; FS > 60; MQRankSum < -12.5; ReadPosRankSum < -8;[7, 8]) because we lacked high-quality known variants for these non-model species. After selecting for only individuals from San Salvador Island, variants were additionally filtered to remove SNPs with a minor allele frequency below 0.05, genotype quality below 20, or containing more than 10% missing data across all individuals at the site using vcftools (v.0.1.15;[9]). Variants in poorly mapped regions were then removed using a mask file generated from the program SNPable (http://bit.ly/snpable; k-mer length =50, and ‘stringency’=0.5). Our final dataset after filtering contained 6.4 million variants with 7.9x median coverage per individual.

#### Testing for signatures of recent hybrids

We first tested whether these ‘wide-mouth’ individuals represented recent (e.g. F1/F2) hybrids of *C. variegatus* and *C. desquamator* in the wild. These two young species do reproduce with each other in the lab and rare hybrid spawning events have been observed in the wild (unpublished data ME St. John). As a first visual assessment, we conducted principal component analysis on *C. desquamator*, generalist, and ‘wide-mouth’ individuals to look for the genome-wide pattern expected in PCAs in which recent hybrids between two populations are included. To perform the PCA, the genetic dataset was first pruned for SNPs in strong linkage disequilibrium using the LD pruning function (--indep-pairwise 50 5 0.5) in plink (v1.9;[10]), leaving 2.6 million variants. We then ran a principal component analysis using the eigenvectors output by plink’s pca function (--pca). The first two principal components were plotted in R (R Core Team 2018 v3.5.0).

As a complementary assessment, we used an admixture model-based approach to estimate the proportion of shared ancestry among individuals in our dataset using ADMIXTURE (v.1.3.0)[11]. The number of populations (K) was decided upon using ADMIXTURE’s cross-validation method (--cv) across 1-10 population values of K. K = 4 was then chosen using the broken-stick method. Ancestry proportions estimated by ADMIXTURE were plotted in R for the K value with the highest likelihood and the two K values surrounding to explore whether the strong signatures of population structure in Crescent Pond *C. desquamator* individuals was masking hybridization signatures in any of the ‘wide-mouth’ populations.

Lastly, to discriminate among alternative evolutionary scenarios for the origin of ‘wide-mouth’ and to estimate divergence time among all four species, we used demographic modeling based on the folded minor allele frequency spectrum (mSFS). The observed mSFS was computed using the SFStools script from D. Marques available on github (https://github.com/marqueda/SFS-scripts). For these demographic model comparisons, we used only individuals from Osprey Lake (*C. desquamator*: n=10; *C. variegatus* : n=12; *C.* sp. ‘wide-mouth’: n=11; *C. brontotheriodes*: n=13) to avoid additional complex population structure across ponds. We contrasted 28 demographic models of different topologies and gene flow scenarios across the four groups (Fig S5;Table 1 and S2). For each model, the fit to the observed multidimensional mSFS was maximized using the composite-likelihood method fastsimcoal (v2.6.0.3;[12] with 100,000 coalescent simulations, 40 expectation-maximization cycles, and pooling all entries with less than 20 SNPs). For parameter estimates, we used a wide search range with log-uniform distributions with the range of some priors informed by previous estimates of effective population size from MSMC and the age of the last glacial maximum (∼20kya) prior to which the lakes would have been dry on SSI. These ranges were not upper bounded by these specified priors and so the simulations were free to explore parameter space that exceeded the priors.

For each demographic model, we ran 100 independent *fastsimcoal2* runs to determine the parameter estimates with the maximum likelihood. The best fitting demographic model was identified using Akaike information criteria (AIC). To get confidence intervals for the parameter estimates from the best fitting model, we simulated 100 mSFS based on the maximum likelihood parameter estimates from our best fitting model and ran 50 independent runs with this model on each simulated SFS. The parameter point estimates from the run with the highest likelihood (of the 50 independent runs) from each simulated SFS were then used to compute 95 percentile confidence intervals as a measure of uncertainty in the parameter estimates from the observed SFS.

#### Testing for shared adaptive alleles

##### Patterns of directional selection

Various demographic histories can shift the distribution of low- and high-frequency derived variants to falsely resemble signatures of hard selective sweeps. In a previous study, we used MSMC analyses to infer histories of population size changes in *C. variegatus*, *C. desquamator* and *C. brontotheriodes* (Richards et al. 2021). In this study, we repeated the same analysis for the ‘wide-mouth’. In order to account for demography of the ‘wide-mouth’ population in downstream analyses, we used the MSMC (v. 1.0.1;[13] to infer historical effective population size (N_e_) changes in the ‘wide-mouth’. We ran MSMC on unphased GATK-called genotypes separately for a single individual ‘wide-mouth’ from Osprey Lake with 16x mean coverage across its genome (Fig S4). As recommended in the MSMC documentation, we masked sites with less than half or more than double the mean coverage for that individual or with a genotype quality below 20. We also excluded sites with <10 reads as recommended by Nadachowska-Brzyska et al. [14]. To scale the output of MSMC to real time and effective population sizes, we used a one-year generation time [15] and the estimated spontaneous mutation rate of 1.56 x10^-8^ estimated from high coverage sequencing of two independent pedigreed parent-offspring crosses of San Salvador Island species from a previous study (Richards et al. 2021).

Across all four populations in Osprey Lake, we looked for regions that appeared to be under strong divergent selection in the form of a hard selective sweep from the site frequency spectrum using SweeD (v.3.3.4;[16]). In this calculation of the composite likelihood ratio (CLR) of a sweep, we incorporated our empirical estimate of the decrease in population size for each focal population estimated from MSMC analyses in 50-kb windows across scaffolds that were at least 100-kb in length (99 scaffolds; 85.6% of the genome). We also calculated CLRs across 100,000 scaffolds consisting of neutrally evolving sequences simulated with ms-move [17], controlling for the impact of the inferred population size decreases over time for each population from MSMC runs mentioned above (Fig. S4; Table S3). The CLR ratios for the simulated datasets were then used to assess outlier CLR ratios from the empirical dataset. Regions with CLR ratios above the 95^th^ percentile value of CLR from the neutral simulated dataset were considered candidate hard selective sweep regions (Table S3). We compared selective sweeps across *C. variegatus, C. desquamator, C. brontotheriodes*, and ‘wide-mouth’ to look for shared and unique selective sweeps among the four groups.

##### Characterization of candidate adaptive alleles

We searched for candidate adaptive alleles underlying divergent traits among the species by overlapping selective sweeps regions with regions of high genetic divergence based on fixed or nearly fixed SNPs between groups. We chose to look at nearly fixed SNPs over only fixed variation to accommodate the ongoing geneflow occurring between these young species. We took two approaches to finding fixed or nearly-fixed variants: 1) a fixed threshold of *F_st_* ≥ 0.95 across all comparisons and 2) and threshold of the 99.9^th^ percentile of *F_st_*, which varied among comparisons (range of *F_st_:* 0.73-0.83).

We made the following pairwise comparisons for *F_st_* calculations including a) between *C. variegatus* and each of the specialists, b) each specialist against all other groups and c) shared between two or three specialists against *C. variegatus*. For nearly fixed variants with the *F_st_* ≥ 0.95 threshold, we looked for putative function of the candidate adaptive alleles by looking at gene annotations of any gene the variant was in or near (within 20-kb of the gene, which is within the 50-kb LD decay estimate). We used available gene annotations from model organisms of mice and zebrafish from MGI, ZFIN, and we checked other annotation databases and studies for verification of putative function, including Phenoscape Knowledgebase (https://kb.phenoscape.org/#/home), NCBI’s PubMed (https://www.ncbi.nlm.nih.gov/pubmed), and the Gene Ontology database using AMIGO2 [18] .

For the 99.9^th^ percentile *F_st_* variants: We performed gene ontology (GO) enrichment analyses for genes near candidate adaptive variants using ShinyGo (v.0.51;[19]). In the *C. brontotheroides* reference genome annotations (described in *de novo* genome assembly and annotation section), gene symbols largely match human gene symbols. Thus, the best matching species when we searched for enrichment across biological process ontologies curated mostly human gene functions.

##### Timing of selection on candidate adaptive alleles

Lastly, we also determined the relative age of candidate adaptive alleles by generating estimates of coalescent times using the R package starTMRCA (v0.6-1;[20]). For each candidate adaptive allele that was unique to the three specialists and the 16 shared alleles between *C. desquamator* and ‘wide-mouth’, a 1-Mb window surrounding the variant was extracted into separate vcfs for individuals within each species. The program requires complete genotype data so we first filtered out any individuals with more than 10% missing data (1 *C. brontotheriodes* and 2 ‘wide-mouth’ individuals). With Tassel5 [21], we then used the LD KKNI command to infer missing sites based on LD for remaining individuals with less than 10% missing data. Subsequently we removed the small number of sites with any missing data across individuals within each population. Since several adaptive alleles fall within the same 1-Mb window, we chose a single adaptive allele at random within each 1-Mb window as the focal allele to estimate timing for. This lead to 6 time estimates for the 16 shared adaptive alleles between *C. desquamator* and *C.* sp. ‘wide-mouth’, 5 time estimates for the unique adaptive alleles to *C.* sp. ‘wide-mouth’, X time estimates for the unique adaptive alleles to *C. desquamator* and 3 time estimates for the unique adaptive alleles to *C. brontotheroides*.

These sets of variants were then analyzed in starTMRCA with the mutation rate of 1.56x10^-8^ substitutions per base pair, and a recombination rate of 3.11x10^-8^ (from genome-wide recombination estimate for stickleback; [22]). Each analysis was run three times per focal adaptive allele and all runs were checked for convergence between and within runs. Most runs rapidly converged within the initial 6000 steps, but 5 runs did not converge after an additional 4000 steps and were discarded from further analysis.

#### Characterization of introgression in C. sp ‘wide-mouth’ genome

##### Introgression with C. desquamator in candidate adaptive allele regions

To assess whether shared adaptive alleles between *C.* sp ‘wide-mouth’ and *C. desquamator* may be due to introgession between the two populations, we calculated Relative Node Depth (RND) statistics [23] in 10-kb sliding windows across the genome. RND statistics measure the relative node depth of two taxa compared to an outgroup and is calculated as the quotient of *D_xy_* between two species to the average distance from each to an outgroup. This relative amount of divergence compared to an outgroup allows the statistic to be robust to mutation rate variation across the genome, in which low neutral mutation rate regions might be mistaken for a locus that has experienced a recent introgression event. RND was calculated using a custom script that used R package PopGenome (v2.7.5;[24]) first calculate *D_xy_* values calculated for *C. desquamator*, *C.* sp. ‘wide-mouth’ and *C. artifrons* in 10-kb windows across the genome and then RND for each scaffold that contained a candidate adaptive variant that was unique or shared between *C. desquamator* and *C.* sp wide-mouth. Significance of RND values was evaluated using simulations with no migration using ms-move [17]. We used estimates of changes in effective population size and divergence times for each population from our *fastsimcoal2* analyses and ms-move to simulate neutral genetic divergence between two isolated populations. The threshold for significant introgression regions was determined by simulating genetic sequences under a coalescent model with no gene flow, consisting of 150,000 10-kb windows each containing the mean number of alleles observed in our dataset and running these simulated sequences through the same custom PopGenome script (RND_PopGenome_Artout_null.R) to calculate RND values. Empirical windows were considered candidates for introgression if the RND statistic was in the bottom 5^th^ percentile of simulated values (RND < 0.22; Table S10). To assess whether the sympatric coexistence of four incompletely reproductively isolated species in Osprey Lake has lead to introgression across specialist and generalist population boundaries as well, we assessed whether RND test statistics calculated between *C. variegatus* and *C. desquamator* in Osprey Lake were similarly detected as introgressed in these regions. None of these regions appeared to be introgressed with the generalist *C. variegatus* (Table S10), suggesting the introgressed signature between *C. desquamator* and *C*. sp. ‘wide-mouth’ is not likely due to neutral introgression likely occurring between all species in the lake due to their incomplete reproductive isolation.

##### Characterizing introgression with outgroup Caribbean generalist populations in candidate adaptive allele regions

We also investigated whether any of these candidate adaptive alleles in *C. desquamator* and *C.* sp ‘wide-mouth’ fell in regions that appear to be introgressed from an outgroup generalist population (similar to analyses done in our previous study [25]). First, we extracted the genomes of individuals assigned to our 5 focal outgroup generalist populations in which we had 8 or more individuals sequenced in that previous study. We filtered variants across these 5 populations and our San Salvador Island samples down to those that has a quality of 20, and no more than 10% missing in any population for a dataset of 10,876,882 SNPs across 10 populations (including the four SSI species and the outgroup *C. artifrons*) and the 26 scaffolds that candidate adaptive alleles were found on. To detect introgression involving outgroup populations we used the differential test of introgression, *df*-statistic [26], which is designed to look for signatures of introgression across sliding genomic windows [27]. The *df-*statistic is a modified version of the *D*-statistic, looks at allele frequencies fitting two allelic patterns referred to as ABBA and BABA based on the tree (((P1,P2),P3),O), where O is an outgroup species in which no gene flow is thought to occur with the other populations [27]. We used 2 individuals of *C. artifrons* from Cancun, Mexico as our outgroup population for this test, which forms the deepest divergence event with *C. variegatus* within the *Cyprinodon* clade [28], and focused on introgression between *C. desquamator* or *C. sp.* ‘wide-mouth’ and outgroup Caribbean generalist populations that was not shared with *C. brontotheroides*. Based on the tree (((P1,P2),P3), *C. artifrons*), the *df* statistic was calculated for the combinations of populations in which the focal population (P2) was either the scale-eater or the molluscivore, the other specialist population was the sister group (P1), and P3 was one of the Caribbean outgroup populations sequenced in a previous study (Fort Fisher, North Carolina Coast;Isla Margarita, Venezuela; Lagunas Bavoros, Dominican Republic, and New Providence Island, Bahamas. *Fd-*statistics were calculated from 10-kb sliding windows using R package PopGenome. Empirical windows were considered candidates for introgression if the *df*-statistic was in the top 95^th^ percentile of simulated values (*df* > 0.55).

##### Characterization of admixture history on San Salvador Island with outgroup generalist populations

The results from sliding window tests for introgression surrounding candidate adaptive alleles for *C.* sp. ‘wide-mouth’ indicated that this cryptic species may be the product of admixture between SSI species and outgroup generalist lineage related to our North Carolina coast population (Table S8 and S10). To further explore this, we created genome-wide admixture graphs using the program qpgraph [29] in R package admixtools2 (v2.0.0; https://uqrmaie1.github.io/admixtools/). Qpgraph can accommodate many populations and admixture events and creates an admixture graph that summarizes all pairwise *f_2_*_-_ and *f_3_*-statistics across included populations and estimates admixture proportions and drift weights between populations. Admixture graphs model the number of admixture events specified by the user and evaluates the fit of the observed *f*-statistic values between populations to those expected from this model to produce a negative log likelihood score that measures the total amount by which the modeled f-statistics failed to match the observed values (where larger likelihood values indicate poorer fit).

Normally in qpgraph analyses, the user manually specifies the topology of the model and the program then solves for the optimal values of the parameters. The recent R package admixtools2 also includes the function find_graphs() that generates and evaluates admixture graphs across user-defined number of iterations to find the a range of feasible and best fitting graph topologies for a set of *f*-statistics. Using the four Osprey Lake populations on San Salvador Island (*C. variegatus, C. brontotheroides, C. desquamator* and *C.* sp. ‘wide-mouth’, and the 5 Caribbean generalist populations and *C. artifrons* outrgroup mentioned in the above two sections, we ran find_graphs() 10 times to model 1-10 admixture events.

The input data consisted of 20,381 SNPs that remained after filtering the dataset that included the outgroup generalist populations used in the sliding-window tests above and an additional filters for heterozygous sites that had an allele balance between 0.3 and 0.7, minimum 3x depth of coverage, and quality filter of 30. The final dataset used as input for qpgraph included 20,381 SNPs across 10 populations and each find_graphs() run was allowed to explore topology space for 1000 iterations. The topology with the best fit was extracted from these runs and assigned to be the admixture graph for that admixture event. The model and admixture graph with the lowest likelihood of all admixture events tested (LnL: range 2.16-16.23 across all 10 models), included 7 admixture events. One of these admixture events depicted admixture between *C. desquamator* and a lineage related to the North Carolina coast generalist population resulting in the C. sp ‘wide-mouth’ population. Other admixture events in this admixture graph, such as an admixture event at the base of the SSI radiation and a separate admixture event involving *C. desquamator* and an outgroup generalist, supported signatures inferred using independent program TreeMix from a previous study[25].

The inference of *C.* sp. ‘wide-mouth’ as admixed between *C. desquamator* and a distantly related generalist species appeared in the 8 admixture graphs modeling 3-10 admixtures events, suggesting it’s a strong signature in the dataset. However, the outgroup Caribbean generalist population that acted as one of the donors varied across different runs from being sister to North Carolina coast population to falling somewhere along the branch that divides the Venezuela population from the rest of the Caribbean populations. This may suggest that we are missing the true donor population in our sample. To further support this signature of *C.* sp. ‘wide-mouth’ experiencing introgression following a potential secondary contact event between San Salvador Island and an outgroup generalist population, we used *f_4_*-statistics to look for genome-wide signatures of differential introgression *between C. desquamator* and *C*. sp ‘wide-mouth’.

Differential signatures of introgression between the two would likely indicate introgression that *C*. sp. ‘wide-mouth’ contains in its genome but is not shared with *C. desquamator* and therefore likely not a signature of shared introgression in the ancestor of the two scale-eating populations on SSI. Genome-wide *f_4_* statistics were calculated using the f4() function in admixtools2 which calculates *f_4_*-statistics, Z scores and p-values for the specified populations. Following the tree ((P1,P2),(P3,P4)), *C. desquamator* was used as sister species P1, *C.* sp. ‘wide-mouth’ as focal species P2, the 5 generalist populations from Fort Fisher, North Carolina Coast; Isla Margarita, Venezuela; Lagunas Bavoros, Dominican Republic, Rum Cay, Bahamams and New Providence Island, Bahamas were rotated in as the potential donor population P3 and *C. artifrons* used as population P4 (Table S11). The *f_4_*-statistic that included North Carolina coast population had highest Z-score and indicated a significant amount of gene flow (p-value=0.004;Table S11). However, tests that included Rum Cay and New Providence Island also resulted in *f_4_*-statistics with marginally significant p-values (p-value=0.056 and 0.067 respectively; Table S11), perhaps mirroring that admixture occurred with an unsampled generalist population somewhere North of SSI and most closely related to North Carolina population.

### Results

#### ‘Wide-mouth’ ecomorph is ecologically intermediate and morphologically distinct

Morphological traits were heritable in a common garden laboratory environment after one generation: lab-reared *C.* sp. ‘wide-mouth’ displayed significantly larger buccal width than *C. desquamator* (t-test; *P* = 0.003) and maintained their characteristic intermediate jaw lengths (ANOVA; *P* = 0.03, figure S3). There was also some evidence of phenotypic plasticity in both *C. desquamator* and *C.* sp ‘wide-mouth’ compared to wild individuals. Two of the focal three traits that distinguished wild populations of *C.* sp. ‘wide-mouth’ from the other pupfish species, lower jaw length and buccal width, remain significantly distinct in the lab-reared populations (based on non-overlapping 95% confidence intervals around mean residuals of each species). However, one trait that distinguishes *C.* sp. ‘wide-mouth’ in the wild appears to be quite plastic under laboratory rearing conditions; there were no significant differences in mean adductor mandibulae height residuals among species in laboratory environments. This is interesting because adductor mandibulae height is a proxy measurement for the cross-sectional width of the adductor muscle directly beneath the skin. The plasticity observed in this proxy may be due to muscle plasticity. In the lab-reared colonies, all species are reared on the same diet, such that the scale-eating populations do not scale-feed as often as they would in the wild and thus lab-reared populations of scale-eaters may not develop the notably increased adductor muscle mass. Interestingly, the distinct differences in adductor mandibulae insertion height among wild populations disappeared in the lab-reared populations (figure S3), indicating phenotypic plasticity associated with the common diet of pellet foods and lack of scale-eating attacks in the lab.

We found that ‘wide-mouth’ ingest scales, but at a significantly lower frequency than *C. desquamator* (Wilcoxon Rank Sum test, *P* = 0.004; figure 3A). We did not detect any scales in *C. variegatus* guts (figure 3A). Detritus made up the only other gut contents present besides scales in all the specimen of *C.* sp ‘wide-mouth’ from Osprey Lake that we dissected. This was similar for the other scale-eater *C. desquamator* individuals from Osprey Lake, in which we also found only scales and detritus in their gut. For the generalist species *C. variegatus*, their guts contained only detritus except for a single individual in which we found a snail shell. The low diversity in gut contents among species may be due to the relatively small sample size of individuals we had that we could perform gut content analysis on for this study (n=10 for each species). However, the predominance of detritus in the gut contents of Osprey Lake populations of all three species is not surprising given the thick (<1m) bottom layer of mud and flocculent in this lake[30]. Additionally, detritus made the largest percentage (49-71%) of San Salvador Island pupfish diet in other lakes across the island as well. A previous study that conducted gut content analyses with 2-4X larger sample sizes also found that the majority of *C. variegatus, C. brontotheroides,* and *C. desquamator*’s diet is detritus. Except for one population of *C. desquamator* in Crescent Pond where scales made up 50% of their gut contents and detritus only made up 30%. Therefore, the ecological intermediacy of *C.* sp ‘wide-mouth’ is supported by their intermediate ratio of scales to detritus, which is the predominant axis of dietary divergence between *C. variegatus* and *C. desquamator* in this lake. The lack of other dietary items in the gut contents of *C.* sp ‘wide-mouth’ further supports our arguments that their ecological divergence is along the same specialist axis of scale-eater *C. desquamator* and that they share little ecological overlap with the other molluscivore specialist *C. brontotheroides*.

#### ‘Wide-mouth’ did not result from hybridization between *C. variegatus* and *C. desquamatory*

Several lines of genomic evidence support the ‘wide-mouth’ ecomorph as a distinct species. First, ‘wide-mouth’ individuals do not occupy an intermediate position between *C. desquamator* and *C. variegatus* along either of the first two principal components that represent the major axes of genetic variation in the radiation (figure S4A). In the ADMIXTURE analysis, ‘wide-mouth’ individuals share more ancestry with each other than with either *C. desquamator* or *C. variegatus* under the two most likely K values (K=6 and 7; figure S4B&5). Lastly, significant positive *f_3_*- statistics and the non-significant *f_4_*-statistics support our inference that the ‘wide-mouth’ is not a recent admixed population of any pairwise combinations of the *C. brontotheriodes*, *C. variegatus*, or *C. desquamator* populations (table S1).

#### *C*. sp. ‘wide-mouth’ did not result from hybridization between *C. variegatus* and *C. desquamator*

We compared 28 different demographic models of divergence and gene flow on San Salvador Island (table S2) that represented all possible topologies among the four species and explored zero, early, and current gene flow scenarios. In the best supported model, ‘wide-mouth’ was sister to *C. desquamator* with current gene flow and a divergence time estimate of 11,658 years ago (95 CI: 8,257-20,113 years; figure 4; table 1 and S2). This model indicates that the ancestor to ‘wide-mouth’ and *C. desquamator* populations first diverged from *C. variegatus* 15,390 years ago (95% CI: 10,722-23,927 years; figure 4B). This estimate of the origin of the radiation overlaps well with geological age estimates based on the filling of hypersaline lakes on San Salvador Island after the end of the last glacial maximum period (∼6-19K years ago; [31, 32]). *C. variegatus* and *C. brontotheriodes* populations in Osprey Lake diverged recently about 462 years ago (95% CI: 411-1,121 years; figure 4B).

#### Shared and unique adaptive alleles in *C.* sp. ‘wide-mouth’ and *desquamator*

We discovered that the two scale-eating populations shared a set of the same adaptive alleles not found in *C. brontotheriodes* or *C. variegatus* (Fig 5D). This consisted of 15 alleles in 6 shared hard selective sweeps in *C. desquamator* and ‘wide-mouth’: 10 SNPs were in unannotated regions, two were in the introns of the gene *daam2,* and three were in regulatory regions of the genes *usp50, atp8a1,* and *znf214* (one variant each).

Shared adaptive alleles in the gene *daam2,* a wnt signaling regulator, are intriguing because knockdown of this gene causes abnormal snout morphology, osteoporosis, and changes in insulin and alkaline phosphate levels in mice [33], and abnormal cranial and skeletal development in zebrafish [34]. Craniofacial morphology is one of the rapidly diversifying traits in this system, suggesting that divergence in *daam2* may play a role in the shared craniofacial divergence of the two scale-eating species. Similarly, *usp50* functions in protein metabolism and deubiquination [35] and may play a role in shared metabolic adaptations to the higher-protein content of a scale-eating or snail-eating diet (also note microbiome divergence of *C. desquamator* with increased prevalence of collagenase-digesting bacteria when reared in a common garden:[36]). *atp8a1* is an ion transporter [37], *znf214* is a transcription binding factor, and the ten unannotated variants were not associated with lower jaw size variation in a previous genome-wide association study of the radiation (Richards et al. 2021).

Despite the divergent craniofacial features of the ‘wide-mouth’, none of the adaptive alleles unique to the ‘wide-mouth’ appear to be in or near genes annotated for craniofacial phenotypes in model organisms. In *C. desquamator,* three of the 13 sets of unique adaptive alleles are in or near genes annotated for craniofacial phenotypes: a *de novo* non-synonymous coding substitution in the gene *twist1*, several putative regulatory variants near the gene *gnaq,* and 8 variants in and near the gene *bri3bp*, which is located inside a QTL region for cranial height in pupfish (St. John et al. 2021). In *C. brontotheriodes*, there is also a candidate craniofacial adaptive allele: a non-synonymous coding substitution in the gene *kat6b*, which is associated with abnormal craniofacial morphologies, including shorter mandibles in mice [38, 39] and Ohdo Syndrome and bulbous noses in humans [40].

This pattern of unique alleles relevant to craniofacial phenotypes in *C. brontotheriodes* and *C. desquamator*, but not ‘wide-mouth’, holds even if we lower the threshold to the top 1 percentile of *F_st_* between specialists and generalist (see supplemental for more details). ‘Roof of mouth development’, ‘bone development’ and ‘skeletal system development’ are in the top 20 most significantly enriched GO terms for *desquamator* candidate adaptive alleles, with 6 significantly enriched terms relevant to cranial and skeletal development (out of 165 enriched terms total). Similarly, ‘embryonic skeletal system development’ and ‘skeletal system development’ were significantly enriched terms for *brontotheriodes* candidate adaptive SNP alleles (out of 8 enriched terms total). However, ‘wide-mouth’ adaptive alleles at this lower *F_st_* threshold were not significantly enriched for any GO terms related to craniofacial or skeletal morphology (n=52 terms; Fig S7).

#### Substantial history of admixture in *C.* sp ‘wide-mouth’

To further investigate the signatures of adaptive introgression in this species, we modeled admixture across San Salvador Island and several key generalist populations in the Caribbean with *f_3_*- and *f_4_*-statistics (table S11;figure S10). Previous signatures of admixture involving the base of SSI radiation and outgroup generalist populations [25] were detected again in our admixture graph, alongside a new signature of admixture that appears to be secondary contact between a pupfish lineage most closely related to North Carolina in our dataset and *C*. sp ‘wide-mouth’ (table S11; figure 5 & S10).

#### Shared signatures of selection across the three specialists in the radiation

We found that a higher percentage of the genome was under positive selection in all specialist populations compared to the *C. variegatus* population (figure 5A). This pattern might be expected given the divergent selection pressures the specialists face to adapt to different trophic niches. Of all the specialists, *C. brontotheriodes* exhibited both the most selective sweeps and the longest selective sweeps (figure 5A). This suggests that *C. brontotheriodes* have undergone selection most recently among all the species in Osprey Lake and is supported by the recent divergence time between the *C. brontotheriodes* and *C. variegatus* populations (figure 4D). The shared selective sweeps were shorter than any of the selective sweeps unique to each of the specialists, suggesting that selection for these shared regions was not the most recent or strongest in any of the specialists (figure S11C).

Additionally, all specialists, including the two scale-eating species, were more genetically diverged from each other than to *C. variegatus* species based on the number of SNPs fixed or nearly-fixed between them (figure 5B). This pattern of stronger genetic divergence between specialists also held across fixed SNPs, the top 1% of *F_st_*, and genome-wide average *F_st_* (table S4). Next, we identified a set of candidate adaptive alleles for each specialist by filtering for SNPs that were fixed or nearly fixed (*F_st_*, ≥ 0.95) relative to *C. variegatus* and found within a hard selective sweep in the focal specialist. In this set of candidate adaptive alleles, *C. brontotheriodes* had the fewest adaptive alleles while *C. desquamator* had the most (figure 5B). All three specialists, including the two scale-eating species, were more genetically diverged from each other than to *C. variegatus* based on the number of SNPs fixed or nearly-fixed between them across multiple *F_st_* thresholds (figure S11B; table S4).

### Discussion

The hallmark of adaptive radiation is a rapid burst of diversification which is predicted by theory to slow down over time as niche subdivision increases [41]. An alternative possibility is that radiations can be self-propagating and that the diversity generated within the first stages of radiation helps beget further diversity [42]. This could happen through exploitation of new trophic levels created by new species or physical alterations of the environment by new species that may create additional opportunities for speciation (reviewed in [41, 43]). The diversity begets diversity hypothesis can also be visualized as the exploration of a complex multi-peaked fitness landscape; as species in the radiation colonize new peaks, this provides access to additional neighboring fitness peaks to fuel rapid radiation. At the microevolutionary level, shared adaptive variants can also help other populations colonize new or ecologically similar areas of the fitness landscape. However, the genetic basis and microevolutionary processes underlying major ecological transitions are still poorly understood in nature.

#### An adaptive walk underlies the major ecological transition from generalist to scale-eating specialist

One of the foundational models of adaptation is that it proceeds in ‘adaptive walks’ towards fitness optima that involve the sequential fixation of adaptive alleles that move a population in the phenotypic direction of the local optimum [44–48]. The distinct timing of selection across different adaptive alleles suggests that the ecological transition from generalist to novel scale-eating specialist involved such an adaptive walk rather than a sudden burst of concurrent selection events after some major environmental shift. These distinct, multiple bouts of selection could be caused by mutation-limited (i.e. waiting for new beneficial alleles; [49–51]) or mutation-order processes (i.e. epistatic interactions; [52–54].

We found that all of the shared adaptive alleles between the scale-eating populations occurred at low frequency in generalist populations on other Caribbean islands in a previous study ([25]; Table S7). For example, three copies of the shared *usp50* adaptive allele were found outside of San Salvador Island (Table S7). This indicates that initial bouts of selection occurred on available standing genetic variation and that staggered timing of hard selective sweeps in each trophic specialist most likely reflects mutation-order processes in which selection on a beneficial allele was contingent on prior fixation of other adaptive alleles in each specialists’ genetic background. However, several adaptive alleles originated from introgression or de novo mutations found only on San Salvador Island (Table S7), so part of the adaptive walk may have also been mutation-limited.

#### Cryptic scale-eating species reveals features of the adaptive walk towards scale-eating specialization

Although the two scale-eating species shared a set of adaptive alleles and a recent common ancestor, 5 of 6 shared adaptive regions swept at significantly different times between the two species. This difference in timing may result from several different scenarios: 1) independent adaptive walks to the same scale-eating niche, 2) independent adaptive walks to different scale-eating niches or 3) the adaptive walk of one population depends on the adaptive walk of the other population. We explore each of these scenarios in turn.

First, the difference in timing of selection on the same shared adaptive alleles could indicate independent adaptive walks to the same scale-eating peak that occurred at different times and/or by slightly different routes. *C. desquamator* and ‘wide-mouth’ populations have predominantly abutting ranges, with only a small amount of geographic overlap on San Salvador Island in four lakes, Osprey, Oyster, Little Lake, and Mermaid’s Pond (Fig 1). If this current distribution is representative of their historical ranges and the two lineages began diverging about 11,000 years ago, it is possible that the adaptive walks took place in different lakes and were largely independent of one another. The differences in timing on the shared adaptive alleles and the presence of unique alleles observed in each population are reminiscent of mutation-order speciation [53, 55]. In mutation-order speciation the same alleles are favored in populations that are adapting to similar environments, yet by chance and/or epistatic interactions with different genetic backgrounds, similar adaptive alleles fix in just one population. At least one set of alleles near the gene *slitrk5* are unique to the ‘wide-mouth’ scale-eater and fixed well before selection occurred on the shared adaptive variants between the two scale-eater populations (Fig 5). Epistatic interactions of these *slitrk5* alleles may have prevented the ‘wide-mouth’ from fixing additional segregating alleles that are uniquely fixed in *C. desquamator*.

Second, the difference in timing of selection may have occurred because the two scale-eating populations are adapting to two different scale-eating niches. The intermediate diet, distinctly sized morphological traits, and smaller body size in the ‘wide-mouth’ scale-eater compared to *C. desquamator* may indicate the ‘wide-mouth’ scale-eater is adapting to a different scale-eating niche. While the two scale-eating populations do share some overlap in adaptive alleles and selective sweeps, the majority of selective sweeps are unique to each species (Fig 5 & 6), including neurogenesis, brain, and nervous system development (*slitrk5, sema4g, and smarce1*), whereas unique adaptive alleles in *desquamator* include craniofacial development annotations (*olfm1, gnaq, twist1*) as well. This difference in gene annotations can also be seen at the broader level of regions of the genome under selection (Fig S7). Therefore, the differences in timing at shared adaptive alleles might reflect differences in relative strength of selection due to different selective regimes experienced by the two scale-eating populations. While starTMRCA is fairly robust in its timing estimates to different strengths of the selection [20], we cannot rule out that differences in relative timing of selection might partially reflect differences in strength of selection occurring in the two scale-eating populations.

Third, another scenario that could have led to the older timing of selection across shared adaptive alleles in *desquamator* than ‘wide mouth’ is one in which the adaptive walk of one population depended on adaptive alleles from another. The significantly older timing of selection in *desquamator* on shared adaptive alleles and the low absolute genetic divergence between these two scale-eating population in these regions (Fig 6) suggests that these alleles may have introgressed between the two populations. Given that these shared adaptive alleles appear as standing genetic variation across Caribbean populations, albeit at very low frequency in our sampling, it is likely that the shared alleles were present in the ancestor and were segregating in both populations after they diverged. There are several reasons why adaptive introgression may have been necessary for adaptive divergence of ‘wide-mouth’ despite the alleles initially segregating in the ancestral population. In line with mutation-order processes (e.g. Mani and Clarke 1990; Schluter 2009; Good et al. 2017), these alleles might not have been adaptive until the right genetic background was present in the ‘wide-mouth’. Introgression from *C. desquamator* in which the allele was already swept to high frequency could have raised the frequency of these alleles in ‘wide-mouth’ and increased the likelihood of fixation for the beneficial alleles (reviewed in [56]). This is consistent with previously proposed hypotheses for how diversity begets further diversity in adaptive radiations (Whittaker 1977; Stroud and Losos 2016; Martin and Richards 2019).

#### Shared aspects of genomic landscape of selection among all specialists additionally supports specialization promoting further specialization

Intriguingly we also find evidence of selective sweeps shared across all three specialists despite their divergent adaptations and lack of a shared specialist ancestor. However, there are no commonly shared adaptive alleles fixed against *C. variegatus* in these shared selective sweep regions. Hard selective sweeps indicate a selection scenario in which a single beneficial allele of large effect on a trait is swept to high frequency (reviewed in [57]). The lack of highly divergent alleles in these regions might indicate polygenic selection events underlie the shared signatures across specialists. The broad-scale transition from dietary generalist to dietary specialist may involve polygenic selection with small shifts in the frequency of many alleles. Despite very different expectations about the genetic basis of adaptation, polygenic selection events may have been detected as a hard selective sweep in this study as the two selection types can be challenging to distinguish based solely on patterns of genetic variation [58–61]. Recently developed frameworks provide additional criteria to help distinguish between the two selective regimes [59] but are beyond the scope of this current study.

These shared selective sweeps across all specialists are not the longest selective sweeps in any of the specialist genomes (> 50-kb; Fig 5C), indicating they are not the most recent or strongest selection events. However, these shared selective sweeps appear to be relevant to dietary specialization as these regions are enriched for genes annotated for metabolic processes such as short chain fatty acid and propionate metabolism (Fig 5). Propionates and other short chain fatty acids are common microbiome metabolites [62]. A recent microbiome study in lab-reared populations found that *C. desquamator* microbiomes appear enriched for *Burkholderiaceae* bacteria that can digest collagen, further supporting an adaptive role for microbiomes in the radiation [36].

We also see pairwise shared selective sweeps between all combinations of two specialists. In contrast to the sweeps shared between all specialists, we see stronger genetic divergence among species in these regions and a proportion of the selective sweeps shared between two specialists do contain fixed or nearly-fixed alleles. However, the specialists that share selective sweeps in these regions are fixed for different adaptive alleles, consistent with previously observed patterns of parallel differential gene expression in specialists despite divergent genotypes [63]. These alleles were possibly under balancing selection in the ancestor (most of these alleles are segregating at intermediate frequency in *C. variegatus* currently) and the alternate alleles were driven to fixation between the specialists during their respective divergences. While the species have discrete phenotypes across several morphological traits important to divergence in this system, these traits often lie on a continuous axis (i.e. shorter oral jaw lengths in *C. brontotheriodes*, intermediate jaw lengths in *C. variegatus*, and longer oral jaw lengths in ‘wide-mouth’ and *C. desquamator*). Some of the alleles that are alternatively fixed between specialists at the same position may have incomplete dominance effects on such traits.

In line with adaptation to divergent trophic niches across the species, we do still see many unique signatures of selection among the specialists. The *C. variegatus* population had the fewest selective sweep signatures. All three specialists had more selective sweeps potentially resulting from strong directional selection for adapting to new fitness peaks. However, while we might expect more selective sweeps in *C. desquamator* genomes given the deeper valley on the fitness landscape isolating the scale-eater phenotypes [3, 64], C*. brontotheriodes* had the most selective sweeps detected in their genome. They also have the longest sweeps detected (e.g. 100-kb), suggesting that selection may have occurred most recently in the *C. brontotheriodes*. This matches with the most recent divergence event being between generalist and *C. brontotheriodes* in Osprey Lake. An intriguing implication of this result is the potentially long wait time for speciation of *C. brontotheriodes* species compared *C. desquamator* despite the shallower fitness valley isolating *C. brontotheroides* from generalists. However, we cannot yet rule out that the recent divergence time estimated in Osprey Lake reflects a more recent arrival of *C. brontotheriodes* to that lake after initially diverging long ago in another lake not incorporated in our demographic model.

**Fig S1.**
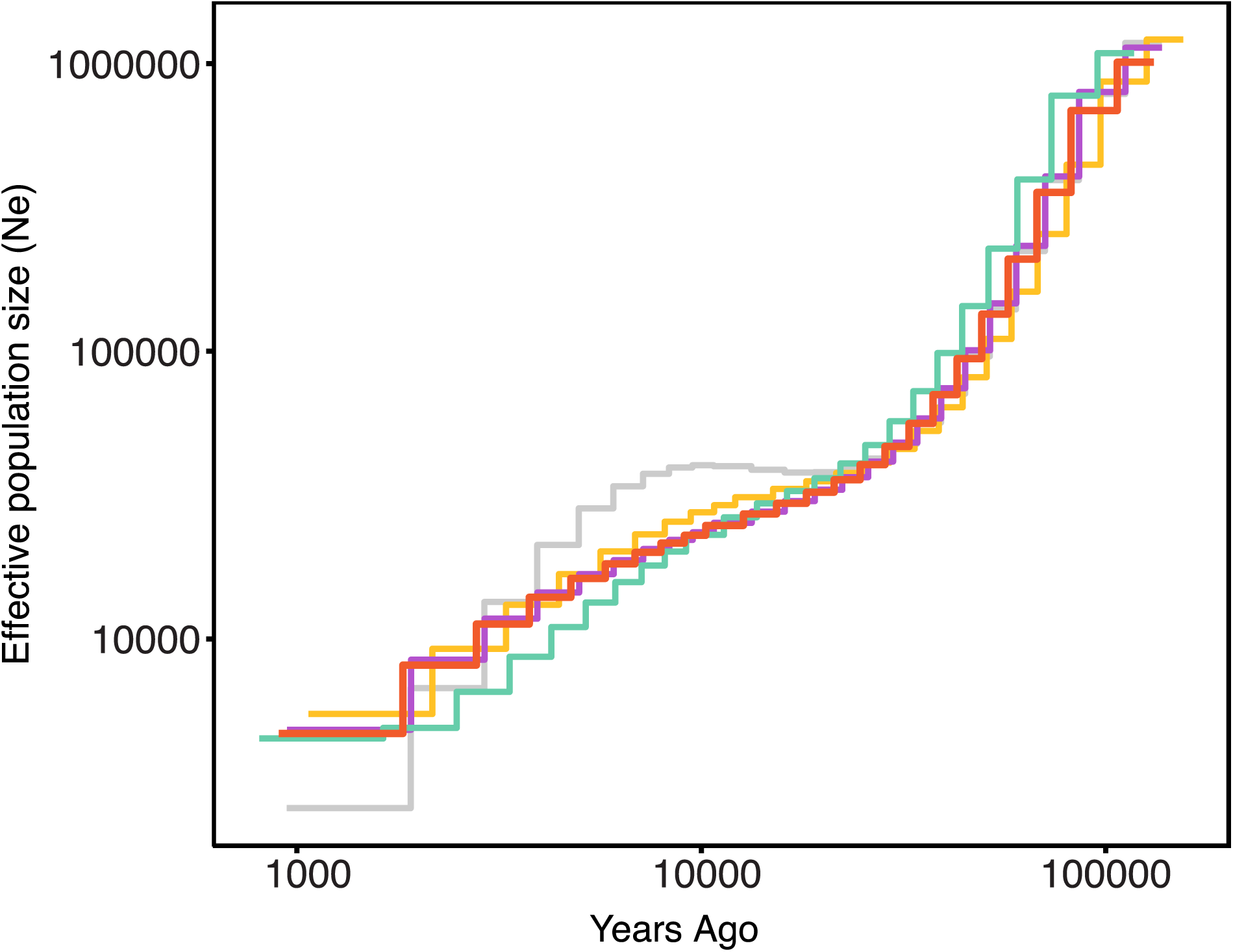
**Changes in effective population size over time for Osprey populations.** Inferred using MSMC [13] on high-coverage (18-36X) genomes from single individuals of each of the four species in Osprey Lake, San Salvador Island Bahamas: *C. variegatus* (gold), *C. brontotheroides* (purple), *C. desquamator* (teal), and C. sp. ‘wide-mouth’ (red-orange). Time was scaled using a mutation rate of 1.56x10-8 mutations/basepair/generation and a generation time of 1 year.

**Figure S2.**
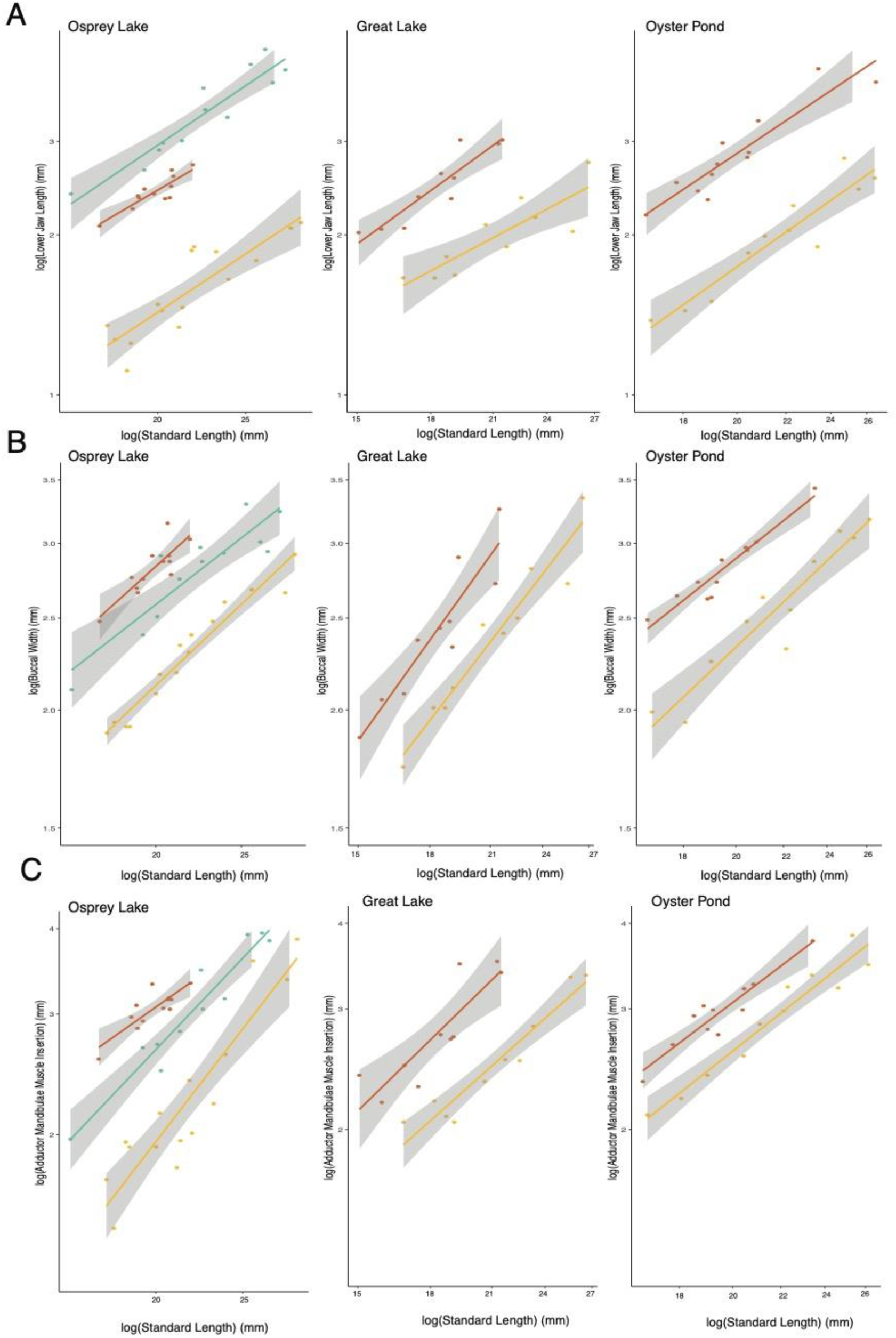
**Relationships between three divergent traits and standard length across ponds.** The relationship between standard length (mm) of individuals and their D) lower jaw length, E) buccal cavity width, and F) adductor mandibulae muscle insertion height across individuals the three species *C. desquamator* (teal), *C. variegatus* (gold), and *C.* sp. ‘wide-mouth’ (red-orange) across different ponds (Osprey, Oyster, and Great Lake). Colored lines represent linear model of these relationships for each species with their 95% confidence bands in gray.

**Figure S3.**
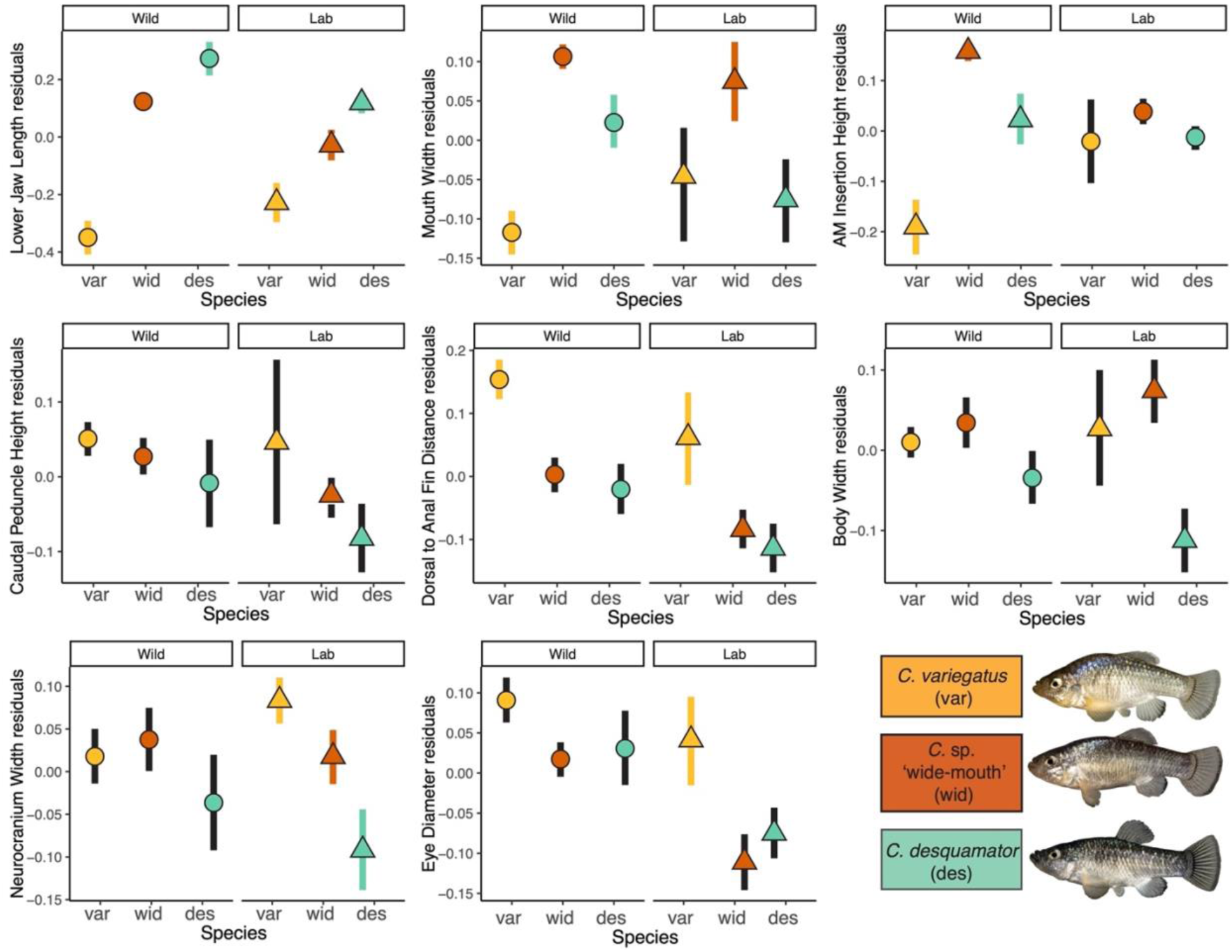
**Comparison of the three focal divergent traits in lab and wild populations.** A. 95% CI of the standardized sizes of 8 traits measured (see y-axis label on each panel for particular trait) for *C. variegatus* (gold;n=5), *C.* sp. ‘wide-mouth’ (red-orange;n=20) and *C. desquamator* (teal;n=20). Each plot is broken into two panels of wild caught and lab reared individuals (wild: circle; lab: triangle) represent the mean values of traits from each population. Confidence intervals that overlap between species are highlight in black. Lab raised individuals include F1, F2 and F3 generations. AM insertion height is a proxy for cross-sectional area of the adductor mandibulae

**Figure S4.**
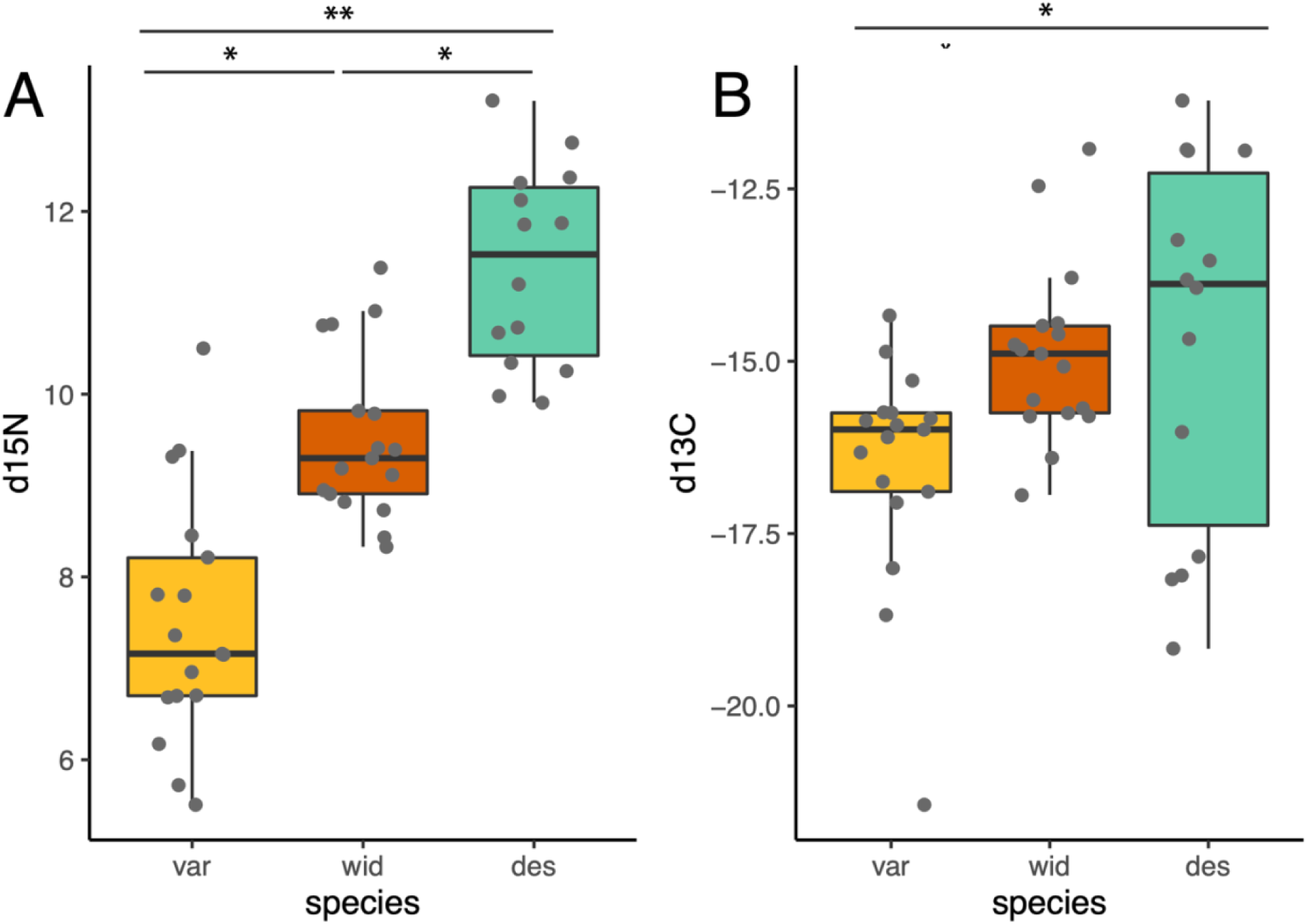
Trophic level comparison among populations in Osprey Lake. d15N values from isotope analysis of muscle tissue from 48 individuals across the three species that span the ecological spectrum from generalist to specialized scale-eater: generalist *C. variegatus* (var;gold), *C. sp.* ‘wide-mouth’ (wid;red-orange), and *C. desquamator* (des; teal). ANOVA results indicate significant differences in d15N levels across populations indicating that each occupies a distinct trophic level.

**Figure S5.**
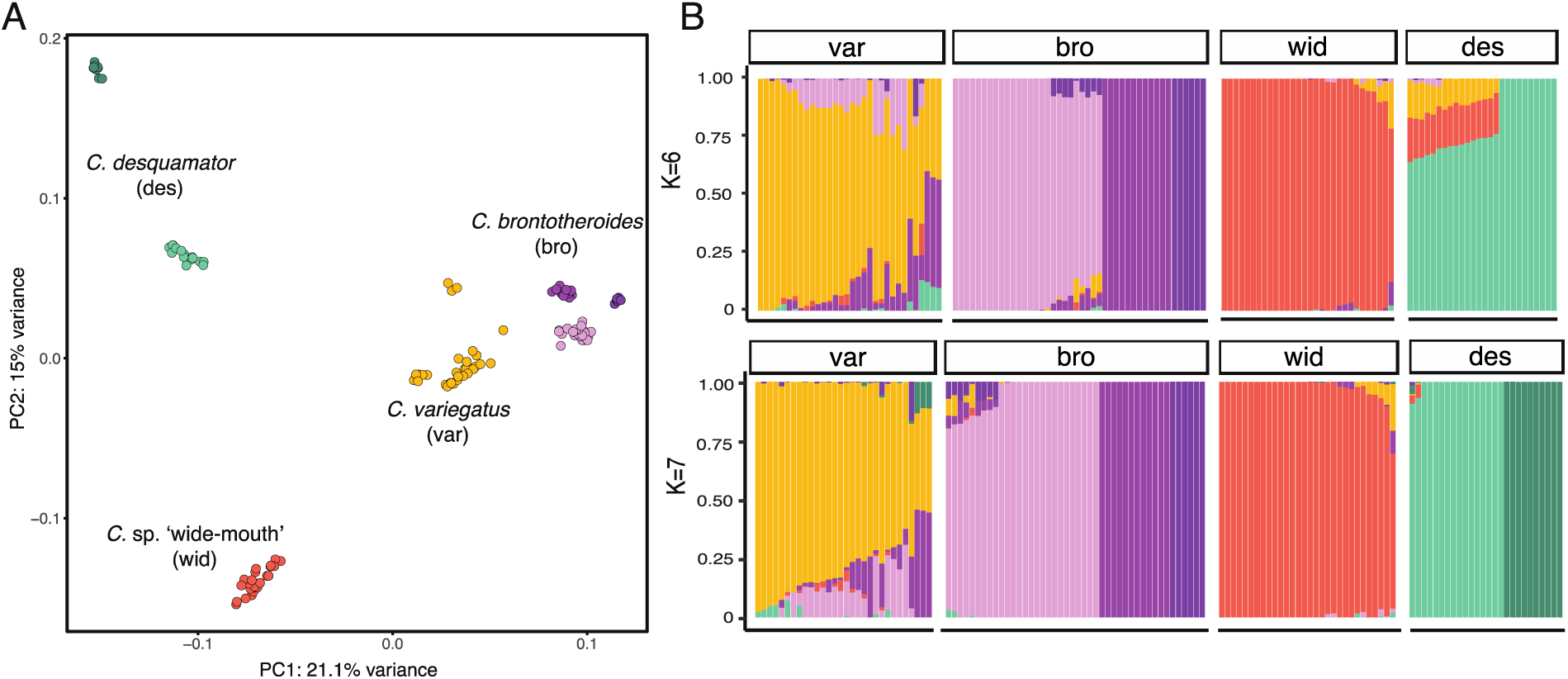
***C.* sp. ‘wide-mouth’ population did not result from recent hybridization.** A) Principal components analysis of the four focal groups on San Salvador Island based on an LD-pruned subset of genetic variants (78,840 SNPs). B) Ancestry proportions across individuals of the four focal groups. Proportions were inferred from ADMIXTURE analyses with 2 values of K with the highest likelihood on the same LD-pruned dataset in A.

**Figure S6.**
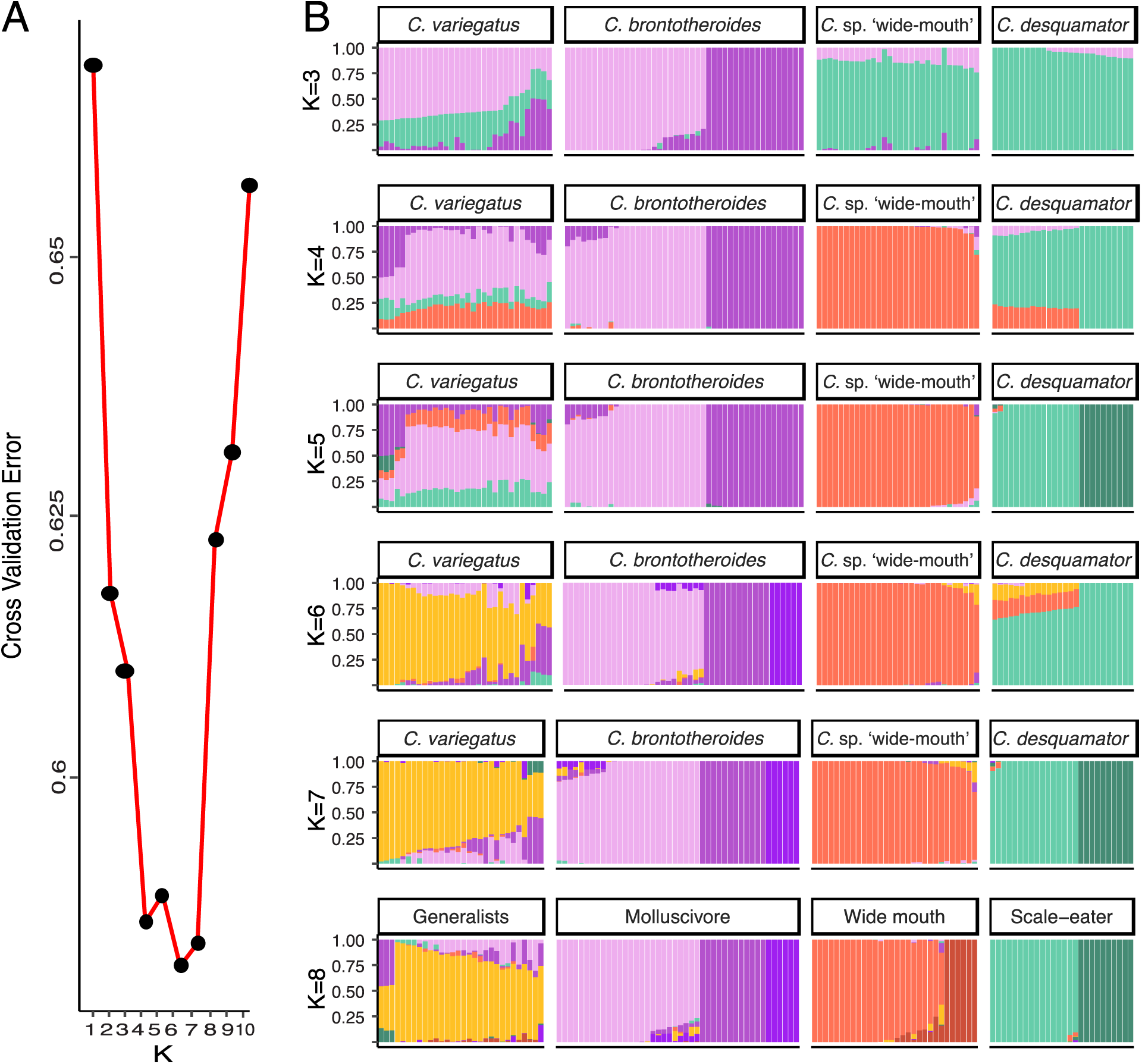
***C.* sp. ‘wide-mouth’ is not an admixed population**. ADMIXTURE analysis of San Salvador Island populations using an LD-pruned subset of genetic variants from 109 individuals (78,840 variants). A) Cross validation error estimates from ADMIXTURE that indicate a model of K=6-7 was the best fit. B) Ancestry proportions across individuals of the three focal groups. Proportions were inferred from ADMIXTURE analyses with 2 values of K with the highest likelihood on the same LD-pruned dataset in A.

**Fig S7.**
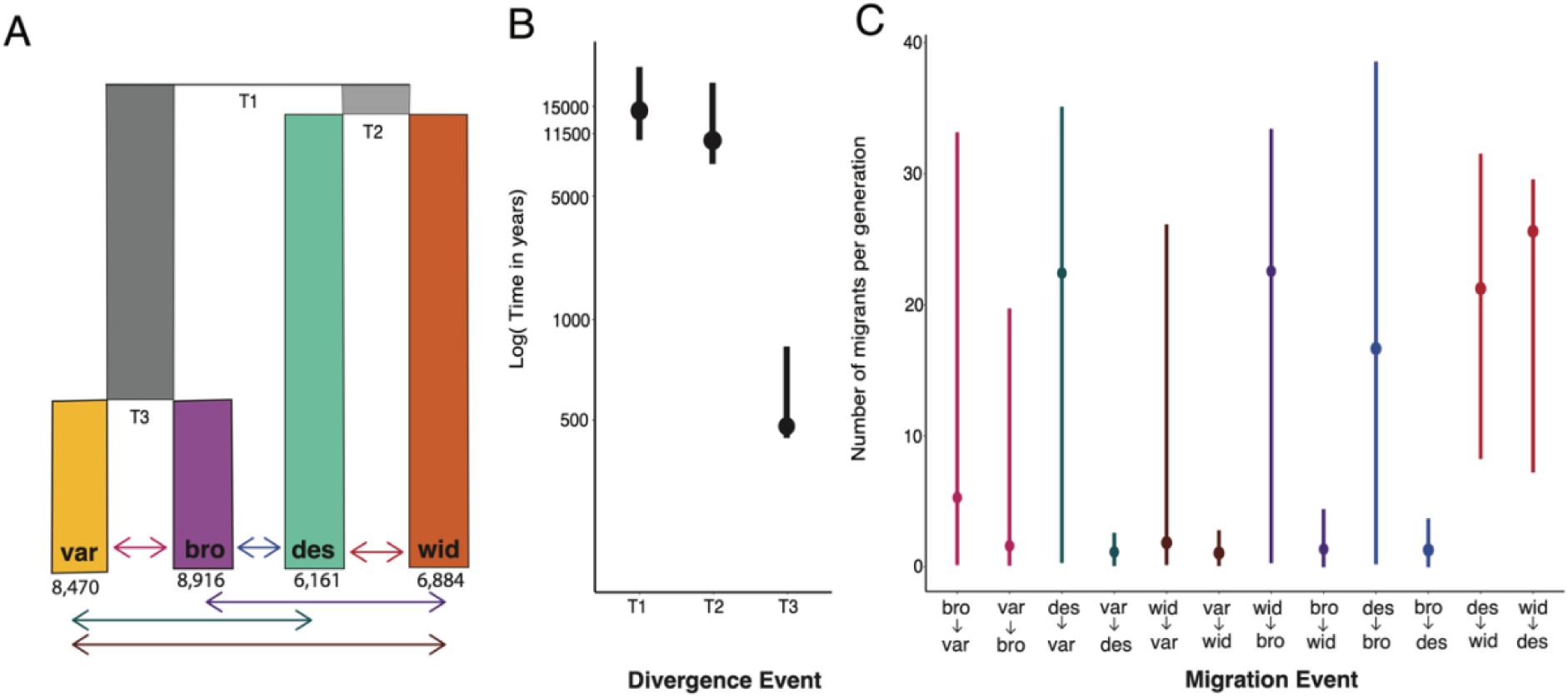
**Demographic history of San Salvador Island radiation**. A) Best supported demographic model in which *C. desquamator* diverged from the *C.* sp. ‘wide-mouth’ based on an LD-pruned dataset with no missing information across all Osprey Lake individuals (67,400 SNPs). Divergence time is shown with the maximum likelihood point estimate from the run with the best fit and the 95% confidence interval for that parameter estimate based on 100 bootstrap replicates. B) Maximum likelihood point estimate and 95% confidence intervals for divergence time based on 100 bootstrap replicates from this model. C) Maximum likelihood point estimate and 95% confidence intervals for migration rate parameters involved in the best fitting model depicted in units of the number of migrants per generation.

**Fig S8.**
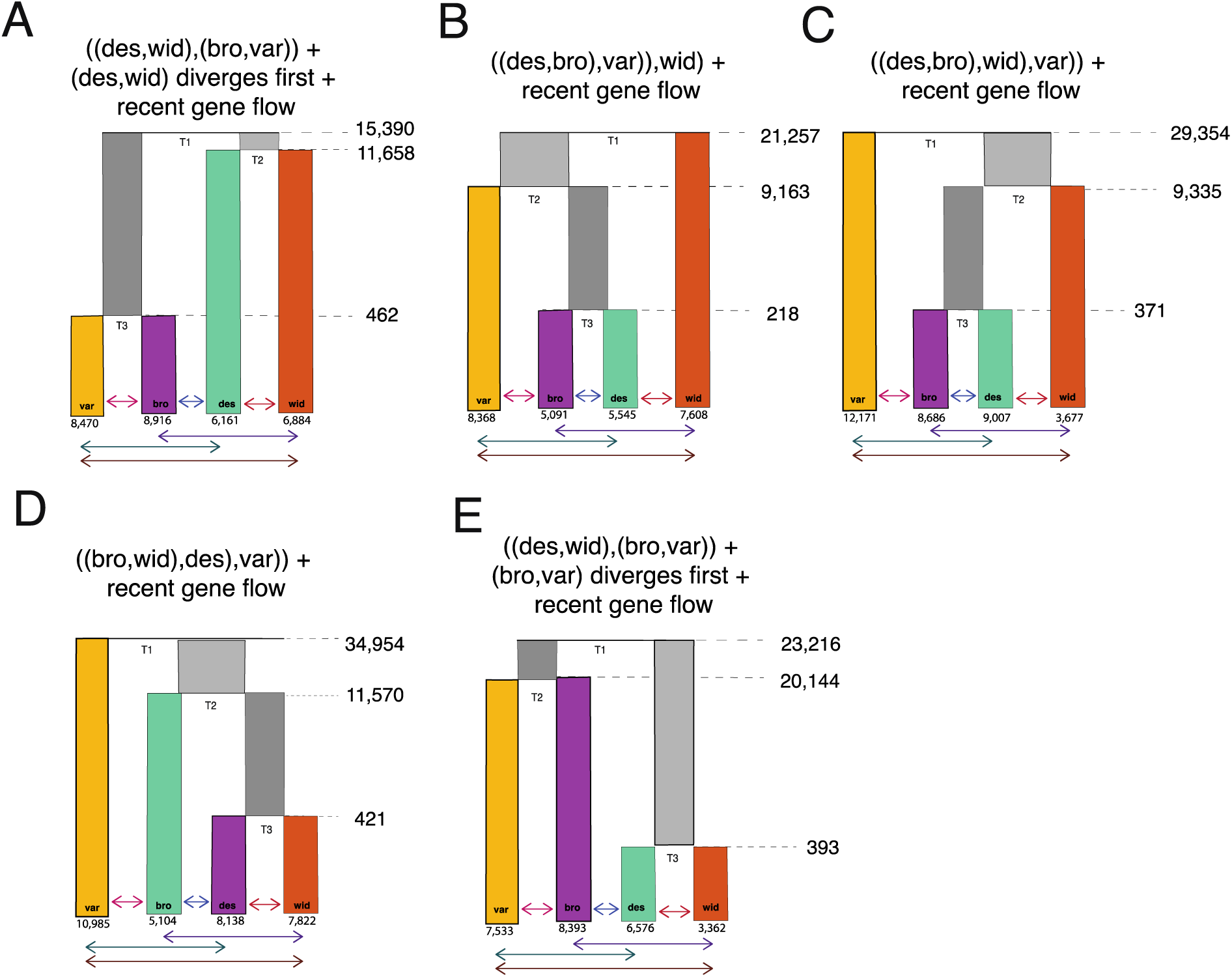
**Top five best fitting demographic models for Osprey Lake.** From a genetic dataset that was LD-pruned and had no missing information across individuals comprised of 67,400 SNPs. Listed in order of lowest AIC scores (Table 1 and S2) with best supported model in A. All top models were of recent bidirectional gene flow but varied in topology among the four species.

**Fig S9.**
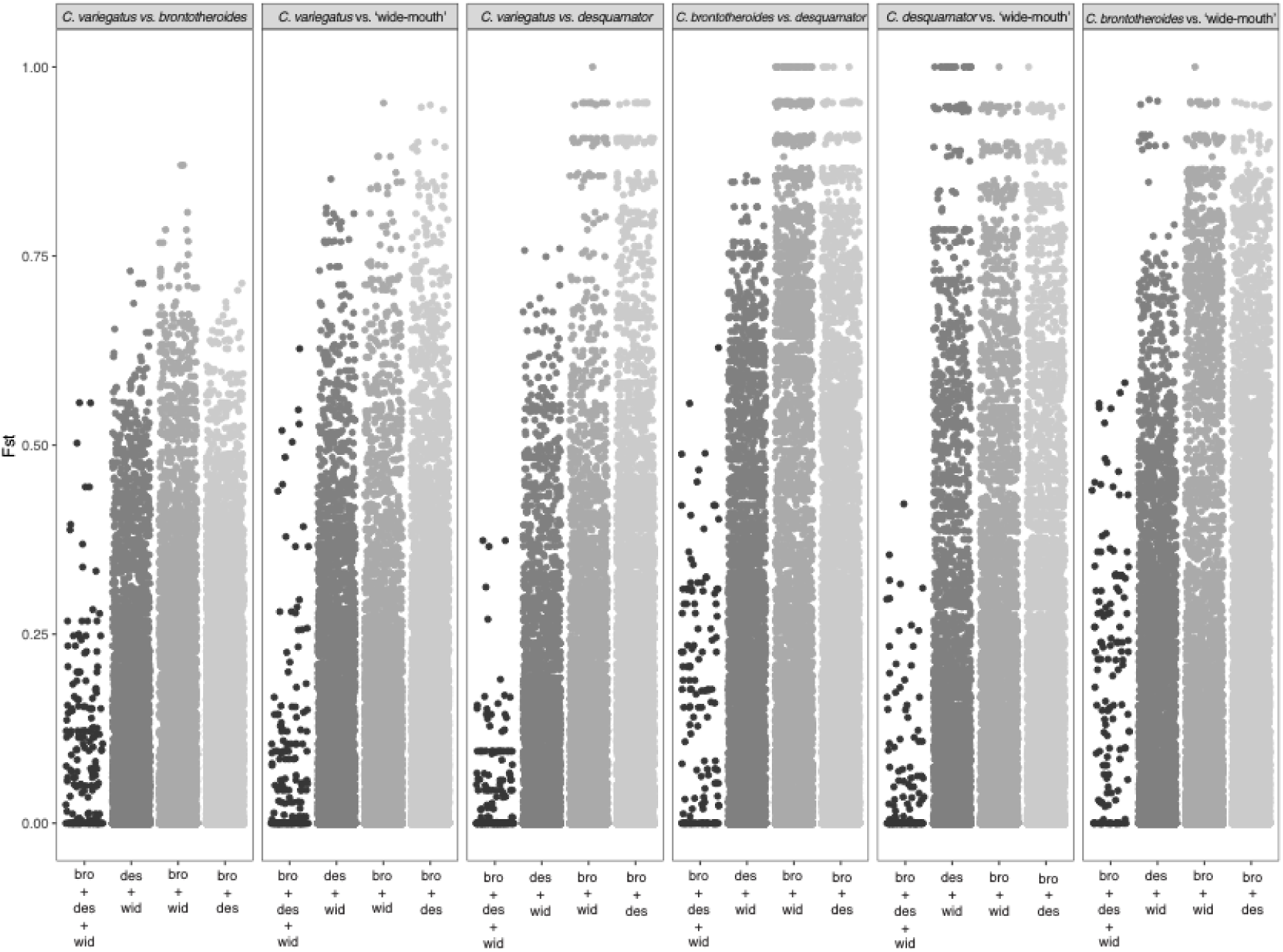
**Genetic divergence among populations in Osprey Lake in regions of shared selection across two or more specialists.** Each panel represents pairwise *F_st_* comparison between two populations for every SNP found in regions under shared selection across two or more specialist species on San Salvador Island: C. brontotheroides (bro); C. desquamator (des), C. sp. ‘wide-mouth’ (wid). Regions shared across all three specialists (bro+des+wid) do not contain highly divergent SNPs between any of the species in the radiation. Highly divergent SNPs (Fst > 0.75) are only found in regions shared between one or two specialists.

**Fig S10.**
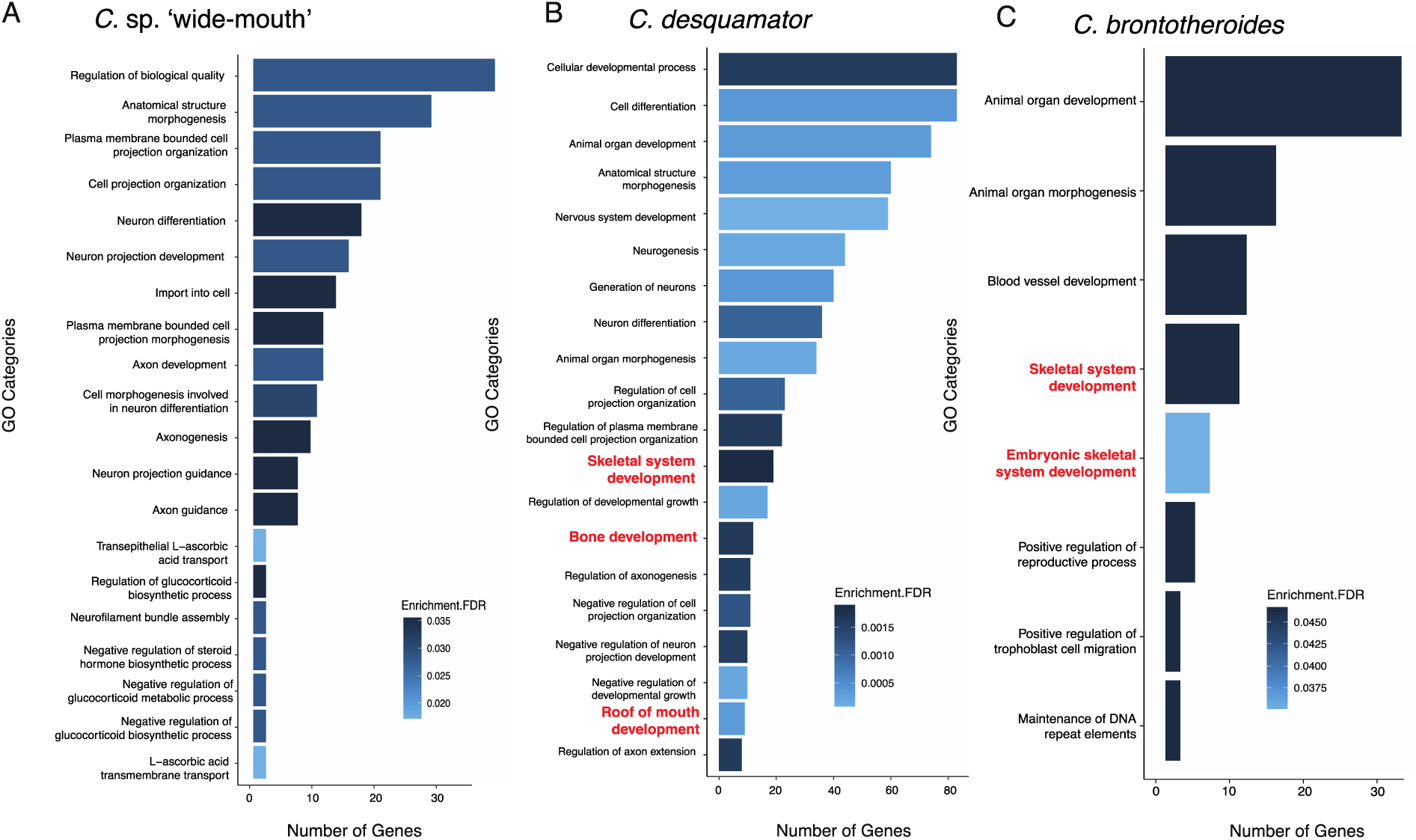
**Top 20 enriched GO categories for divergent alleles in each of the three specialists.** GO enrichment analyses were performed on genes in or near (within 20-kb) of a SNP that was under a hard selective sweep and strongly diverged from generalist species (top 1% of Fst values across genome) for a) ‘wide-mouth’ b) *C. desquamator*, and C) *C. brontotheriodes*. GO categories that were significantly enriched for relevant terms corresponding to craniofacial development, the major axes of morphological divergence in this radiation, are highlighted in red. All terms included were significant at an FDR < 0.05 and full list of terms in Data S1-3.

**Figure S11.**
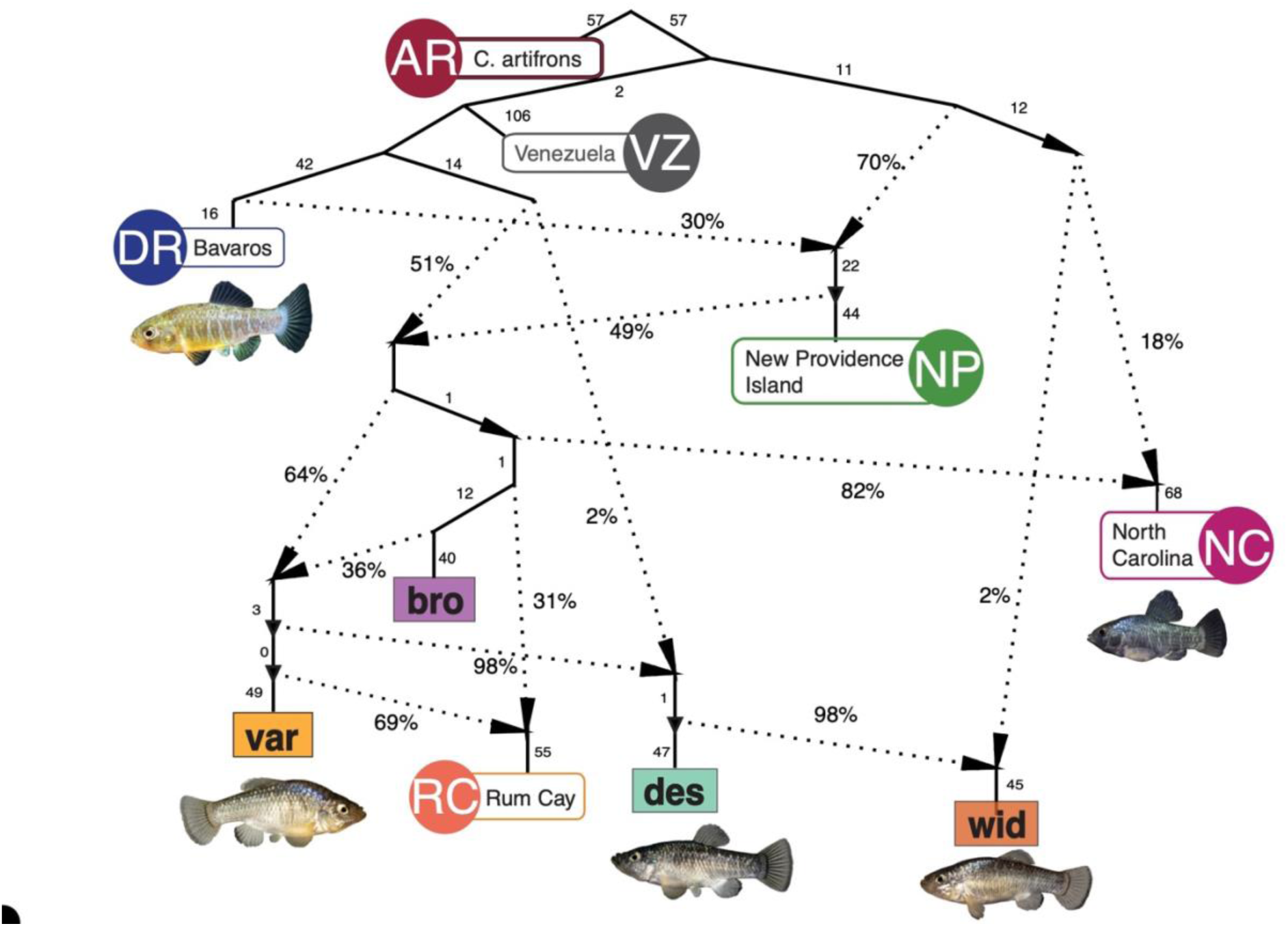
Caribbean pupfish admixture graph. Best-fiting admixture graph from *qpGraph* approach that estimated *f_3_*-statistics across Caribbean populations. This admixture model features 7 admixture events represented by admixture edges and proportions (percentages on dotted branches) and drift edges (numbers on solid branches) that minimize the difference between fitted and estimated *f_3_*-statistics.

**Figure S12.**
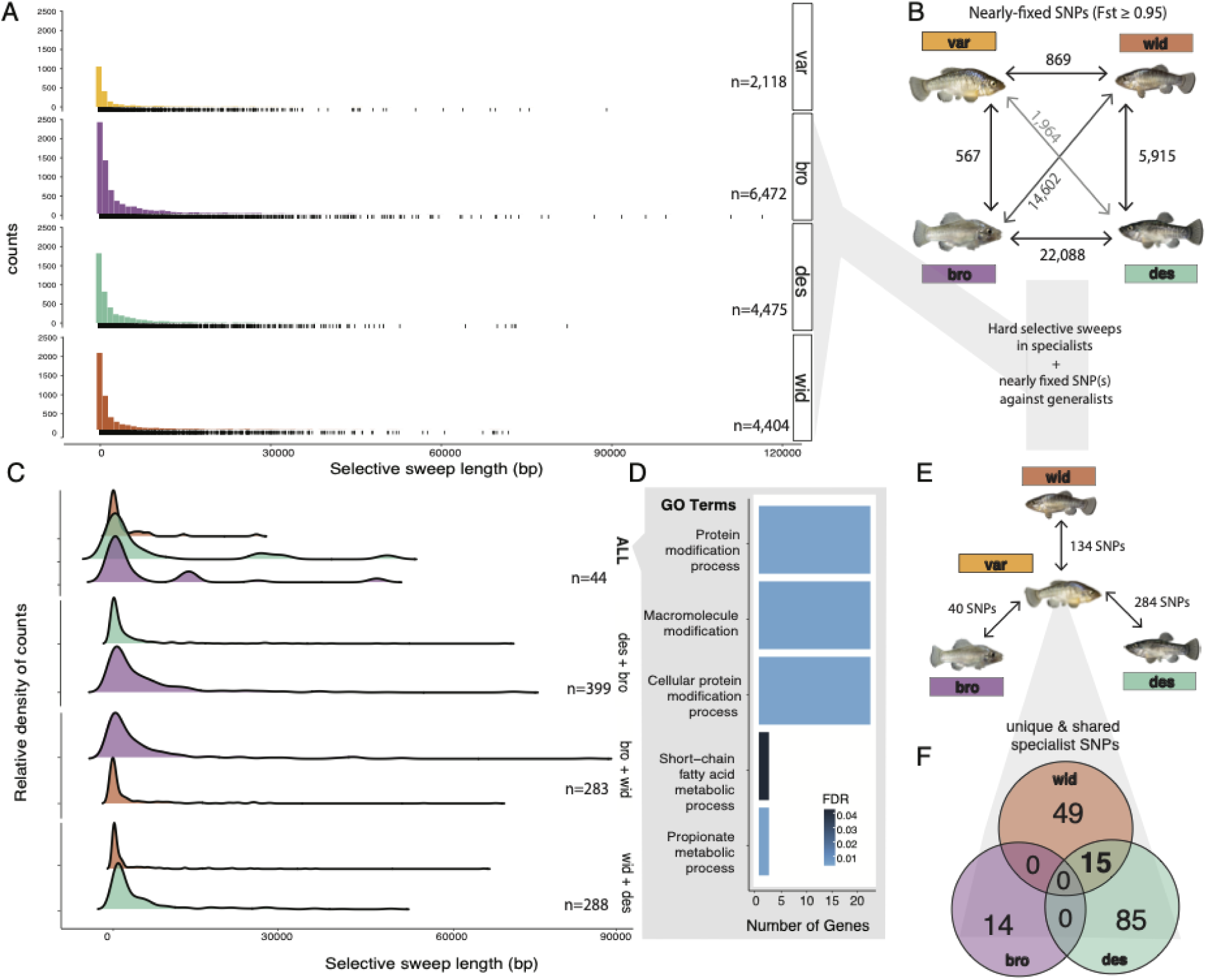
Patterns of selection and genetic divergence in specialist genomes. A) Selective sweep length distributions across all four San Salvador Island species. Rug plot below each histogram represents the counts of selective sweeps in different length bins. B) The total number of fixed or nearly-fixed SNPs (*F_st_*≥ 0.95) between each group in Osprey Pond. C) Ridgeline plots for length distributions of selective sweeps shared between different combinations of specialists (*C. brontotheriodes*: bro, *C. desquamator*: des, *C.* sp. ‘wide-mouth’: wid). D) GO enrichment analysis of the 57 genes contained in the shared selective sweep regions across all specialists. E) The number of adaptive alleles (fixed or nearly-fixed SNPS [*F_st_*≥ 0.95] relative to *C. variegatus* and under selection in each population of specialists in Osprey Lake. F) Venn diagram highlights those adaptive alleles that are unique to each specialist and shared with other specialists.

**Table S1.**
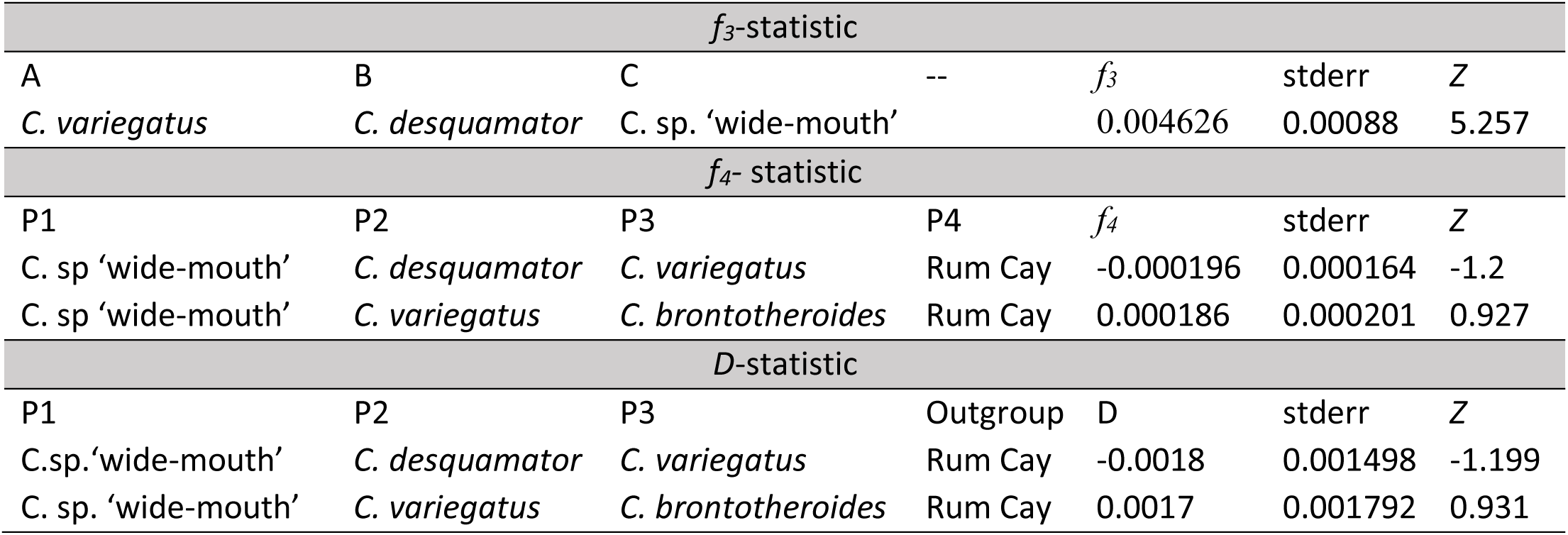
**No significant genome-wide signature of admixture among populations on San Salvador Islands.** Results from three formal tests for introgression to assess whether ‘wide-mouth’ populations are hybrids between generalist *C. variegatus* and scale-eater *C. desquamator*: D-, *f_3_-* and *f_4_-*statistics across all possible combinations of Osprey Lake populations of generalist species. Significant *f_3_-*statistic had Z-scores > -2; significant *f_4_* and D-statistic Z-scores were smaller than -2 and greater than 2. The only significant signature of admixture (bolded Z-scores) comes from D- and *f_4_-*statistics based on relationships that violate the expected tree (((C. desquamator; ‘wide-mouth’),Generalist),Outgroup) and therefore should not be interpreted as evidence of ‘wide-mouth’ being an admixed population. *C. variegatus* population from Rum Cay, the nearest neighbor island in the Bahamas to San Salvador Island was used as an outgroup population for D- and *f_4_-*statistics.

**Table S2.**
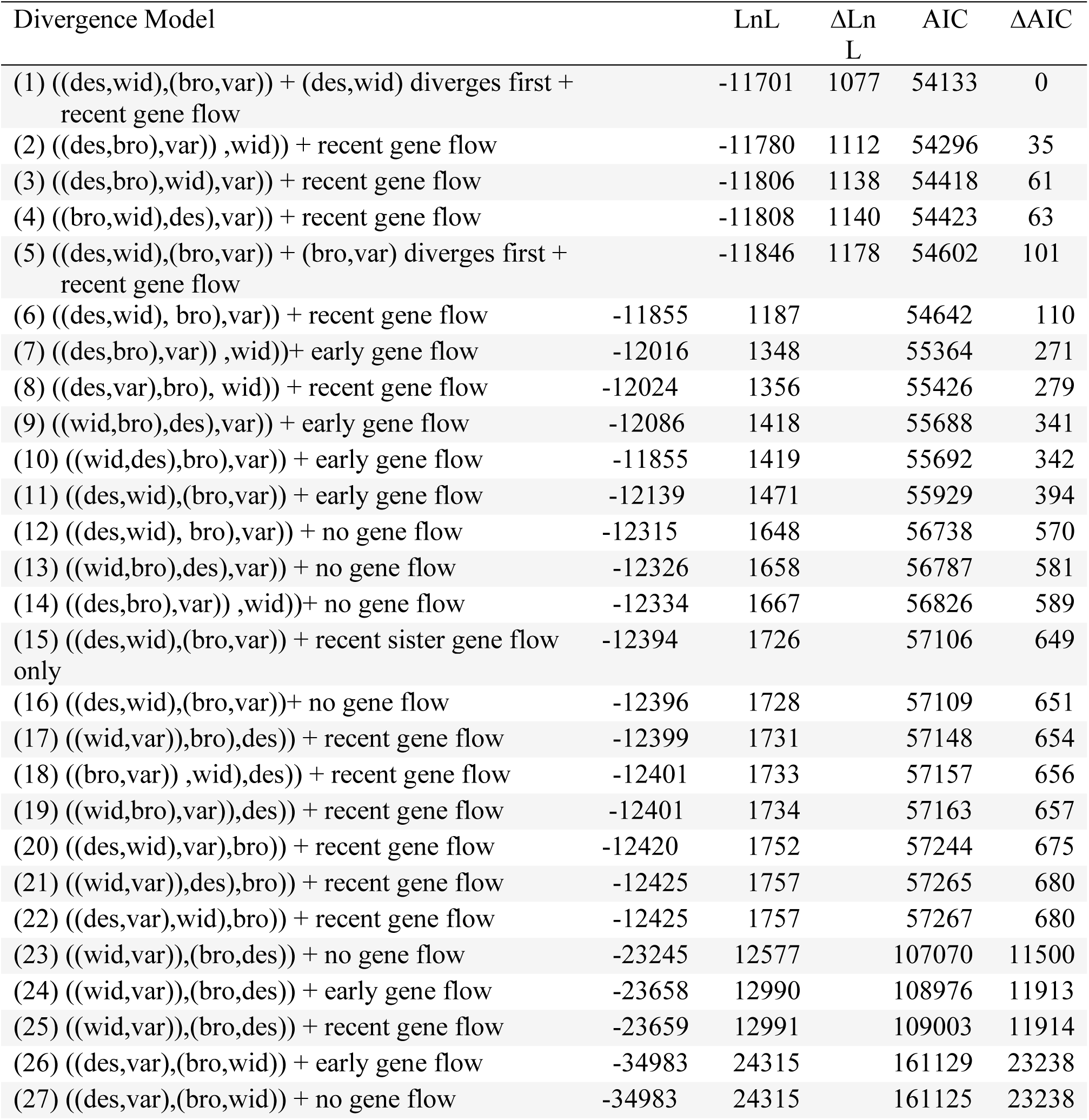
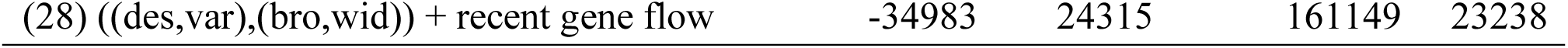
**Support for the 28 demographic models for the evolution of the ‘wide-mouths’ from the site frequency spectrum.** The likelihood and AIC scores all demographic models estimated in *fastsimcoal2* for the Osprey populations of *C. variegatus* var), *C. brontotheroides* (bro), *C. desquamator* (des), and *C.* sp. ‘wide-mouth’ (wid) are presented here with a complete list of all models tested reported in Table S2. Change in likelihood (ΔLnL) represents the difference in likelihood from that of a simulated SFS expected by the demographic model tested. Change in AIC (ΔAIC) represents the difference in AIC scores from that of the model with the smallest ΔLnL. All models presented here represent different divergence scenarios with recent gene flow allowed, which had better support from the data than models with no gene flow or early gene flow. Visual representations of the top five models are depicted in Figure S5.

**Table S3.**
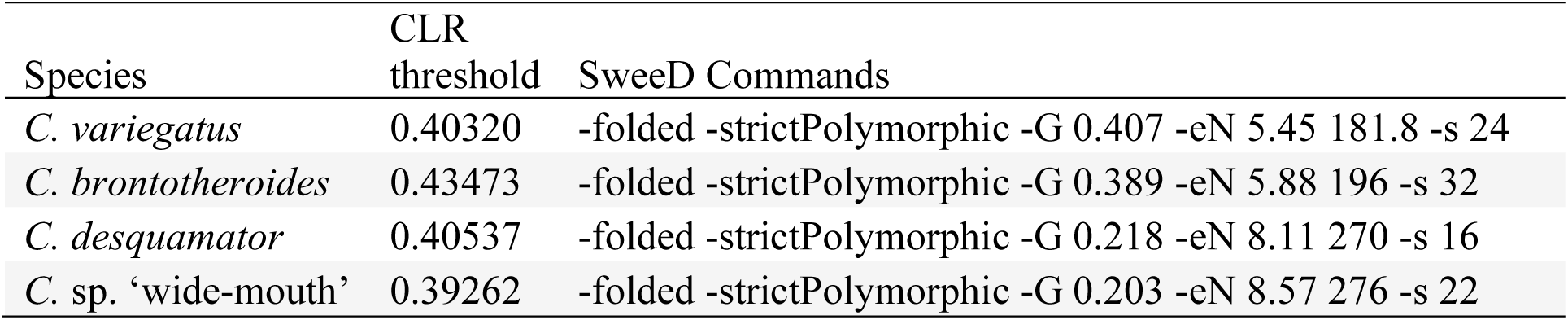
**Parameters for selective sweep analyses in SweeD.** The 95^th^ percentile of composite likelihood ratio threshold based on neutral simulations under the demographic scenario of decreasing population size through time inferred with MSMC (Fig S4), and the population size change and haplotype number information parameters required by SweeD for each species.

| Species | CLR threshold | SweeD Commands |
| --- | --- | --- |
| *C. variegatus* | 0.40320 | -folded -strictPolymorphic -G 0.407 -eN 5.45 181.8 -s 24 |
| *C. brontotheroides* | 0.43473 | -folded -strictPolymorphic -G 0.389 -eN 5.88 196 -s 32 |
| *C. desquamator* | 0.40537 | -folded -strictPolymorphic -G 0.218 -eN 8.11 270 -s 16 |
| *C. sp. 'wide-mouth'* | 0.39262 | -folded -strictPolymorphic -G 0.203 -eN 8.57 276 -s 22 |

**Table S4.** **Genetic divergence among the four populations in Osprey Lake, San Salvador Island across different thresholds of *F_st_*.** Relative measure of genetic divergence was calculated in pairwise combinations of the different species as the number of fixed SNPs between them, the number of fixed or nearly fixed SNPs between them, the top 1% of *F_st_* between them from the distribution of *F_st_* between SNPs and the genome-wide average *F_st_* across all SNPs in the genome.

| $F_{st}$ Comparison | Number of fixed SNPs ($F_{st}=1$) | Number of nearly fixed SNPs ($F_{st} \geq 0.95$) | Top 1% | Genome-wide average |
| --- | --- | --- | --- | --- |
| *C. sp.* ‘wide-mouth’ vs *C. desquamator* | 5,212 | 5,915 | 0.84 | 0.131 |
| *C. sp.* ‘wide-mouth’ vs *C. brontotheroides* | 5238 | 14,602 | 0.85 | 0.15 |
| *C. sp.* ‘wide-mouth’ vs *C. variegatus* | 247 | 869 | 0.68 | 0.104 |
| *C. desquamator* vs *C. brontotheroides* | 6941 | 22,088 | 0.86 | 0.17 |
| *C. desquamator* vs *C. variegatus* | 414 | 1,964 | 0.71 | 0.116 |
| *C. variegatus* vs *C. brontotheroides* | 110 | 567 | 0.637 | 0.09 |

**Table S5.** **Comparison of the average absolute genetic divergence (*D_xy_*) between the four species in Osprey Lake ‘wide-mouth’, *C. variegatus, C. desquamator and C. brontotheriodes* and the shared adaptive alleles between scale-eater populations.**

| Comparison | Mean $D_{xy}$ |
| --- | --- |
| *C. variegatus* vs *C. brontotheroides* | 0.167 |
| *C. variegatus* vs *C. desquamator* | 0.169 |
| *C. variegatus* vs *C. sp.* ‘wide-mouth’ | 0.165 |
| *C. brontotheroides* vs *C. desquamator* | 0.162 |
| *C. brontotheroides* vs *C. sp.* ‘wide-mouth’ | 0.156 |
| *C. desquamator* vs *C. sp.* ‘wide-mouth’ | 0.157 |
| Selective sweeps in *C. sp.* ‘wide-mouth’ | 0.123 |
| Shared genetic variants with *C. desquamator* | 0.0814 |
| Unique genetic variants in *C. sp.* ‘wide-mouth’ | 0.24 |

**Table S6.**
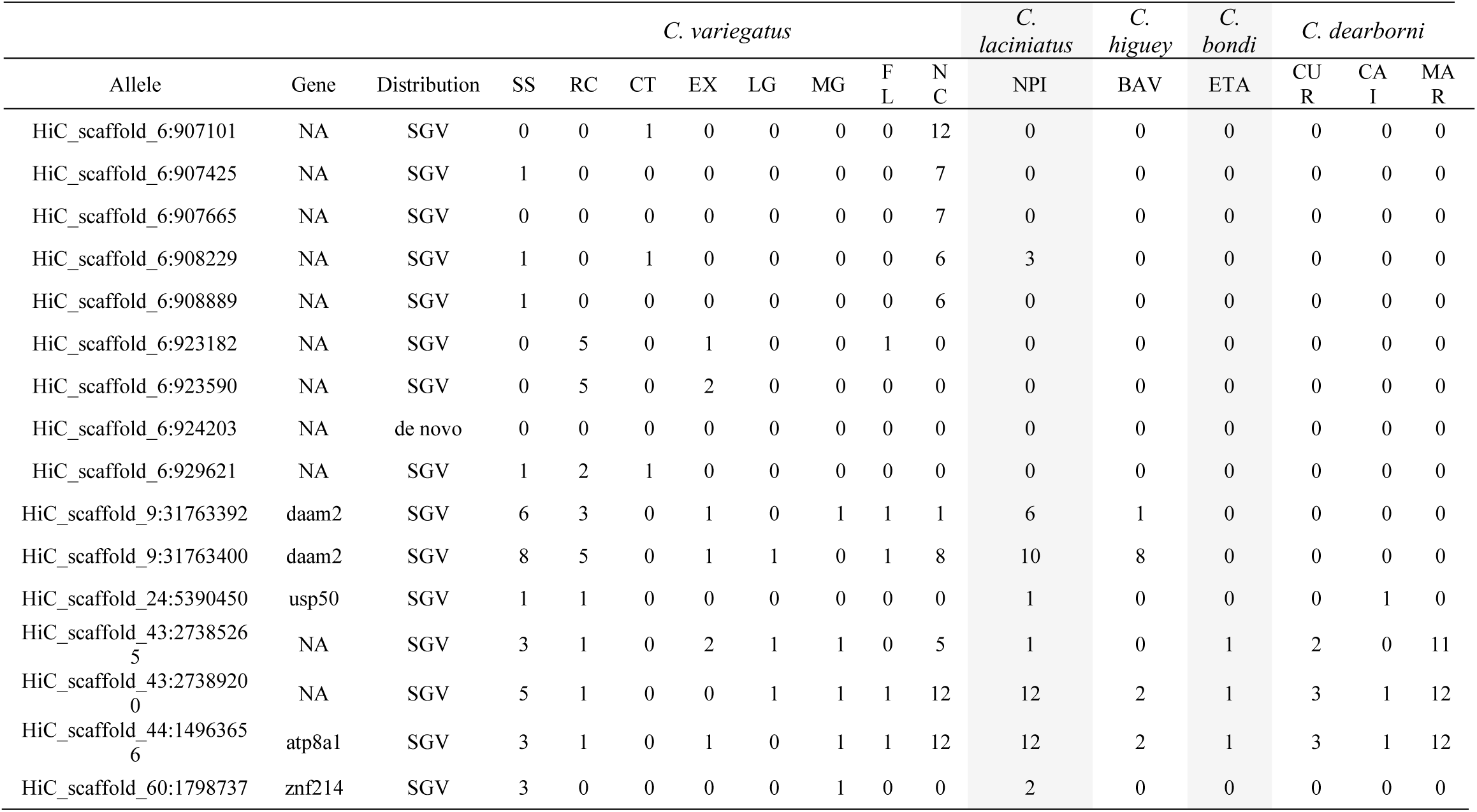
**The distribution of shared adaptive alleles between the two scale-eating species across the Caribbean.** Numbers represent the copies of the *C. desquamator* and ‘wide-mouth’ allele present in outgroup populations collected from San Salvador Island (SS), Rum Cay (RC), Cat Island (CT), Exumas Islands (EX), Long Island (LG), Mayaguana (MG), New Providence Island (NPI) in the Bahamas. Lagunas Bavaros (BAV) and Etang Saumautre (ETA) in Dominican Republic, Sarasota Estuary in Florida (FL), Curacoa (CUR), Caicos Island (CAI) and Isla Margarita (MAR) in Venezuela.

**Table S7.**
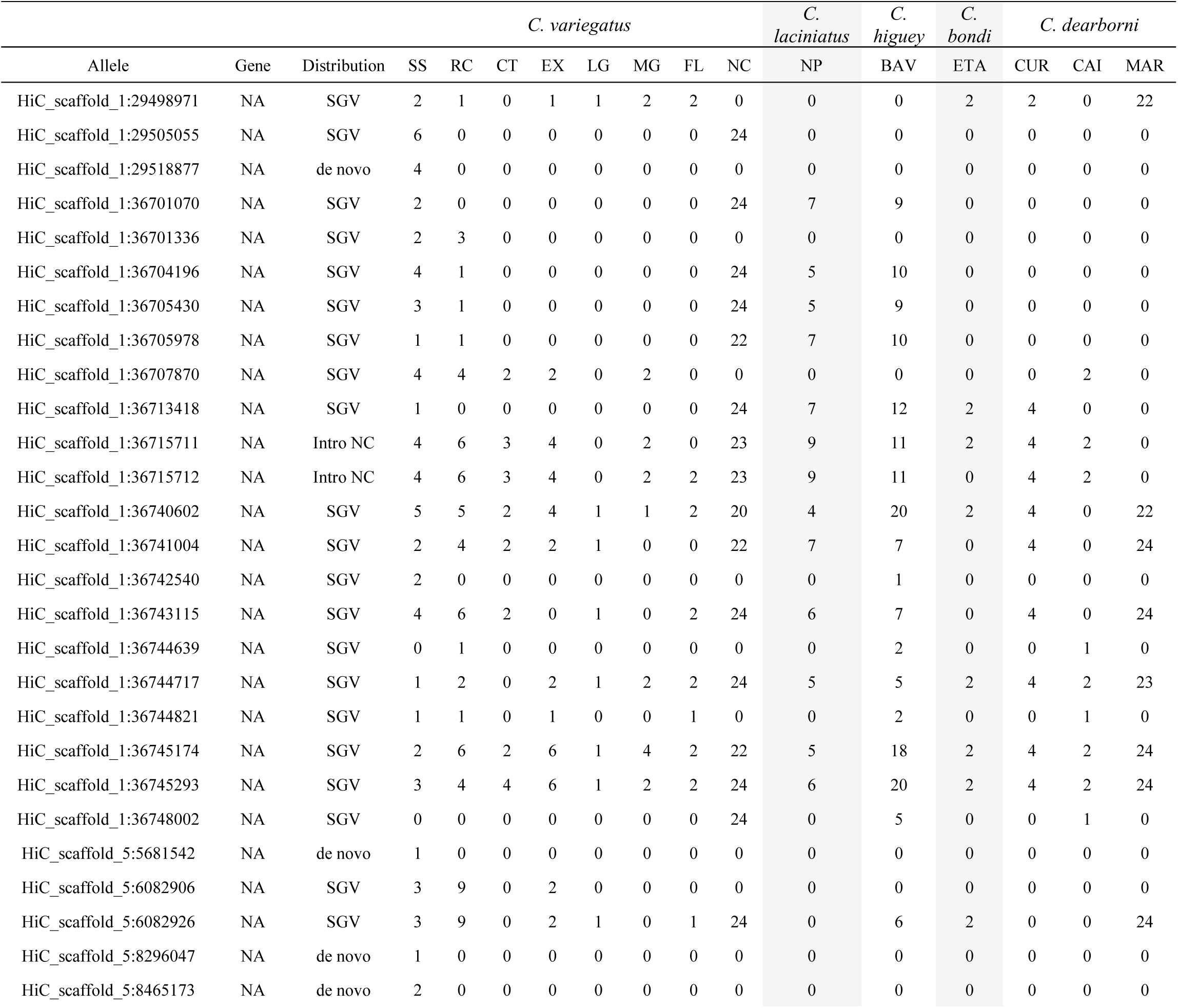
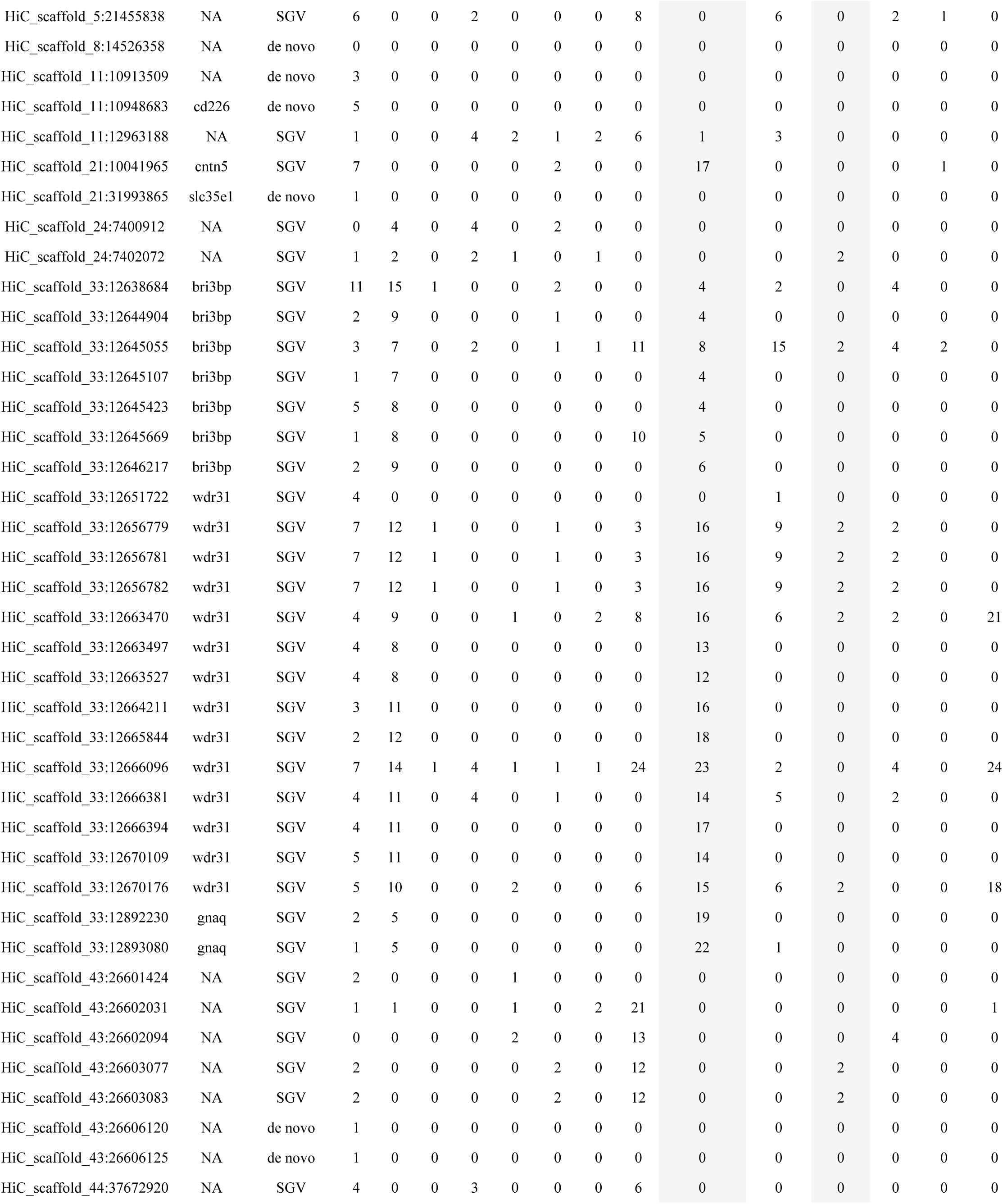
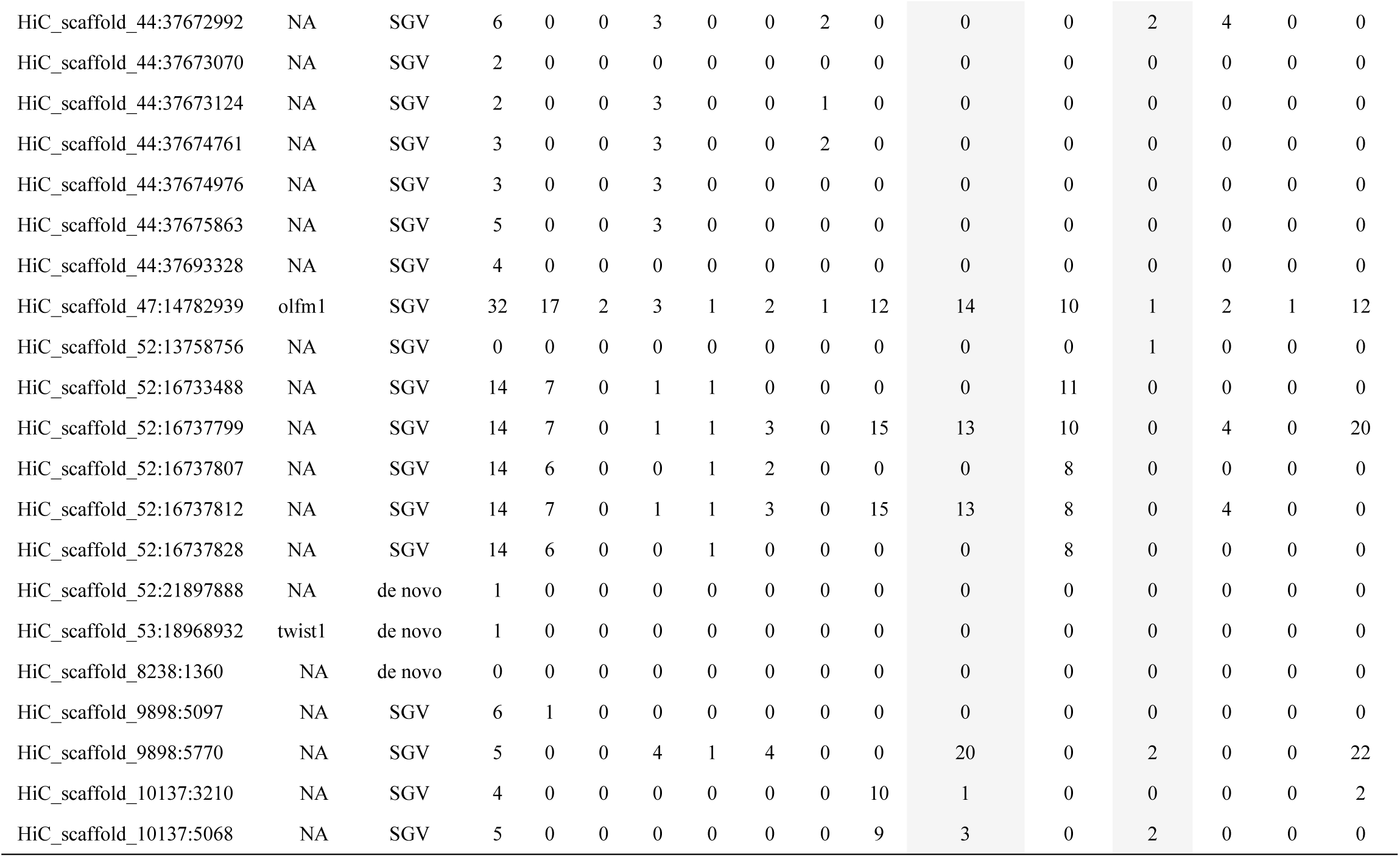
**The distribution of unique adaptive alleles to the highly specialized scale-eater *C. desquamator* across the Caribbean.** Numbers represent the copies of the *C. desquamator* allele present in outgroup populations collected from San Salvador Island (SS), Rum Cay (RC), Cat Island (CT), Exumas Islands (EX), Long Island (LG), Mayaguana (MG), New Providence Island (NPI) in the Bahamas. Lagunas Bavaros (BAV) and Etang Saumautre (ETA) in Dominican Republic, Sarasota Estuary in Florida (FL), Curacoa (CUR), Caicos Island (CAI) and Isla Margarita (MAR) in Venezuela. The spatial distribution of each allele is summarized into three categories: standing genetic variation (SGV); de novo to San Salvador Island populations (de novo) and alleles found in introgressed regions (Intro).

**Table S8.** **The distribution of unique adaptive alleles to the intermediate scale-eater *C. sp.* ‘wide-mouth’ across the Caribbean.** Numbers represent the copies of the *C. desquamator* allele present in outgroup populations collected from San Salvador Island (SS), Rum Cay (RC), Cat Island (CT), Exumas Islands (EX), Long Island (LG), Mayaguana (MG), New Providence Island (NPI) in the Bahamas. Lagunas Bavaros (BAV) and Etang Saumautre (ETA) in Dominican Republic, Sarasota Estuary in Florida (FL), Curacoa (CUR), Caicos Island (CAI) and Isla Margarita (MAR) in Venezuela. The spatial distribution of each allele is summarized into three categories: standing genetic variation (SGV); de novo to San Salvador Island populations (de novo) and alleles found in introgressed regions (Intro). Putative donor populations for introgressed regions are indicated by the same location codes.

| *C. variegatus* |  |  |  |  |  |  |  |  |  |  | *C.*<br>*laciniatus* | *C.*<br>*higuey* | *C.*<br>*bondi* | *C.*<br>*dearborni* |  |  |
| --- | --- | --- | --- | --- | --- | --- | --- | --- | --- | --- | --- | --- | --- | --- | --- | --- |
| Allele | Gene | Distribution | SS | RC | CT | EX | LG | MG | FL | NC | NP | BAV | ETA | CUR | CAI | MAR |
| HiC_scaffold_1:34344664 | smarce1 | Intro NC | 1 | 0 | 0 | 4 | 1 | 2 | 2 | 24 | 2 | 11 | 2 | 4 | 0 | 20 |
| HiC_scaffold_1:34346071 | smarce1 | Intro NC | 1 | 0 | 0 | 0 | 0 | 1 | 0 | 24 | 0 | 0 | 0 | 0 | 0 | 0 |
| HiC_scaffold_1:34346073 | smarce1 | Intro NC | 1 | 0 | 0 | 0 | 0 | 1 | 0 | 24 | 0 | 0 | 0 | 0 | 0 | 0 |
| HiC_scaffold_5:20660128 | NA | SGV | 1 | 0 | 0 | 0 | 0 | 0 | 0 | 0 | 0 | 0 | 0 | 0 | 0 | 0 |
| HiC_scaffold_5:20941055 | NA | SGV | 1 | 1 | 0 | 3 | 0 | 2 | 0 | 0 | 5 | 6 | 0 | 0 | 1 | 0 |
| HiC_scaffold_5:20941063 | NA | SGV | 1 | 1 | 0 | 3 | 0 | 2 | 0 | 0 | 5 | 6 | 0 | 0 | 1 | 0 |
| HiC_scaffold_5:20945163 | NA | SGV | 3 | 1 | 0 | 3 | 0 | 2 | 0 | 0 | 8 | 11 | 0 | 0 | 1 | 2 |
| HiC_scaffold_5:20945417 | NA | SGV | 1 | 1 | 0 | 4 | 0 | 2 | 0 | 0 | 5 | 7 | 0 | 0 | 1 | 0 |
| HiC_scaffold_5:20945569 | NA | SGV | 1 | 1 | 0 | 3 | 0 | 3 | 0 | 0 | 4 | 9 | 0 | 4 | 2 | 0 |
| HiC_scaffold_5:20964282 | NA | SGV | 1 | 1 | 0 | 2 | 1 | 2 | 0 | 15 | 5 | 9 | 0 | 0 | 0 | 0 |
| HiC_scaffold_5:20964283 | NA | SGV | 1 | 1 | 0 | 2 | 1 | 2 | 0 | 15 | 5 | 9 | 0 | 0 | 0 | 0 |
| HiC_scaffold_5:20968573 | NA | SGV | 0 | 1 | 0 | 4 | 1 | 3 | 0 | 0 | 7 | 7 | 0 | 4 | 0 | 0 |
| HiC_scaffold_5:20969017 | NA | SGV | 1 | 1 | 0 | 4 | 1 | 4 | 0 | 0 | 7 | 7 | 0 | 0 | 0 | 0 |
| HiC_scaffold_5:20969195 | NA | SGV | 2 | 1 | 0 | 2 | 1 | 2 | 0 | 0 | 7 | 6 | 0 | 0 | 0 | 0 |
| HiC_scaffold_5:20969509 | NA | SGV | 0 | 1 | 0 | 3 | 1 | 2 | 0 | 16 | 6 | 6 | 0 | 4 | 0 | 0 |
| HiC_scaffold_5:20969704 | NA | SGV | 1 | 2 | 0 | 3 | 1 | 4 | 0 | 0 | 6 | 6 | 0 | 0 | 0 | 0 |
| HiC_scaffold_5:20969759 | NA | SGV | 1 | 1 | 0 | 3 | 1 | 2 | 0 | 10 | 5 | 5 | 0 | 4 | 0 | 0 |
| HiC_scaffold_5:20970496 | NA | SGV | 0 | 2 | 0 | 3 | 0 | 2 | 0 | 14 | 6 | 6 | 0 | 4 | 1 | 0 |
| HiC_scaffold_5:20971701 | NA | SGV | 1 | 1 | 0 | 3 | 1 | 4 | 0 | 0 | 8 | 5 | 0 | 0 | 0 | 1 |
| HiC_scaffold_5:20973584 | NA | SGV | 1 | 2 | 0 | 4 | 1 | 4 | 0 | 0 | 9 | 6 | 0 | 0 | 0 | 0 |
| HiC_scaffold_5:20974017 | NA | SGV | 0 | 1 | 0 | 1 | 0 | 4 | 2 | 3 | 8 | 6 | 0 | 0 | 0 | 0 |
| HiC_scaffold_5:20977516 | NA | SGV | 1 | 2 | 0 | 2 | 1 | 4 | 0 | 0 | 4 | 7 | 0 | 0 | 0 | 0 |
| HiC_scaffold_5:20979898 | NA | SGV | 0 | 2 | 2 | 3 | 1 | 2 | 1 | 0 | 6 | 5 | 0 | 0 | 1 | 0 |
| HiC_scaffold_5:20986384 | NA | SGV | 0 | 2 | 0 | 2 | 1 | 2 | 0 | 0 | 8 | 0 | 0 | 0 | 2 | 0 |
| HiC_scaffold_5:20988210 | NA | SGV | 0 | 3 | 0 | 2 | 1 | 2 | 0 | 0 | 7 | 0 | 0 | 0 | 0 | 0 |
| HiC_scaffold_5:20988292 | NA | SGV | 0 | 4 | 0 | 3 | 1 | 4 | 0 | 0 | 6 | 1 | 0 | 0 | 2 | 0 |
| HiC_scaffold_5:20988346 | NA | SGV | 0 | 5 | 0 | 3 | 0 | 2 | 0 | 0 | 8 | 0 | 0 | 0 | 1 | 0 |
| HiC_scaffold_11:21559293 | NA | SGV | 1 | 22 | 0 | 4 | 0 | 2 | 1 | 0 | 13 | 2 | 0 | 0 | 0 | 0 |
| HiC_scaffold_11:21571679 | NA | SGV | 1 | 17 | 0 | 1 | 1 | 2 | 1 | 5 | 13 | 2 | 0 | 0 | 1 | 0 |
| HiC_scaffold_11:21571720 | NA | SGV | 1 | 22 | 0 | 1 | 1 | 0 | 0 | 5 | 0 | 3 | 0 | 0 | 0 | 0 |
| HiC_scaffold_11:21571742 | NA | SGV | 1 | 22 | 0 | 1 | 1 | 0 | 0 | 5 | 0 | 3 | 0 | 0 | 1 | 0 |
| HiC_scaffold_24:15255378 | NA | Intro NC | 0 | 0 | 0 | 0 | 1 | 0 | 1 | 23 | 0 | 0 | 0 | 0 | 0 | 24 |
| HiC_scaffold_43:19331844 | semag4 | SGV | 7 | 0 | 2 | 0 | 0 | 0 | 0 | 0 | 0 | 0 | 0 | 0 | 0 | 0 |
| HiC_scaffold_44:14922408 | NA | SGV | 4 | 11 | 1 | 0 | 0 | 0 | 0 | 0 | 0 | 1 | 1 | 0 | 0 | 0 |
| HiC_scaffold_44:14922640 | NA | SGV | 2 | 9 | 0 | 0 | 0 | 0 | 0 | 0 | 0 | 0 | 2 | 0 | 2 | 0 |
| HiC_scaffold_44:14922976 | NA | SGV | 2 | 11 | 0 | 0 | 0 | 0 | 0 | 0 | 3 | 1 | 0 | 4 | 2 | 0 |
| HiC_scaffold_52:9589311 | slitrk5 | Intro BAV | 1 | 0 | 0 | 2 | 1 | 0 | 0 | 21 | 7 | 20 | 0 | 0 | 0 | 20 |
| HiC_scaffold_52:9589319 | slitrk5 | Intro BAV | 1 | 0 | 0 | 2 | 1 | 0 | 0 | 21 | 7 | 20 | 0 | 0 | 0 | 20 |
| HiC_scaffold_52:9592382 | slitrk5 | Intro BAV | 0 | 1 | 0 | 0 | 2 | 0 | 0 | 2 | 1 | 1 | 2 | 0 | 0 | 26 |
| HiC_scaffold_5743:1358 | NA | SGV | 0 | 4 | 0 | 2 | 1 | 2 | 0 | 0 | 9 | 0 | 0 | 0 | 2 | 0 |
| HiC_scaffold_5789:9634 | NA | SGV | 0 | 3 | 0 | 2 | 0 | 2 | 0 | 0 | 7 | 0 | 0 | 0 | 0 | 0 |
| HiC_scaffold_5789:10010 | NA | de novo | 0 | 0 | 0 | 0 | 0 | 0 | 0 | 0 | 0 | 0 | 0 | 0 | 0 | 0 |
| HiC_scaffold_5789:11490 | NA | SGV | 0 | 4 | 0 | 2 | 2 | 2 | 0 | 0 | 5 | 0 | 0 | 0 | 0 | 0 |
| HiC_scaffold_5789:12093 | NA | SGV | 0 | 4 | 0 | 2 | 0 | 2 | 0 | 0 | 7 | 0 | 0 | 0 | 1 | 0 |
| HiC_scaffold_13976:3987 | NA | SGV | 0 | 0 | 0 | 2 | 1 | 2 | 0 | 0 | 5 | 0 | 0 | 0 | 2 | 0 |

**Table S10.** **Assessment of introgression signatures between population in Osprey Lake, San Salvador Island in regions that contain putative adaptive alleles unique to *C. sp*. wide-mouth, unique to *C. desquamator* and shared between both scale-eating populations.** Introgressed regions are highlighted in bold. Introgressed regions were those that had a relative node depth (RND) value between two populations. RND statistics were calculated in 10-kb windows between *C. desquamator* and *C.* sp. ‘wide-mouth’ (des vs wid) and *C. desquamator* and *C.* sp. ‘wide-mouth’ (des vs var), with the latter test serving as a control for recent and ongoing gene exchange due to sympatric overlap in the lake that is not relevant for scale-eating adaptation. Adaptive allele, gene names, and RND values that are bolded represent candidate introgressed regions with adaptive alleles in them. Candidate introgressed regions were regions that fell below a significance threshold value (RND < 0.28) based on the lower 5^th^ percentile of RND values calculated from simulations of two populations that experience no gene flow during divergence. The number of exons present in each 10-kb window is also included in this table.

| Adaptive allele | Gene | RND<br>des vs wid | RND<br>des v var | Introgressed Region | Number of<br>exons |
| --- | --- | --- | --- | --- | --- |
| Unique *C. sp.* 'wide-mouth' adaptive alleles |  |  |  |  |  |
| HiC_scaffold_5:20941063 | NA | 1.48663438 | 1.11402561 | 20940001-20950000 | 0 |
| HiC_scaffold_5:20964282 | NA | 1.53346729 | 1.28660246 | 20960001-20970000 | 0 |
| HiC_scaffold_5:20970496 | NA | 1.52647461 | 1.29854143 | 20970001-20980000 | 0 |
| HiC_scaffold_5:20986384 | NA | 1.34806367 | 0.90675156 | 20980001-20990000 | 0 |
| HiC_scaffold_11:21559293 | NA | 1.24490382 | 0.78312989 | 21550001-21560000 | 0 |
| HiC_scaffold_11:21571679 | NA | 1.74507532 | 0.87626263 | 21570001-21580000 | 0 |
| HiC_scaffold_24:15255378 | NA | 0.60467498 | 0.54123148 | 15250001-15260000 | 0 |
| HiC_scaffold_43:19331844 | Sema4g | 1.18018089 | 0.94111453 | 19330001-19340000 | 0 |
| HiC_scaffold_44:14922408 | atp8a1 | 1.60947983 | 0.97866515 | 14920001-14930000 | 0 |
| HiC_scaffold_52:9589311 | Slitrk5 | 1.47013193 | 1.14495019 | 9580001-9590000 | 0 |
| *C. desquamator* and *C. sp.* 'wide-mouth' adaptive alleles |  |  |  |  |  |
| **HiC_scaffold_6:907101** | **NA** | **0.01764957** | 0.304609 | 900001-910000 | 0 |
| **HiC_scaffold_6:923182** | **NA** | **0.03560806** | 0.41432253 | 920001-930000 | 0 |
| **HiC_scaffold_6:934123** | **NA** | **0.01180417** | 0.29362216 | 930001-940000 | 0 |
| **HiC_scaffold_6:941221** | **NA** | **0.03304484** | 0.52001902 | 940001-950000 | 0 |
| HiC_scaffold_6:951028 | NA | 0.33613727 | 0.34973512 | 950001-960000 | **0** |
| **HiC_scaffold_9:31763392** | **DAAM2** | **0.0241461** | 0.07809267 | 31750001-31760000 | 4 |
| **HiC_scaffold_9:31763392** | **DAAM2** | **0.06961605** | 0.83262139 | 31760001-31770000 | 5 |
| **HiC_scaffold_24:5390450** | **Usp50** | **0.0833701** | 0.90291628 | 5390001-5400000 | 0 |
| HiC_scaffold_43:27385265 | NA | 1.30551148 | 2.06180406 | 27380001-27390000 | 0 |
| **HiC_scaffold_44:14963656** | **ATP8A1** | **0.13299985** | 1.26725109 | 14960001-14970000 | 2 |
| **HiC_scaffold_60:1798737** | **ZNF214** | **0.08551992** | 1.60256069 | 1790001-1800000 | 1 |
| Unique *C. desquamator* adaptive alleles |  |  |  |  |  |
| HiC_scaffold_5:6082906 | NA | 1.61515973 | 1.39739811 | 6080001-6090000 | 0 |
| HiC_scaffold_11:10913509 | cd226 | 0.25420561 | 1.04030645 | 10910001-10920000 | 2 |
| HiC_scaffold_11:10948683 | cd226 | 1.11058873 | 0.75472269 | 10940001-10950000 | 2 |
| HiC_scaffold_21:10041965 | cntn5 | 1.51795767 | 1.5514369 | 10040001-10050000 | 3 |
| HiC_scaffold_21:31993865 | SLC35E1 | 4.35 | 4.81944444 | 31990001-32000000 | 3 |
| HiC_scaffold_24:7401348 | NA | 0.62028279 | 0.62001004 | 7400001-7410000 | 0 |
| HiC_scaffold_33:12638684 | bri3bp | 1.87610126 | 1.80369995 | 12630001-12640000 | 1 |
| HiC_scaffold_33:12641481 | bri3bp | 2.05234267 | 1.93074547 | 12640001-12650000 | 2 |
| HiC_scaffold_33:12656933 | wdr31 | 2.08801498 | 1.91525971 | 12650001-12660000 | 8 |
| HiC_scaffold_33:12664211 | wdr31 | 2.04327131 | 1.88029401 | 12660001-12670000 | 0 |
| HiC_scaffold_33:12670176 | wdr31 | 2 | 1.87301587 | 12670001-12680000 | 0 |
| HiC_scaffold_33:12891903 | gnaq | 1.19586694 | 1.21312932 | 12890001-12900000 | 0 |
| HiC_scaffold_43:26606125 | NA | 1.7651081 | 1.60827345 | 26600001-26610000 | 0 |
| HiC_scaffold_44:37673070 | NA | 1.3785441 | 1.29438333 | 37670001-37680000 | 0 |
| HiC_scaffold_44:37693328 | NA | 1.2303647 | 1.09117949 | 37690001-37700000 | 0 |
| **HiC_scaffold_47:14782939** | **olfm1** | **0.19804878** | 0.7647762 | 14780001-14790000 | 1 |
| HiC_scaffold_52:16737828 | NA | 1.19630072 | 1.25855174 | 16730001-16740000 | 0 |
| **HiC_scaffold_53:18968932** | **twist1** | **0.05856256** | 0.54868019 | 18960001-18970000 | 2 |

**Table S11.** **Genome-wide signatures of differential introgression between *C.* sp. ‘wide-mouth’ and *C. desquamator*.** *f_4_*-statistics, standard error, z-scores and p-values for genome-wide test with 5 outgroup generalist populations from across the Caribbean. *C. artifrons* used as outgroup species in which we expect minimal gene flow.

| P1 | P2 | P3 | Outgroup | *f*<sub>4</sub> | Standard Error | Z-score | P-value |
| --- | --- | --- | --- | --- | --- | --- | --- |
| **wid** | des | Dominican Republic | *Artifrons* | -0.0003 | 0.000173 | -1.7136 | 0.082 |
| **wid** | des | Rum Cay | *Artifrons* | 0.00035 | 0.000188 | 1.90484 | 0.05 |
| **wid** | des | North Carolina | *Artifrons* | 0.00053 | 0.000186 | 2.85 | 0.0044 |
| **wid** | des | Venezuela | *Artifrons* | 0.00011 | 0.000143 | 0.78891 | 0.43 |
| **wid** | des | New Providence Island | *Artifrons* | 0.00027 | 0.000152 | 1.81370 | 0.069 |

## References

1. Dieckmann U, Doebeli M. 1999 On the origin of species by sympatric speciation. Nature 400, 354.

2. Polechová J, Barton NH, Gavrilefs S. 2005 Speciation thrrough competition: a critical review. Evolution (N. Y). 59, 1194–1210. (doi:10.1554/04-691)

3. Bolnick DI. 2006 Multi-species outcomes in a common model of sympatric speciation. J. Theor. Biol. 241, 734–744. (doi:10.1016/j.jtbi.2006.01.009)

4. Kondrashov AS, Kondrashov FA. 1999 Interactions among quantitative traits in the course of sympatric speciation. Nature 400, 351–354.

5. Gavrilets S. 2014 Models of speciation: where are we now? J. Hered. 105, 743–755. (doi:10.1093/jhered/esu045)

6. Martin CH, Richards EJ. 2019 The Paradox behind the Pattern of Rapid Adaptive Radiation: How Can the Speciation Process Sustain Itself through an Early Burst? Annu. Rev. Ecol. Evol. Syst.

7. Arnold SJ, Pfrender ME, G. JA. 2001 The Apaptive Landscape as a conceptual bridge between mico- and macroevolution. Genetica 112–113, 9–32.

8. Svennson E, Calsbeek R. 2012 The adaptive landscape in evolutionary biology. Oxford University Press.

9. Martin CH, Wainwright PC. 2013 Multiple fitness peaks on the adaptive landscape drive adaptive radiation in the wild. Science 339, 208–11. (doi:10.1126/science.1227710)

10. Martin CH, Gould KJ. 2020 Surprising spatiotemporal stability of a multi-peak fitness landscape revealed by independent field experiments measuring hybrid fitness. Evol. Lett., 530–544. (doi:10.1002/evl3.195)

11. Calsbeek R, Irschick DJ. 2007 The quick and the dead: Correlational selection on morphology, performance, and habitat use in island lizards. Evolution (N. Y). 61, 2493– 2503. (doi:10.1111/j.1558-5646.2007.00206.x)

12. Grant PR, Grant RB. 2008 How and Why Species Multiply. Princeton University Press.

13. Givnish TJ, Sytsma KJ, Smith J, Hahn W, DH B, Burkhardt E. 1997 Molecular evolution and adaptive radiation in *Brocchinia* (Bromeliaceae: Pitcairnioideae) atop tepuis of the Guyana Shield. In Molecular evolution and adaptive radiation, pp. 259–311. Cambridge: Cambridge University Press.

14. Givnish TJ, Burkhardt EL, Happel RE, Weintraub JD. 1984 Carnivory in the Bromeliad Brocchinia reducta, with a Cost / Benefit Model for the General Restriction of Carnivorous Plants to Sunny, Moist , Nutrient-Poor Habitats. Am. Nat. 124, 479–497.

15. Erwin DH. 2021 A conceptual framework of evolutionary novelty and innovation. Biol. Rev. 96, 1–15. (doi:10.1111/brv.12643)

16. Blount ZD, Barrick JE, Davidson CJ, Lenski RE. 2012 Genomic analysis of a key innovation in an experimental *Escherichia coli* population. Nature 489, 513–518. (doi:10.1038/nature11514)

17. Quandt EM, Deatherage DE, Ellington AD, Georgiou G, Barrick JE. 2014 Recursive genomewide recombination and sequencing reveals a key refinement step in the evolution of a metabolic innovation in Escherichia coli. Proc. Natl. Acad. Sci. U. S. A. 111, 2217– 2222. (doi:10.1073/pnas.1314561111)

18. Seehausen O. 2004 Hybridization and adaptive radiation. Trends Ecol. Evol. 19, 198–207. (doi:10.1016/j.tree.2004.01.003)

19. John MES, Dixon KE, Martin CH. 2020 Oral shelling within an adaptive radiation of pupfishes : Testing the adaptive function of a novel nasal protrusion and behavioural preference., 1–9. (doi:10.1111/jfb.14344)

20. Martin CH, Wainwright PC. 2013 A remarkable species flock of *Cyprinodon* pupfishes endemic to San Salvador Island, Bahamas. Bull. Peabody Museum Nat. Hist. 54, 231–240.

21. Turner BJ, Duvernell DD, Bunt TM, Barton MG. 2008 Reproductive isolation among endemic pupfishes (Cyprinodon) on San Salvador Island, Bahamas: Microsatellite evidence. Biol. J. Linn. Soc. 95, 566–582. (doi:10.1111/j.1095-8312.2008.01079.x)

22. Clark PU, Dyke AS, Shakun JD, Carlson AE, Clark J, Wohlfarth B, Mitrovica JX, Hostetler SW, McCabe AM. 2009 The Last Glacial Maximum. Science (80-.). 325, 710– 714. (doi:10.1126/science.1172873)

23. Richards EJ, Martin CH. 2017 Adaptive introgression from distant Caribbean islands contributed to the diversification of a microendemic adaptive radiation of trophic specialist pupfishes. PLoS Genet. 13, 1–35. (doi:10.1371/journal.pgen.1006919)

24. Jackson AL, Inger R, Parnell AC, Bearhop S. 2011 Comparing isotopic niche widths among and within communities: SIBER - Stable Isotope Bayesian Ellipses in R. J. Anim. Ecol. 80, 595–602. (doi:10.1111/j.1365-2656.2011.01806.x)

25. Richards EJ, McGirr JA, Wang JR, St. John ME, Poelstra JW, Solano MJ, O’Connell DC, Turner BJ, Martin CH. 2021 A vertebrate adaptive radiation is assembled from an ancient and disjunct spatiotemporal landscape. Proc. Natl. Acad. Sci. U. S. A. 118. (doi:10.1073/pnas.2011811118)

26. Peter BM. 2016 Admixture, population structure, and f-statistics. Genetics 202, 1485– 1501. (doi:10.1534/genetics.115.183913)

27. Excoffier L, Dupanloup I, Huerta-Sánchez E, Sousa VC, Foll M. 2013 Robust Demographic Inference from Genomic and SNP Data. PLOS Genet. 9, e1003905. (doi:10.1371/journal.pgen.1003905)

28. Pavlidis P, Živković D, Stamatakis A, Alachiotis N. 2013 SweeD: Likelihood-based detection of selective sweeps in thousands of genomes. Mol. Biol. Evol. 30, 2224–2234. (doi:10.1093/molbev/mst112)

29. Danecek P et al. 2011 The variant call format and VCFtools. Bioinformatics 27, 2156– 2158. (doi:10.1093/bioinformatics/btr330)

30. Smith J, Coop G, Stephens M, Novembre J. 2018 Estimating Time to the Common Ancestor for a Beneficial Allele. Mol. Biol. Evol. 35, 1003–1017. (doi:10.1101/071241)

31. Roesti M, Moser D, Berner D. 2013 Recombination in the threespine stickleback genome - Patterns and consequences. Mol. Ecol. 22, 3014–3027. (doi:10.1111/mec.12322)

32. St. John ME, Holzman R, Martin CH. 2020 Rapid adaptive evolution of scale-eating kinematics to a novel ecological niche. J. Exp. Biol. 223, jeb217570. (doi:10.1242/jeb.217570)

33. Martin CH, Wainwright PC. 2013 On the measurement of ecological novelty: scale-eating pupfish are separated by 168 my from other scale-eating fishes. PLoS One 8, e71164. (doi:10.1371/journal.pone.0071164)

34. Richards EJ, Martin CH. 2017 Adaptive introgression from distant Caribbean islands contributed to the diversification of a microendemic adaptive radiation of trophic specialist pupfishes. PLOS Genet. 13, e1006919.

35. Dickinson ME et al. 2016 High-throughput discovery of novel developmental phenotypes. Nature 537, 508–514. (doi:10.1038/nature19356)

36. Kida YS, Sato T, Miyasaka KY, Suto A, Ogura T. 2007 Daam1 regulates the endocytosis of EphB during the convergent extension of the zebrafish notochord. Proc. Natl. Acad. Sci. U. S. A. 104, 6708–6713. (doi:10.1073/pnas.0608946104)

37. St. John, Michelle E., Dunker JC, Richards, Emilie J., Romero S, Martin CH. 2021 Parallel genetic changes underlie integrated craniofacial traits in an adaptive radiation of trophic specialist pupfishes., 1–46.

38. Thomas T, Voss AK, Chowdhury K, Gruss P. 2000 Querkopf, a MYST family histone acetyltransferase, is required for normal cerebral cortex development. Development 127, 2537–2548.

39. Mcgirr JA, Martin CH. 2018 Parallel evolution of gene expression between trophic specialists despite divergent genotypes and morphologies. Evol. Lett. 2, 62–75. (doi:10.1002/evl3.41)

40. Whittaker R. 1977 Evolution of species diversity in land communities. Evol. Biol.

41. Stroud JT, Losos JB. 2016 Ecological Opportunity and Adaptive Radiation. Annu. Rev. Ecol. Evol. Syst. 47, 507–532. (doi:10.1146/annurev-ecolsys-121415-032254)

42. Kim BY, Huber CD, Lohmueller KE. 2018 Deleterious variation shapes the genomic landscape of introgression. PLoS Genet. 14, 1–30. (doi:10.1371/journal.pgen.1007741)

43. Zhang X, Kim B, Lohmueller KE, Huerta-Sánchez E. 2020 The impact of recessive deleterious variation on signals of adaptive introgression in human populations. Genetics 215, 799–812. (doi:10.1534/genetics.120.303081)

44. Martin CH, Wainwright PC. 2011 Trophic novelty is linked to exceptional rates of morphological diversification in two adaptive radiations of cyprinodon pupfish. Evolution (N. Y). 65, 2197–2212. (doi:10.1111/j.1558-5646.2011.01294.x)

45. Orr HA. 2005 The genetic theory of adaptation: A brief history. Nat. Rev. Genet. 6, 119– 127. (doi:10.1038/nrg1523)

46. Patton AH, Richards EJ, Gould KJ, Buie LK, Christopher H. 2021 Adaptive introgression and de novo mutations increase access to novel fitness peaks on the fitness landscape during a vertebrate adaptive radiation.

47. Weinreich DM, Delaney NF, DePristo MA, Hartl DL. 2006 Darwinian Evolution Can Follow Only Very Few Mutational Paths to Fitter Proteins. Science (80-.). 312, 111 LP – 114. (doi:10.1126/science.1123539)

48. Otte KA, Nolte V, Mallard F, Schlötterer C. 2021 The genetic architecture of temperature adaptation is shaped by population ancestry and not by selection regime. Genome Biol. 22, 1–25. (doi:10.1186/s13059-021-02425-9)

49. Hansen TF. 2013 Why epistasis is important for selection and adaptation. Evolution (N. Y). 67, 3501–3511. (doi:10.1111/evo.12214)

50. Fragata I, Blanckaert A, Dias Louro MA, Liberles DA, Bank C. 2019 Evolution in the light of fitness landscape theory. Trends Ecol. Evol. 34, 69–82. (doi:https://doi.org/10.1016/j.tree.2018.10.009)

## References

1. Wainwright PC, Bellwood DR, Westneat MW, Grubich JR, Hoey AS. 2004 A functional morphospace for the skull of labrid fishes: Patterns of diversity in a complex biomechanical system. Biol. J. Linn. Soc. 82, 1–25. (doi:10.1111/j.1095-8312.2004.00313.x)

2. Patricia Hernandez L, Gibb AC, Ferry-Graham L. 2009 Trophic apparatus in cyprinodontiform fishes: Functional specializations for picking and scraping behaviors. J. Morphol. 270, 645–661. (doi:10.1002/jmor.10711)

3. Martin CH, Wainwright PC. 2013 Multiple fitness peaks on the adaptive landscape drive adaptive radiation in the wild. Science 339, 208–11. (doi:10.1126/science.1227710)

4. Jackson AL, Inger R, Parnell AC, Bearhop S. 2011 Comparing isotopic niche widths among and within communities: SIBER - Stable Isotope Bayesian Ellipses in R. J. Anim. Ecol. 80, 595–602. (doi:10.1111/j.1365-2656.2011.01806.x)

5. Richards E, McGirr J, Wang J, St. John M, Poelstra J, Solano M, O’Connell D, Turner B, Martin C. 2020 Major stages of vertebrate adaptive radiation are assembled from a disparate spatiotemporal landscape. (doi:10.1101/2020.03.12.988774)

6. Li H, Durbin R. 2011 Inference of human population history from individual whole-genome sequences. Nature 475, 493–496. (doi:10.1038/nature10231)

7. DePristo MA et al. 2011 A framework for variation discovery and genotyping using next-generation DNA sequencing data. Nat. Genet. 43, 491–8. (doi:10.1038/ng.806)

8. Marsden CD, Lee Y, Kreppel K, Weakley A, Cornel A, Ferguson HM, Eskin E, Lanzaro GC. 2014 Diversity, differentiation, and linkage disequilibrium: prospects for association mapping in the malaria vector Anopheles arabiensis. G3 (Bethesda). 4, 121–31. (doi:10.1534/g3.113.008326)

9. Danecek P et al. 2011 The variant call format and VCFtools. Bioinformatics 27, 2156– 2158. (doi:10.1093/bioinformatics/btr330)

10. Purcell S et al. 2007 PLINK: A Tool Set for Whole-Genome Association and Population-Based Linkage Analyses. Am. J. Hum. Genet. 81, 559–575. (doi:10.1086/519795)

11. Alexander DH, Novembre J, Lange K. 2009 Fast model-based estimation of ancestry in unrelated individuals. Genome Res. 19, 1655–1664.

12. Excoffier L, Dupanloup I, Huerta-Sánchez E, Sousa VC, Foll M. 2013 Robust Demographic Inference from Genomic and SNP Data. PLOS Genet. 9, e1003905. (doi:10.1371/journal.pgen.1003905)

13. Schiffels S, Durbin R. 2014 Inferring human population size and separation history from multiple genome sequences. Nat. Genet. 46, 919. (doi:10.1038/ng.3015)

14. Nadachowska-Brzyska K, Burri R, Smeds L, Ellegren H. 2016 PSMC analysis of effective population sizes in molecular ecology and its application to black-and-white Ficedula flycatchers. Mol. Ecol. 25, 1058–1072. (doi:10.1111/mec.13540)

15. Martin CH, Crawford JE, Turner BJ, Simons LH. 2016 Diabolical survival in Death Valley: recent pupfish colonization, gene flow and genetic assimilation in the smallest species range on earth. Proc. R. Soc. B Biol. Sci. 283, 20152334. (doi:10.1098/rspb.2015.2334)

16. Pavlidis P, Živković D, Stamatakis A, Alachiotis N. 2013 SweeD: Likelihood-based detection of selective sweeps in thousands of genomes. Mol. Biol. Evol. 30, 2224–2234. (doi:10.1093/molbev/mst112)

17. Garrigan D, Geneva A. 2014 msmove: A modified version of Hudson’s coalescent simulator ms allowing for finer control and tracking of migrant genealogies. (doi:10.6084/m9.figshare.1060474)

18. Balsa-Canto E, Henriques D, Gabor A, Banga JR. 2016 AMIGO2, a toolbox for dynamic modeling, optimization and control in systems biology. Bioinformatics 32, 1–2. (doi:10.1093/bioinformatics/btw411)

19. Ge SX, Jung D. 2018 ShinyGO: a graphical enrichment tool for animals and plants. bioRxiv, 2. (doi:10.1101/315150)

20. Smith J, Coop G, Stephens M, Novembre J. 2018 Estimating Time to the Common Ancestor for a Beneficial Allele. Mol. Biol. Evol. 35, 1003–1017. (doi:10.1101/071241)

21. Bradbury PJ, Zhang Z, Kroon DE, Casstevens TM, Ramdoss Y, Buckler ES. 2007 TASSEL: Software for association mapping of complex traits in diverse samples. Bioinformatics 23, 2633–2635. (doi:10.1093/bioinformatics/btm308)

22. Roesti M, Moser D, Berner D. 2013 Recombination in the threespine stickleback genome - Patterns and consequences. Mol. Ecol. 22, 3014–3027. (doi:10.1111/mec.12322)

23. Feder JL, Xie X, Rull J, Velez S, Forbes A, Leung B, Dambroski H, Filchak KE, Aluja M. 2005 Mayr, Dobzhansky, and Bush and the complexities of sympatric speciation in Rhagoletis. Proc. Natl. Acad. Sci. U. S. A. 102 **Suppl**, 6573–80. (doi:10.1073/pnas.0502099102)

24. Pfeifer B, Wittelsbürger U, Ramos-Onsins SE, Lercher MJ. 2014 PopGenome: An efficient swiss army knife for population genomic analyses in R. Mol. Biol. Evol. 31, 1929–1936. (doi:10.1093/molbev/msu136)

26. Pfeifer B, Kapan DD. 2019 Estimates of introgression as a function of pairwise distances. BMC Bioinformatics 20, 1–11. (doi:10.1186/s12859-019-2747-z)

27. Martin SH, Davey JW, Jiggins CD. 2015 Evaluating the use of ABBA-BABA statistics to locate introgressed loci. Mol. Biol. Evol. 32, 244–257. (doi:10.1093/molbev/msu269)

28. Echelle AA, Carson EW, Echelle AF, Bussche D, Dowling TE. 2005 Historical Biogeography of the New-World Pupfish Genus Cyprinodon (Teleostei: Cyprinodontidae). Copeia 2005, 320–339.

29. Patterson N, Moorjani P, Luo Y, Mallick S, Rohland N, Zhan Y, Genschoreck T, Webster T, Reich D. 2012 Ancient admixture in human history. Genetics 192, 1069–1093. (doi:10.1534/genetics.112.145037)

30. Edwards CD. 2001 Effect of Salinity on the Ecology of Molluscs in the Inland Saline Waters of San Salvador Island: A Experiment in Progress. Proc. Eight Symp. Nat. Hist. Bahamas., 15–26.

31. Turner BJ, Duvernell DD, Bunt TM, Barton MG. 2008 Reproductive isolation among endemic pupfishes (Cyprinodon) on San Salvador Island, Bahamas: Microsatellite evidence. Biol. J. Linn. Soc. 95, 566–582. (doi:10.1111/j.1095-8312.2008.01079.x)

32. Clark PU, Dyke AS, Shakun JD, Carlson AE, Clark J, Wohlfarth B, Mitrovica JX, Hostetler SW, McCabe AM. 2009 The Last Glacial Maximum. Science (80-.). 325, 710–714. (doi:10.1126/science.1172873)

33. Dickinson ME et al. 2016 High-throughput discovery of novel developmental phenotypes. Nature 537, 508–514. (doi:10.1038/nature19356)

34. Kida YS, Sato T, Miyasaka KY, Suto A, Ogura T. 2007 Daam1 regulates the endocytosis of EphB during the convergent extension of the zebrafish notochord. Proc. Natl. Acad. Sci. U. S. A. 104, 6708–6713. (doi:10.1073/pnas.0608946104)

35. Lee JY, Seo D, You J, Chung S, Park JS, Lee JH, Jung SM, Lee YS, Park SH. 2017 The deubiquitinating enzyme, ubiquitin-specific peptidase 50, regulates inflammasome activation by targeting the ASC adaptor protein. FEBS Lett. 591, 479–490. (doi:10.1002/1873-3468.12558)

36. Heras J, Martin C. 2021 Nonadaptive radiation of the gut microbiome in an adaptive radiation of Cyprinodon pupfishes with minor shifts for scale-eating. bioRxiv

37. Soupene E. 2008 ATP8A1 activity and phosphatidylserine transbilayer movement. J. Receptor. Ligand Channel Res. Volume 1, 1–10. (doi:10.2147/jrlcr.s3773)

39. Kraft M et al. 2011 Disruption of the histone acetyltransferase MYST4 leads to a noonan syndrome-like phenotype and hyperactivated MAPK signaling in humans and mice. J. Clin. Invest. 121, 3479–3491. (doi:10.1172/JCI43428)

40. Clayton-Smith J et al. 2011 Whole-exome-sequencing identifies mutations in histone acetyltransferase gene KAT6B in individuals with the say-barber-biesecker variant of Ohdo syndrome. Am. J. Hum. Genet. 89, 675–681. (doi:10.1016/j.ajhg.2011.10.008)

41. Martin CH, Richards EJ. 2019 The Paradox behind the Pattern of Rapid Adaptive Radiation: How Can the Speciation Process Sustain Itself through an Early Burst? Annu. Rev. Ecol. Evol. Syst.

42. Whittaker R. 1977 Evolution of species diversity in land communities. Evol. Biol.

43. Stroud JT, Losos JB. 2016 Ecological Opportunity and Adaptive Radiation. Annu. Rev. Ecol. Evol. Syst. 47, 507–532. (doi:10.1146/annurev-ecolsys-121415-032254)

44. Orr HA. 1998 The Population Genetics of Adaptation: The Distribution of Factors Fixed during Adaptive Evolution. Evolution (N. Y). 52, 935. (doi:10.2307/2411226)

46. Gillespie JH. 1984 Molecular Evolution Over the Mutational Landscape. Evolution (N. Y). 38, 1116–1129. (doi:10.2307/2408444)

47. Kauffman S, Levin S. 1987 Towards a general theory of adaptive walks on rugged landscapes. J. Theor. Biol. 128, 11–45. (doi:https://doi.org/10.1016/S0022-5193(87)80029-2)

48. Maynard Smith J. 1970 Natural Selection and the Concept of a Protein Space. Nature 225, 563–564. (doi:10.1038/225563a0)

49. Bell G. 2013 Evolutionary rescue and the limits of adaptation. Philos. Trans. R. Soc. B Biol. Sci. 368, 20120080. (doi:10.1098/rstb.2012.0080)

50. Lindsey HA, Gallie J, Taylor S, Kerr B. 2013 Evolutionary rescue from extinction is contingent on a lower rate of environmental change. Nature 494, 463–467. (doi:10.1038/nature11879)

51. Rousselle M, Simion P, Tilak M-K, Figuet E, Nabholz B, Galtier N. 2020 Is adaptation limited by mutation? A timescale-dependent effect of genetic diversity on the adaptive substitution rate in animals. PLOS Genet. 16, e1008668.

52. Mani GS, Clarke BC. 1990 Mutational order: a major stochastic process in evolution. Proc. R. Soc. London. B. Biol. Sci. 240, 29–37. (doi:10.1098/rspb.1990.0025)

53. Schluter D. 2009 Evidence for Ecological Speciation and Its Alternative. Science (80-.). 323, 737 LP – 741. (doi:10.1126/science.1160006)

54. Good BH, McDonald MJ, Barrick JE, Lenski RE, Desai MM. 2017 The dynamics of molecular evolution over 60,000 generations. Nature 551, 45–50. (doi:10.1038/nature24287)

55. Schluter D, Conte GL. 2009 Genetics and ecological speciation. Proc. Natl. Acad. Sci. 106, 9955–9962. (doi:10.1073/pnas.0901264106)

56. Patwa Z, Wahl LM. 2008 The fixation probability of beneficial mutations. J. R. Soc. Interface 5, 1279–1289. (doi:10.1098/rsif.2008.0248)

57. Pavlidis P, Alachiotis N. 2017 A survey of methods and tools to detect recent and strong positive selection. *J*. Biol. Res. 24, 7. (doi:10.1186/s40709-017-0064-0)

58. Höllinger I, Pennings PS, Hermisson J. 2019 Polygenic adaptation: From sweeps to subtle frequency shifts. PLOS Genet. 15, e1008035.

59. Barghi N, Hermisson J, Schlötterer C. 2020 Polygenic adaptation: a unifying framework to understand positive selection. Nat. Rev. Genet. 21, 769–781. (doi:10.1038/s41576-020-0250-z)

60. Chevin LM, Hospital F. 2008 Selective sweep at a quantitative trait locus in the presence of background genetic variation. Genetics 180, 1645–1660. (doi:10.1534/genetics.108.093351)

61. Thornton KR. 2019 Polygenic adaptation to an environmental shift: Temporal dynamics of variation under Gaussian stabilizing selection and additive effects on a single trait. Genetics 213, 1513–1530. (doi:10.1534/genetics.119.302662)

62. Flint HJ, Scott KP, Louis P, Duncan SH. 2012 The role of the gut microbiota in nutrition and health. Nat. Rev. Gastroenterol. Hepatol. 9, 577–589. (doi:10.1038/nrgastro.2012.156)

63. Mcgirr JA, Martin CH. 2018 Parallel evolution of gene expression between trophic specialists despite divergent genotypes and morphologies. Evol. Lett. 2, 62–75. (doi:10.1002/evl3.41)

64. Martin CH, Gould KJ. 2020 Surprising spatiotemporal stability of a multi-peak fitness landscape revealed by independent field experiments measuring hybrid fitness. Evol. Lett., 530–544. (doi:10.1002/evl3.195)

